# Mosquito fertility is (mostly) resilient to sublethal heat shocks

**DOI:** 10.64898/2026.08.04.742888

**Authors:** Apeksha L. Warusawithana, Belinda van Heerwaarden, Ary A. Hoffmann, Perran A. Ross

## Abstract

Insects can experience a loss of fertility under acute heat stress, but impacts vary among species and depend on life stage and sex. We examined the effects of sublethal heat shocks on the fertility of three mosquito species (*Aedes aegypti*, *Aedes notoscriptus* and *Culex quinquefasciatus*) when males or females were exposed at the pupal and adult stages. Reproductive performance was assessed through egg-laying success, fecundity, egg hatchability and viable offspring production. Female heat shock had limited effects on fertility in all three species, though exposure of adult females to 41 °C for 1 hr reduced viable offspring production in *Ae. notoscriptus*. Male heat shock reduced egg hatchability in *Cx*. *quinquefasciatus* and viable offspring production in *Ae. aegypti* depending on the life stage exposed. We investigated several other aspects of heat shock exposure on *Ae. aegypti* fertility. In this species, male fertility partially recovered within 24-48 hours following heat shock exposure, while effects of heat shocks on female fertility occurred regardless of whether heat exposure occurred pre- or post-mating. In addition, maternal heat shock exposure enhanced offspring fecundity and viable offspring production under subsequent sublethal heat stress. Overall, our findings demonstrate that mosquito fertility is largely robust to short-term heat shocks, but there are also complex trait, sex, stage, temporal and species-specific effects, with implications for predicting mosquito population persistence and species persistence under future climate extremes.

## Introduction

Mosquito fertility is a key determinant of population persistence and a critical contributor to vectorial capacity (Brady et al., 2016; Mitchell & Catteruccia, 2017). High fecundity and successful egg hatchability facilitate rapid population growth (Clements, 2023), amplifying the potential for mosquito-borne pathogen transmission (Mitchell & Catteruccia, 2017). Consequently, environmental factors affecting mosquito reproductive success, particularly fertility, directly influence vectorial capacity and the likelihood of disease outbreaks (Chandrasegaran et al., 2020; Mordecai et al., 2019).

As ectotherms, mosquitoes are sensitive to temperature, with reproductive performance peaking within a species-specific thermal optimum and declining at both lower and upper thermal extremes (Carrington et al., 2013; Mordecai et al., 2019). Anthropogenic climate change has increased mean environmental temperatures and the frequency and intensity of extreme heat events (King et al., 2016), such that environmental conditions in many regions, particularly the tropics, now approach or exceed the upper thermal limits of insects (García-Robledo & Baer, 2021; Holzmann et al., 2026). Evidence from *Drosophila* (Parratt et al., 2021; van Heerwaarden & Sgro, 2021) and other insects (Weaving et al., 2024) indicates that exposure to temperatures below the critical thermal maximum for survival can substantially impair fertility. These findings underscore the importance of examining thermal effects on mosquito fertility rather than survival alone.

Previous studies have shown that chronic exposure to elevated temperatures disrupts multiple aspects of mosquito reproductive biology, including reductions in testicular volume, sperm count, and sperm velocity (Padde et al., 2024), ovariole count (Bader & Williams, 2013), egg count (Delatte et al., 2009; Ezeakacha & Yee, 2019; Padde et al., 2024; Pekľanská et al., 2025), egg hatchability (Padde et al., 2024), and shortening of the gonotrophic cycle (Delatte et al., 2009; Rúa et al., 2005). These reproductive impairments are further exacerbated when immature stages are reared at elevated temperatures (Agyekum et al., 2022; Padde et al., 2024; Pekľanská et al., 2025). Elevated temperatures also alter key reproductive behaviours, including mating activity (Schoof, 1967), host-seeking and blood-feeding propensity (Costanzo & Occhino, 2023), blood feeding frequency (Scott et al., 2000), and biting rate (Agyekum et al., 2022).

Although the effects of constant elevated temperatures on mosquito reproduction are well documented (Agyekum et al., 2022; Delatte et al., 2009; Hernández et al., 2026; Pekľanská et al., 2025), the effects of acute sublethal heat stress on mosquito fertility remain poorly understood. Studies in *Drosophila* show that sublethal heat shock can impair fertility in a stage-specific and sex-specific manner (Meena et al., 2025; Walsh et al., 2021), and that effects on male fertility can be delayed or partially reversible following stress depending on exposure intensity and duration (Canal Domenech & Fricke, 2022; Meena et al., 2024), . To our knowledge, only Mourya et al. (2004) have partially addressed this question in mosquitoes, reporting reduced fecundity in *Aedes aegypti* following repeated sublethal acute heat stress (44.5 °C for 10 min) during larval development across multiple generations. Brief but intense temperature spikes experienced during the hottest part of the day are becoming increasingly frequent under current climate change (Buckley & Huey, 2016; Calvin et al., 2023). Evaluating the effects of sublethal acute heat stress on mosquito fertility is therefore important, particularly during the pupal stage, when spermatogenesis and early oogenesis occur, and during adulthood, when vitellogenesis, fertilisation and embryogenesis take place (Clements, 2023; Wang & Li, 2024). However, there is limited information on sublethal acute heat stress applied to immature or adult stages on subsequent reproductive output, and it is also not known if there are transgenerational effects of parental acute heat stress on offspring fertility, even though these effects are known in insects (Sales et al., 2018) and may form an important component of the broader acclimation response (Hoffmann & Bridle, 2022).

This study addresses these knowledge gaps by examining acute heat shock effects on the reproductive biology of three epidemiologically important mosquito vectors: *Aedes aegypti*, the principal dengue vector (Brady & Hay, 2020; Jansen & Beebe, 2010); *Aedes notoscriptus*, a vector of Ross River virus (Claflin & Webb, 2015) and *Mycobacterium ulcerans* (Buruli ulcer) in Australia (Mee et al., 2024); and *Culex quinquefasciatus*, a primary vector of lymphatic filariasis (Negi & Verma, 2018). Specifically, we investigate: (i) the effects of acute heat shocks applied separately to pupae and adults of both sexes on fertility across the three vector species; (ii) the temporal dynamics of heat stress on male fertility in *Ae. aegypti*; (iii) the influence of female mating status on fertility in *Ae. aegypti*; and (iv) the transgenerational effects of maternal heat stress on offspring fertility in *Ae. aegypti*.

## Methodology

### Mosquito rearing and colony maintenance

The three mosquito colonies representing *Aedes aegypti*, *Aedes notoscriptus* and *Culex quinquefasciatus* originated from eggs collected in Queensland, Australia were established and maintained under controlled standard insectary conditions (26 ± 0.5 °C, 50-70% relative humidity (RH) and 12:12 h light: dark cycle) for at least 50 generations.

Eggs (*Ae. aegypti* and *Ae. notoscriptus,* < 1 week old; *Cx. quinquefasciatus,* 1 day old) were hatched by adding 5-10 grains of active dry yeast to 500 mL of reverse osmosis (RO) water. Larval density was controlled to approximately 500 larvae per plastic tray (40 x 30 x 6.5 cm) containing 4 L of water (RO water for *Ae. aegypti* and *Ae. notoscriptus*; hay-infused (≥ 1 week fermented) RO water for *Cx. quinquefasciatus*)) to reduce density-dependent variation in immature development. Larval food was provided daily at 0.05 mg per larva (Hikari Tropical Sinking Wafers, Kyorin Food Co. Ltd., Himeji, Japan). Seven days post hatching, pupae were collected into 500 mL plastic cups with 250 mL larval rearing water and transferred to 27 L BugDorm-1 cages (Megaview Science Co. Ltd., Taichung, Taiwan) targeting approximately 500 adults per cage. Adults were maintained on 10% sucrose solution replaced fortnightly, and sucrose was substituted with water approximately 24 hours prior to blood feeding to enhance female feeding propensity. Female mosquitoes were blood fed on the forearm of a single adult human volunteer under ethics approval from the University of Melbourne Human Ethics Committee (Project ID 28583). Eggs were collected 4-5 days post blood feeding on a strip of sandpaper (20 cm x 4 cm, Norton Master Painters P80: Saint-Gobain Abrasives Pty. Ltd., Thomastown, Victoria, Australia) that lined a plastic cup (500 mL) containing 250 mL of larval rearing water for *Ae. aegypti* and *Ae. notoscriptus,* or directly on the water surface for *Cx. quinquefasciatus*.

### Individual egg laying for fertility assessment

Fully engorged females were randomly selected and assigned to meshed, individual plastic cups (125 mL for *Ae. aegypti* and *Cx. quinquefasciatus*; 500 mL for *Ae. notoscriptus*, as most females avoided laying eggs in 125 mL cups based on a pilot experiment), partially filled (25 mL for a 125 mL cup; 100 mL for a 500 mL cup) with RO water mixed with pre-larval rearing water (∼4:1 ratio) for egg laying. The lower part of the egg-laying cups was lined with a strip of sandpaper (20 cm x 4 cm, Norton Master Painters P80: Saint-Gobain Abrasives Pty. Ltd., Thomastown, Victoria, Australia) for the *Aedes* species. Egg papers were collected 4-5 days post blood feeding, wrapped in paper towels to maintain moisture, and stored in 500 mL plastic containers. Egg papers containing matured eggs (3-4 days old) were hatched by adding 2-3 grains of active yeast per cup for 24 hours. In contrast, *Cx. quinquefasciatus* egg rafts (1 day old) were left to hatch directly in the egg-laying cup. Fecundity and hatchability were quantified by counting the total number of eggs laid and the number of eggs hatched per female.

### Sublethal heat shock effects on female fertility (*Ae. aegypti, Ae. notoscriptus* and *Cx. quinquefasciatus*)

To determine the effect of sublethal heat shock at both pupal and adult stages on *Ae. aegypti* female fertility, two sets of female pupae (sex separation was based on body size and cephalothorax shape), each containing approximately 100 individuals in ∼5 mL of RO water, were transferred to ∼250 mL of preheated RO water in 500 mL plastic containers in two pre-programmed incubators (PHcbi MIR-254) at 42 °C and 43 °C, respectively. After a one-hour heat shock, pupae were immediately transferred to ∼250 mL RO water in 500 mL plastic containers maintained at 26 °C. Heat treated and untreated pupae were separated by sex and allowed to emerge separately in BugDorm cages under standard insectary conditions and fed with 10% sucrose solution.

For adult treatments, two sets of 2-3 days old unmated females, each with approximately 100 individuals in 1.5 L plastic cages without access to water, were exposed to one-hour heat shocks in pre-programmed incubators at 42 °C and 43 °C, separately. Following exposure, adults were transferred to 26 °C and allowed to recover for 24 hours with access to 10% sucrose solution. Females emerging from the two heat-treated pupal groups and the two heat-treated adult groups, along with untreated control females (100 individuals), were exposed to untreated males derived from same cohort and maintained at 26 °C for mating at a 1:1 sex ratio. Two days post-mating, female groups were blood fed and individually isolated for egg-laying to assess fertility.

The same experimental procedure was repeated for *Ae. notoscriptus* female pupae and adults exposed to sublethal temperatures of 40 °C and 41 °C, and for *Cx. quinquefasciatus* female pupae exposed to 41 °C and 42 °C, with adult females of this species exposed to 39 °C and 40 °C. These species-specific sublethal heat shock exposure temperatures were selected based on previously estimated critical thermal maxima (CTmax) (*Ae. aegypti*, 44.4 °C; *Ae. notoscriptus,* 42.1 °C; *Cx. quinquefasciatus,* 40.8 °C) (Warusawithana et al., 2026) and pilot experimental exposures. In pilot experiments, although adult *Cx. quinquefasciatus* died above their CTmax, pupae survived even at 42 °C, hence the different exposure temperatures between life stages. Fertility was assessed in 30 blood-fed females per treatment for *Aedes* species and 35 per treatment for *Culex quinquefasciatus* (Figure 1A).

**Figure 1.**
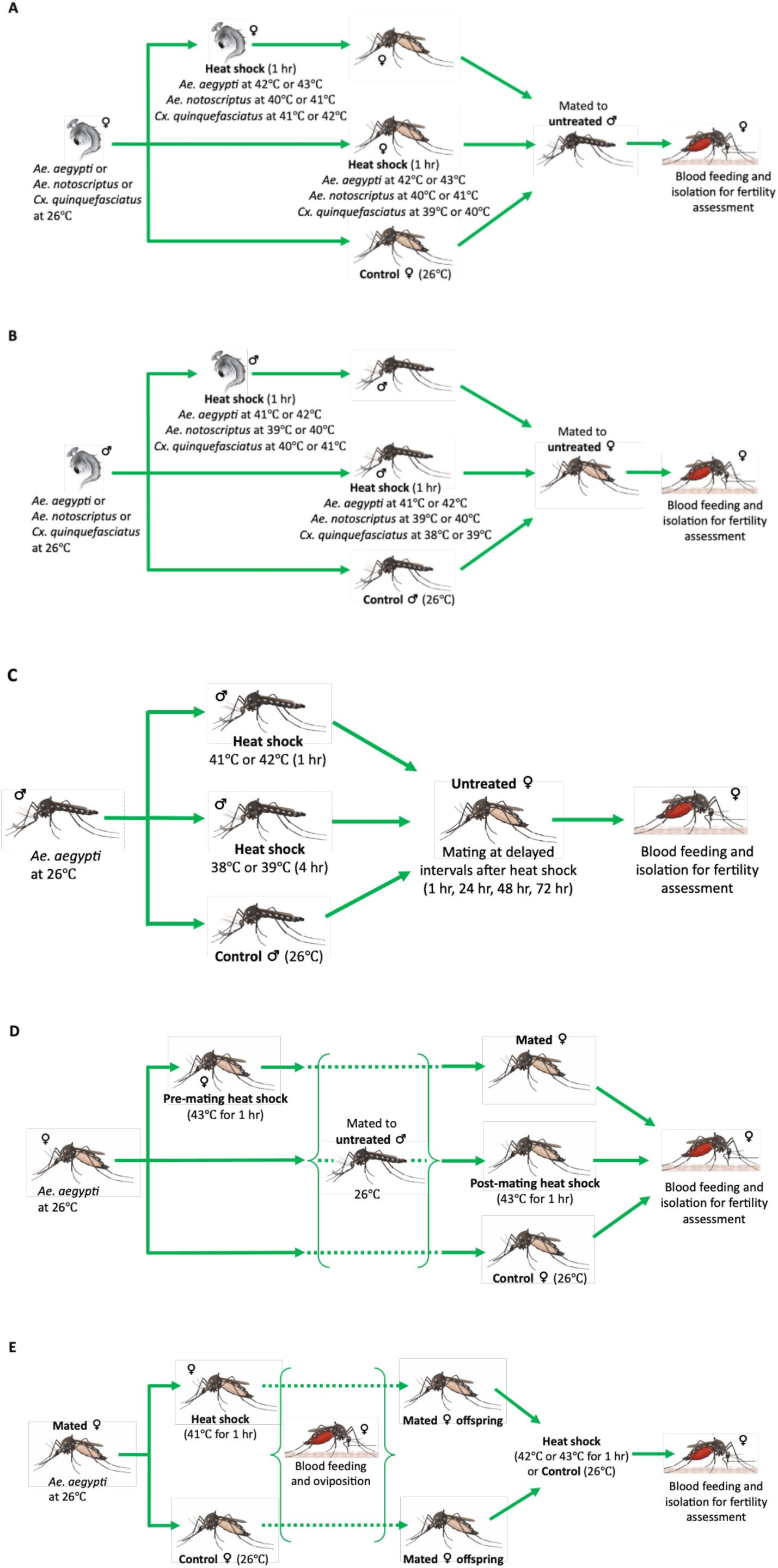
Experimental designs used to quantify the effect of sublethal heat stress on mosquito fertility across sexes, life stages and generations. (A) Female fertility assessment in *Ae. aegypti, Ae. notoscriptus* and *Cx. quinquefasciatus*: pupae and adult females were exposed to sublethal heat shocks, then mated with untreated males for fertility assessment. (B) Male fertility assessment in *Ae. aegypti, Ae. notoscriptus* and *Cx. quinquefasciatus*: pupae and adult males were exposed to sublethal heat shocks, then mated with untreated females for fertility assessment. (C) Delayed effects of sublethal heat shock in male *Ae. aegypti*: males were exposed to heat shocks and allowed a defined recovery period prior to mating to assess fertility. (D) Influence of mating status in female *Ae. aegypti*: females were heat-shocked either pre-mating or post-mating to assess the effects on fertility. (E) Transgenerational effects in *Ae. aegypti:* both mothers and offspring were exposed to sublethal heat shocks to assess transgenerational effects on fertility.

Sublethal heat shock effects on male fertility (*Ae. aegypti, Ae. notoscriptus* and *Cx. quinquefasciatus*)

To evaluate male fertility, the experimental procedure used for the female fertility experiment was followed, with male pupae and adults exposed to different temperature treatments. Male *Ae. aegypti* pupae and adults were exposed to one-hour heat shocks at 41 °C and 42 °C. Following a 24-hour recovery period at 26 °C, heat-treated adult males and males that had developed from heat-treated pupae were mated with untreated and unmated females (1:1 sex ratio) derived from the same cohort. Two days post-mating, females were blood-fed and individually assessed for fertility, together with untreated control groups. Similarly, *Ae. notoscriptus* male pupae and adults were exposed to 39 °C and 40 °C, while *Cx. quinquefasciatus* male pupae were exposed to 40 °C and 41 °C, and adult males to 38 °C and 39 °C (Figure 1B). Male exposure temperatures were set 1 °C lower than those used for females because males exhibited lower heat tolerance in heat survival assays (Warusawithana et al., 2026).

## Delayed effects of sublethal heat shock on *Ae. aegypti* male fertility

To evaluate the delayed effects of sublethal heat shock on *Ae. aegypti* male fertility, males were exposed to heat shocks and subsequently mated at different post-exposure time points. Specifically, 2-3 day old male mosquitoes (100 per treatment) were exposed either to an acute sublethal one-hour heat shock at 41 °C or 42 °C, consistent with the previous male heat shock experiment, or to a prolonged four-hour heat shock at 38 °C or 39 °C, as commonly used in *Drosophila* heat shock studies (Parratt et al., 2021; Walsh et al., 2021). At four post-exposure intervals (1 h, 24 h, 48 h, and 72 h), 50 males from each treated group were paired with same-aged unmated females reared at 26 °C at a 1:1 sex ratio for a 48-hour mating period. Similarly, 50 untreated control males were paired with untreated females (1:1 sex ratio) at two post-exposure intervals (1 h and 72 h). Following the 48-hour mating period, mated females from all groups were simultaneously blood-fed, and 35 females per group were set up for individual egg-laying to assess fertility (Figure 1C).

## Effect of mating status on heat shock induced changes in *Ae. aegypti* female fertility

To test whether female mating status influences the effect of heat shock on mosquito fertility and its potential effects on stored sperm, females were exposed to a heat shock before or after mating (pre-mating or post-mating). In the pre-mating treatment, 100 unmated *Ae. aegypti* females (2-3 days old) were exposed to a one-hour heat shock at 43 °C in a pre-programmed incubator without access to water. This group, together with two additional cohorts of 100 untreated unmated females (2-3 days old) per group, was allowed to mate with untreated males reared at 26 °C at a 1:1 sex ratio. Following mating, one cohort of untreated mated females was exposed to a one-hour heat shock at 43 °C without access to water (i.e., post-mating treatment). The following day, females from the pre-mating, post-mating, and untreated control groups were blood-fed. Fifty-five females per treatment group were individually isolated for egg laying to assess fertility. Additionally, approximately 40-45 surviving females per group were assessed for fertility during the second gonotrophic cycle (Figure 1D).

## Transgenerational effects of heat shock exposure in *Ae. aegypti* female fertility

To determine the transgenerational effects of heat shock exposure on *Ae. aegypti* fertility, 4-5 day-old mated female mosquitoes were exposed to a one-hour heat shock at 41 °C without access to water (i.e., heat-treated mothers). This temperature was selected because it was previously used as the heat-hardening treatment for *Ae. aegypti*, where it induced a sublethal physiological response without causing knockdown or mortality (Warusawithana et al., 2026). After exposure, females were provided with 10% sucrose solution and allowed to recover for 24 hours under controlled standard insectary conditions (26 ± 0.5 °C, 50-70% relative humidity (RH) and 12:12 h light: dark cycle). Both the heat-treated and untreated females were blood-fed to collect eggs for the next generation. Mated female offspring from both heat-treated and untreated mothers, reared under standard insectary conditions, were exposed to a one-hour heat shock at 42 °C and 43 °C, separately, without access to water. Two additional sets of offspring from heat-treated and untreated mothers were maintained as untreated controls. After 24 hours of recovery at 26 °C with access to 10% sucrose solution, followed by 24 hours of starvation, females from all six groups (offspring of heat-treated mothers: two heat-shocked and one untreated offspring group; offspring of untreated mothers: two heat-shocked and one untreated offspring group) were blood-fed, and 35 females per group were assessed for fertility (Figure 1E).

## Data analysis

Experiments with different species were conducted separately, and statistical analyses were therefore restricted by focussing on treatment comparisons within species. Fertility was assessed using four parameters: egg-laying success (proportion of females laying eggs), fecundity (total number of eggs per female), egg hatchability (proportion of eggs hatched per female), and viable offspring (total number of hatched eggs per female). Females that died before oviposition were excluded from all analyses to distinguish between heat-shock induced sterility and mortality. For the experiment on the delayed effect of heat shock on male fertility where control data were not collected across all sampling time points, the control group was excluded from statistical modelling to maintain a balanced experimental design; however, control data were retained in figures for qualitative comparison.

### Model specification

All statistical analyses and data visualisation were conducted in R using RStudio (version 2025.10.31+748; (RStudio Team, 2024). Data processing and visualisation were performed using the tidyverse packages (version 2.0.0; Wickham et al. (2019). Statistical models were fitted to match the distribution of each reproductive traits via generalised linear models (GLMs) using the brglm2 (version 1.0.1; Kosmidis and Firth (2021) and glmmTMB (version 1.1.14; McGillycuddy et al. (2025) packages. Model assumptions, including appropriate error distribution, overdispersion, and zero-inflation, were evaluated using residual diagnostics and visual inspection of residual and Q-Q plots in the DHARMa package (version 0.4.7; Hartig (2016) using 1,000 simulations. Factorial models included all main effects and biologically relevant two-way interactions.

### Egg laying success

The probability of a female that successfully oviposited (binary response) was modelled using a bias-reduced binomial GLM (logit link) via the brglm2 package (version 1.0.1; Kosmidis and Firth (2021) to provide stable parameter estimates in the presence of complete separation (e.g. 100% egg laying frequencies in certain treatments).

### Fecundity

For females that successfully oviposited (excluding those with zero eggs), total egg counts were analysed using a Conway-maxwell Poisson (COM-Poisson) GLM via the glmmTMB package (version 1.1.14; McGillycuddy et al. (2025). This distribution was selected to accommodate both under-dispersion and over-dispersion in the count data. The dispersion parameter was modelled as a function of treatment to account for heteroscedasticity.

### Egg hatchability

The proportion of eggs that hatched was analysed using a Zero-Inflated Beta-Binomial (ZIBB) GLM via the glmmTMB package (version 1.1.14; McGillycuddy et al. (2025) to account for both overdispersion and an excess of zeros representing complete hatching failure. The zero-inflation was modelled with a constant intercept to ensure numerical stability and model convergence across all experimental groups. Females that did not lay eggs or laid fewer than 5 eggs were excluded to ensure mathematically robust proportional data.

### Viable offspring

The total number of viable offspring (including zero counts) was analysed using a Truncated Negative Binomial Hurdle model using the glmmTMB package (version 1.1.14; McGillycuddy et al. (2025), which effectively addressed the heavy right skew and zero-inflation by partitioning the analysis into a binary component (probability of producing any viable offspring) and a count component (quantity of offspring produced). The zero-inflation was modelled with a constant intercept to ensure numerical stability and model convergence across all experimental groups.

### Post-hoc inference and visualisation

Statistical significance for main effect and interactions was assessed using Type II Wald chi-square tests (χ^2^) via the car package (version 3.1.3; Fox and Weisberg (2018). Post-hoc pairwise comparisons of estimated marginal means (EMMs) were conducted using the emmeans package (version 1.8.5; Lenth (2023) with Tukey’s HSD adjustments for multiple comparisons. Results are reported as odds ratios (for binomial and beta-binomial models) or response ratios (for count models), each with 95% confidence intervals (CIs), test statistics (*Z* or χ^2^), and *p-*values at the significance level of α = 0.05.

Figures were generated using the ggplot2 package (version 4.0.1; Wickham (2016). Egg-laying success was visualised using bar charts representing model-estimated proportions with 95% CIs. For fecundity, egg hatchability and viable offspring, data were visualised using box-and-whisker plots (median and interquartile range) overlaid with jittered raw data points. Model-estimated marginal means and 95% CIs were superimposed to facilitate comparison between observed data distributions and statistical inference.

## Results

### Sublethal heat shock effects on female mosquito fertility

We assessed the effects of sublethal heat shock exposure in females at the pupal and unmated adult stages on reproductive success in all three mosquito species. Treatment (defined as each combination of temperature and life stage of exposure) did not influence egg laying success (all p > 0.178; Figure 2A – C) and egg hatchability (all p > 0.093; Figure 2G – I) in any species. *Ae. notoscriptus* exhibited particularly high variability in egg hatchability across treatments compared to the other species (Figure 2H, Table S1). In contrast, fecundity was significantly affected by treatment in all three species: *Ae. aegypti* (χ^2^ (4) = 18.703, *p* < 0.001; Figure 2D), *Ae. notoscriptus* (χ^2^ (4) = 18.987, *p* < 0.001; Figure 2E) and *Cx. quinquefasciatus* (χ^2^ (4) = 14.225, *p* = 0.007; Figure 2F) (Table S1). The number of viable offspring was not significantly affected by treatment in *Ae. aegypti* (χ^2^ (4) = 8.873, *p* = 0.064; Figure 2J), whereas both *Ae. notoscriptus* (χ^2^ (4) = 14.904, *p* = 0.005; Figure 2K) and *Cx. quinquefasciatus* (χ^2^ (4) = 13.467, *p* = 0.009; Figure 2L) exhibited a significant effect of treatment on the number of viable offspring (Table S1).

**Figure 2.**
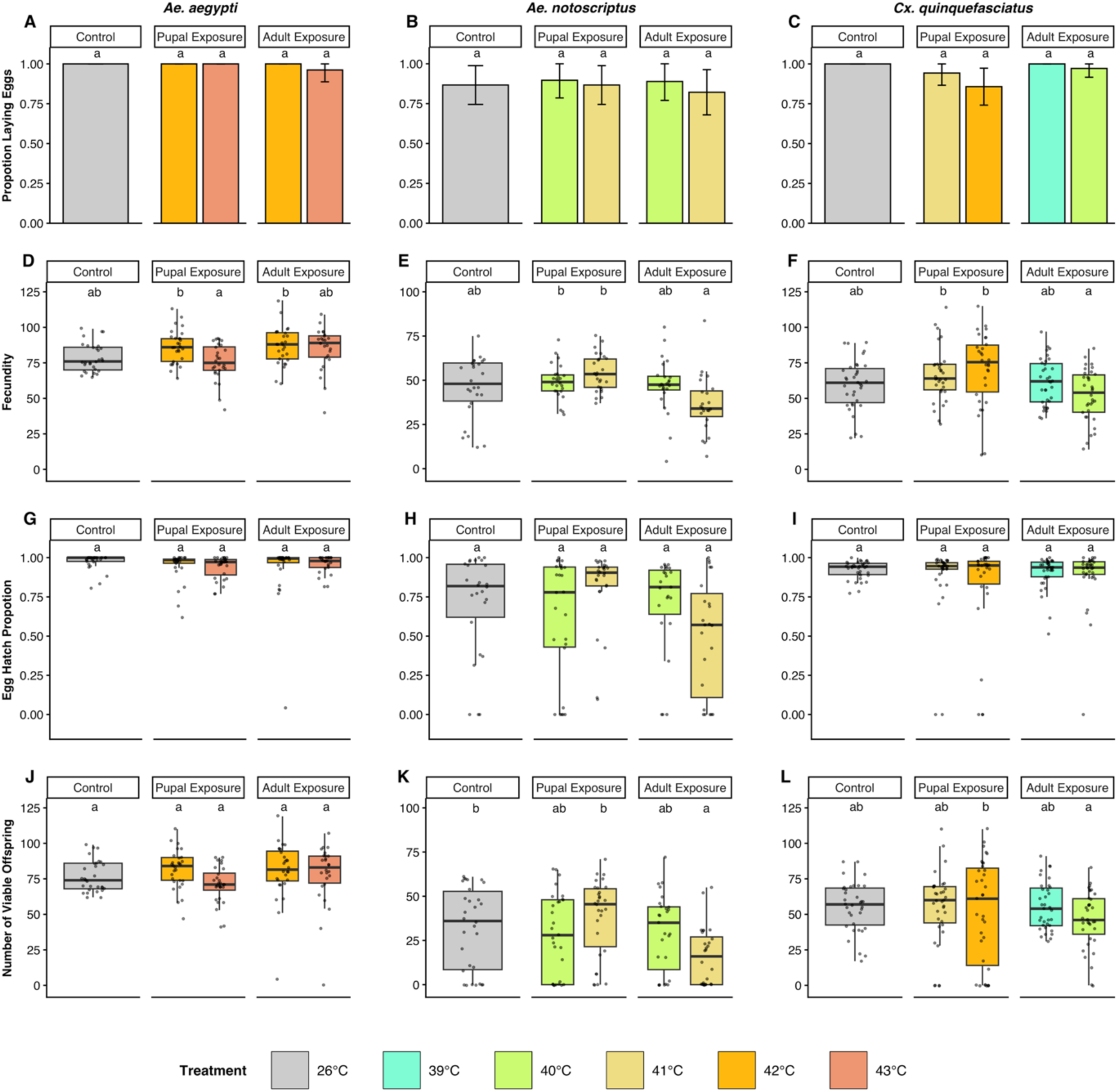
Effects of sublethal heat shock exposure during the pupal and adult stages on the reproductive success of female *Ae. aegypti* (A, D, G, J), *Ae. notoscriptus* (B, E, H, K), and *Cx. quinquefasciatus* (C, F, I, L). We measured the proportion of females laying eggs (A – C), fecundity (D – F), egg hatchability (G – I) and the number of viable offspring (J – L). Untreated controls are shown in grey, while sublethal heat shock treatments are shown in other colours. Bars in A – C represent mean proportions with error bars indicating 95% confidence intervals. Boxplots in D – L show medians and interquartile ranges; grey overlaid jittered dots represent individual observations. Different lowercase letters indicate significant differences; shared letters indicate no significant difference (*p* < 0.05).

In pairwise comparisons between treatments, no heat shock treatment differed significantly from the untreated control for any trait (all *p* > 0.051), except for *Ae. notoscriptus,* where exposure to 41 °C during the adult stage resulted in significantly fewer viable offspring compared to the control (26 °C vs 41 °C: Odds Ratio (OR) = 1.697, *z* = 3.171, *p* = 0.013) (Table S2). Pairwise comparisons among *Ae. notoscriptus* heat shock treatments revealed clear stage-specific differences at 41 °C, with both fecundity (pupa 41 °C vs adult 41 °C: OR = 1.489, *z* = 4.067, *p* = 0.001) and the number of viable offspring (pupa 41 °C vs adult 41 °C: OR = 1.797, *z* = 3.609, *p* = 0.003) being significantly lower following adult exposure compared to pupal exposure at the same temperature (Table S2). Overall, our results indicate limited effects of sublethal heat shocks at both the pupal and adult stages on female fertility across the three species tested.

## Sublethal heat shock effects on male mosquito fertility

We then assessed the effects of sublethal heat shock exposure in males at the pupal and adult stages on female reproductive success. In the male treatments (defined as each combination of temperature and life stage of exposure), female egg laying success (all p > 0.506; Figure 3A – C) and fecundity (all p > 0.069; Figure 3D – F) were not significantly affected by treatment in any species (Table S3). In contrast, egg hatchability was significantly affected in *Ae. aegypti* (χ^2^ (4) = 23.763, *p* < 0.001; Figure 3G) and *Cx. quinquefasciatus* (χ^2^ (4) = 12.044, *p* = 0.017, Figure 3I) (Table S3). In *Ae. aegypti,* pairwise comparisons revealed that egg hatchability in the control did not differ significantly from any heat shock treatment (all *p* > 0.066), but there was significantly lower hatchability following pupal heat exposure compared to adult exposure at 41 °C (pupa 41 °C vs adult 41 °C: OR = 0.227, *z* = -3.457, *p* = 0.005; Table S4). In *Cx quinquefasciatus*, egg hatchability was reduced relative to the control following adult stage heat exposure at 39 °C (26 °C vs 39 °C: OR = 2.015, *z* = 3.209, *p* = 0.012) (Table S4). The number of viable offspring was not significantly affected by treatment except for *Ae. aegypti* (χ^2^ (4) = 9.625, *p* = 0.047; Figure 3J; Table S3), with the strongest decline observed following pupal stage heat exposure compared to the control (26 °C vs 42 °C: OR = 1.199, *z* = 3.003, *p* = 0.023; Table S4). Although treatment effects on viable offspring production were not statistically significant in *Ae. notoscriptus* (*p* = 0.841; Table S3), the highest adult heat exposure resulted in a marked reduction, with a median of zero viable offspring (Figure 3K)

**Figure 3.**
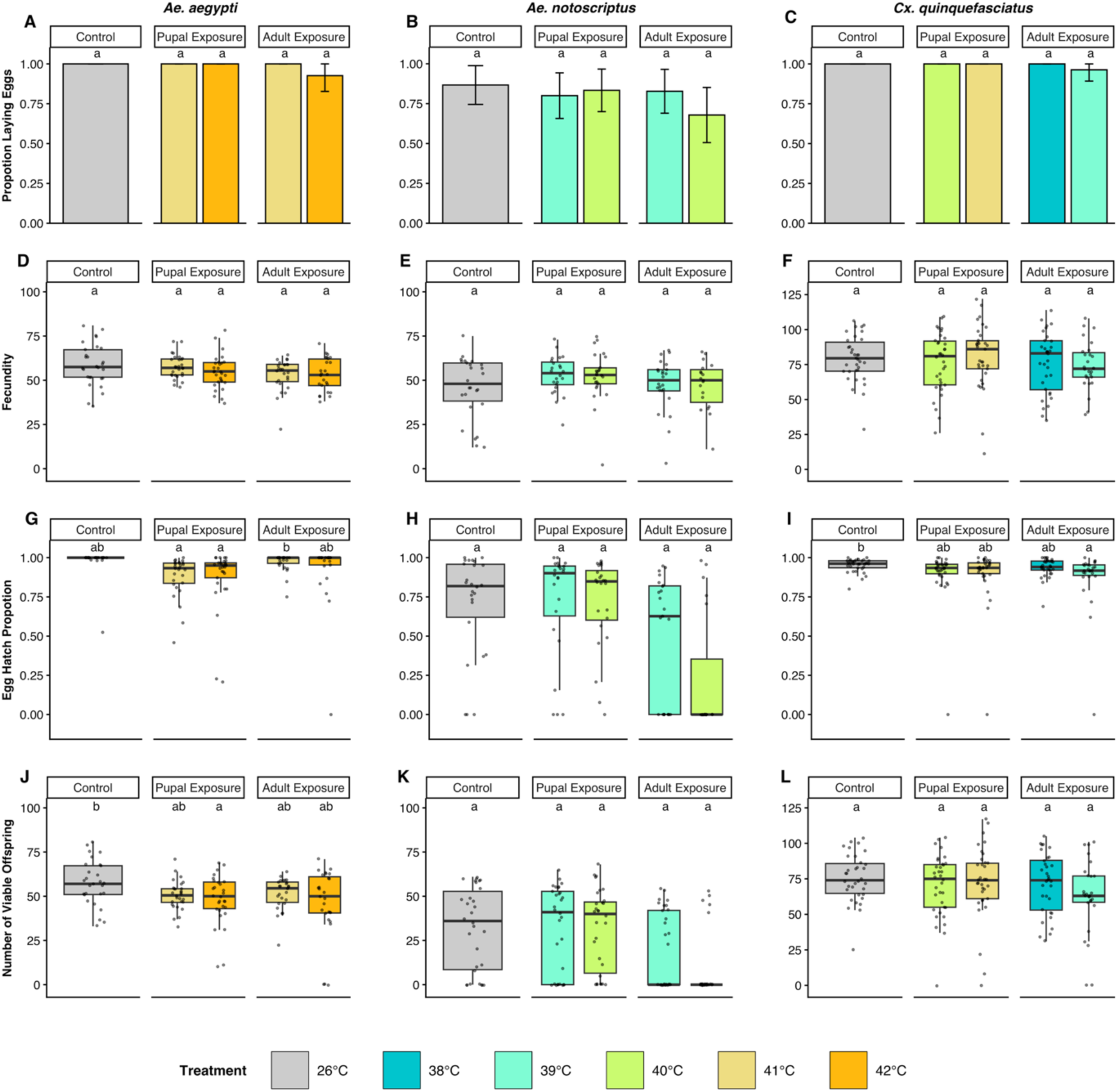
Effects of sublethal heat shock exposure during the pupal and adult stages on the reproductive success of male *Ae. aegypti* (A, D, G, J), *Ae. notoscriptus* (B, E, H, K), and *Cx. quinquefasciatus* (C, F, I, L). We measured the proportion of females laying eggs (A – C), fecundity (D – F), egg hatchability (G – I) and the number of viable offspring (J – L). Untreated controls are shown in grey, while sublethal heat shock treatments are shown in other colours. Bars in A – C represent mean proportions with error bars indicating 95% confidence intervals. Boxplots in D – L show medians and interquartile ranges; grey overlaid jittered dots represent individual observations. Different lowercase letters indicate significant differences; shared letters indicate no significant difference (*p* < 0.05).

## Delayed effects of sublethal heat shock on *Ae. aegypti* male fertility

Previous studies have identified delayed effects of heat treatments on *Drosophila* male fertility (Canal Domenech & Fricke, 2022; Meena et al., 2024; Parratt et al., 2021). We therefore investigated the effects of delayed mating (at intervals of 1, 24, 48, or 72 hr post-treatment) following heat shock treatments (4 hr at 38 °C or 39 °C; 1 hr at 41 °C or 42 °C) in *Ae. aegypti* males on the reproductive performance of their female mates. As the mating interval did not influence any trait in the control group (Figure 4), controls were excluded from subsequent analyses to focus on treatment-specific effects.

**Figure 4.**
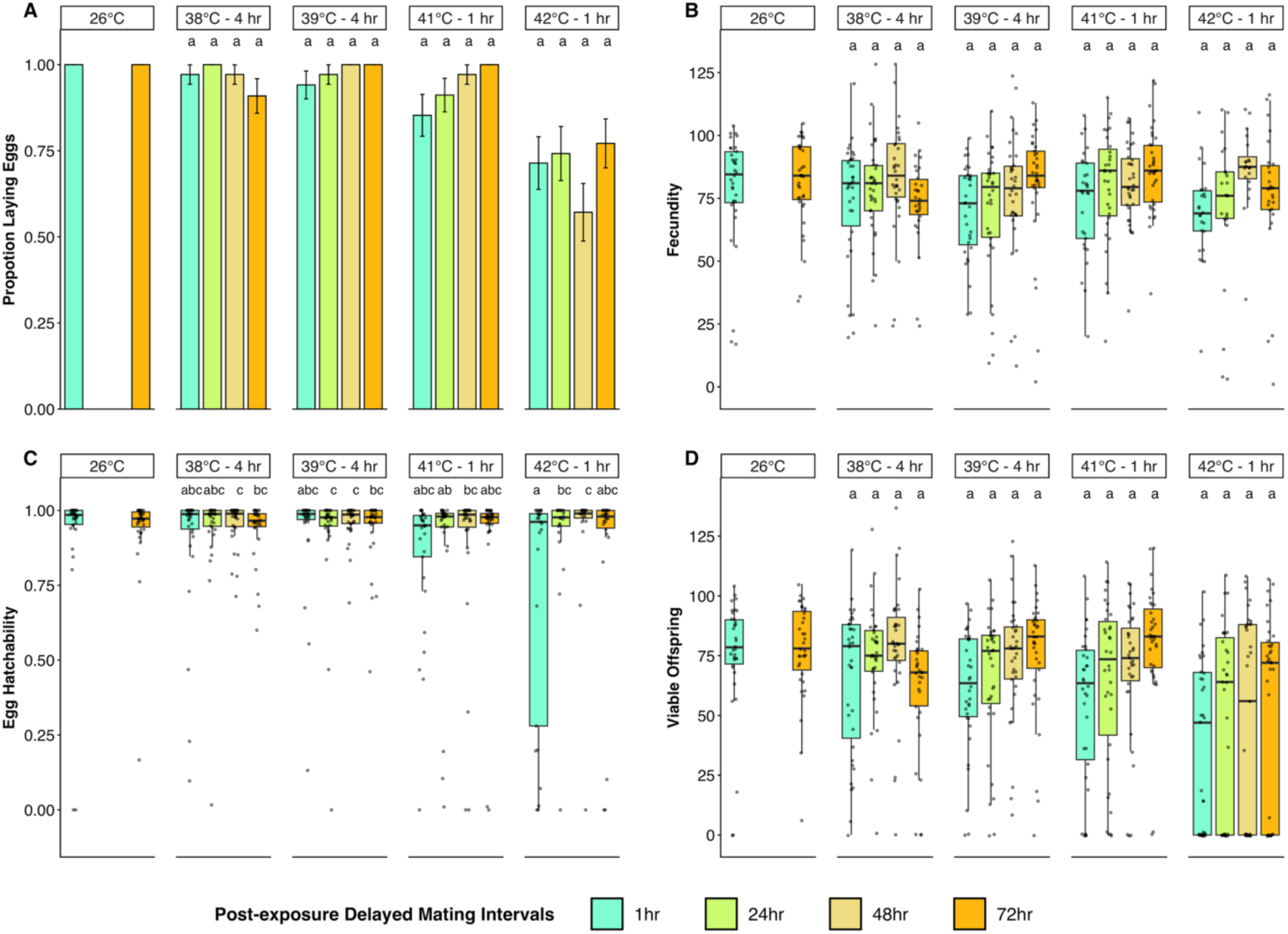
Effects of delayed mating on the reproductive success of male *Ae. aegypti* following sublethal heat shock exposure. We measured the proportion of females laying eggs (A), fecundity (B), egg hatchability (C), and the number of viable offspring (D). Four heat shock treatments (38, 39, 41, and 42 °C) were tested across four delayed mating intervals (1, 24, 48, and 72 hr). Untreated controls (26 °C) were assessed after 1 and 24 hr in parallel with heat-treated groups. Bars in A represent mean proportions with error bars indicating 95% confidence intervals. Boxplots in B – D show medians and interquartile ranges; grey overlaid jittered dots represent individual observations. Different lowercase letters indicate significant differences; shared letters indicate no significant difference (*p* < 0.05).

The proportion of females laying eggs was significantly affected by male heat shock treatment (χ^2^ (3) = 38.417, *p* < 0.001, Figure 4A; Table S5). Specifically, males subjected to 1 hr at 42 °C exhibited the lowest frequencies of successful egg laying (Figure 4A). This trait was not significantly influenced by the mating interval (χ^2^ (3) = 1.569, *p* = 0.666) or by an interaction between heat treatment and mating interval (χ^2^ (9) = 10.892, *p* = 0.283) (Table S5). In contrast, both fecundity (χ^2^ (3) = 15.207, *p* = 0.002, Figure 4B) and the number of viable offspring (χ^2^ (3) = 14.638, *p* = 0.002, Figure 4D) were primarily driven by mating interval, where fertility tended to increase with an increased duration between the heat shock and mating. This effect occurred independently of the initial heat shock treatment for both fecundity and the number viable offspring (all *p* > 0.243) (Table S5).

Egg hatchability was significantly affected by both heat shock treatment (χ^2^ (3) = 11.188, *p* = 0.011) and the mating interval (χ^2^ (3) = 27.029, *p* < 0.001), with a significant interaction between these factors (χ^2^ (9) = 21.311, *p* = 0.011) (Figure 4C, Table S5). In post-hoc comparisons, females that mated with males exposed to 42 °C for 1 hr had significantly lower hatchability when mating 1 hr post-exposure compared to mating at 24 hr (1 hr vs 24 hr: OR = 0.108, *z* = -4.416, *p* = 0.001) or 48 hr (1 hr vs 48 hr: OR = 0.070, *z* = -4.766, *p* < 0.001) post-exposure (Table S6). These results demonstrate a clear recovery of male reproductive fitness, whereby extending the post-exposure interval to 24-48 hr mitigates the immediate reduction in fertility induced by extreme heat.

### Effect of female mating status on heat shock induced changes in *Ae. aegypti* fertility

In the initial experiment in females, we only considered impacts of heat shock prior to mating, which ignores potential impacts on stored sperm (Walsh et al., 2022). We therefore assessed the impact of mating status (defined as heat exposure during the pre-mating or post-mating period) on the reproductive success of *Ae. aegypti* females following a sublethal heat shock. We also considered potential delayed effects by evaluating traits across two consecutive gonotrophic cycles.

There was no significant effect of treatment (defined as each combination of heat shock and timing of heat exposure) on egg laying success (χ^2^ (2) = 0.970, *p* = 0.616; Figure 5A). In contrast, significant effects were detected for fecundity (χ^2^ (2) = 8.539, *p* = 0.014; Figure 5B), egg hatchability (χ^2^ (2) = 6.388, *p* = 0.041; Figure 5C) and viable offspring production (χ^2^ (2) = 8.744, *p* = 0.013; Figure 5D) (Table S7). These effects primarily reflect differences between control and heat-shocked females rather than the timing of heat exposure (pre-mating vs post-mating), as pairwise comparisons among mating status groups were not significant (all *p* > 0.282; Table S8). Fecundity declined in the second gonotrophic cycle compared to the first (χ^2^ (1) = 9.891, *p* = 0.002; Figure 5B), with no interaction effects (χ^2^ (2) = 0.800, *p* = 0.670) (Table S7), suggesting independent, additive effects of heat shock and physiological aging.

**Figure 5.**
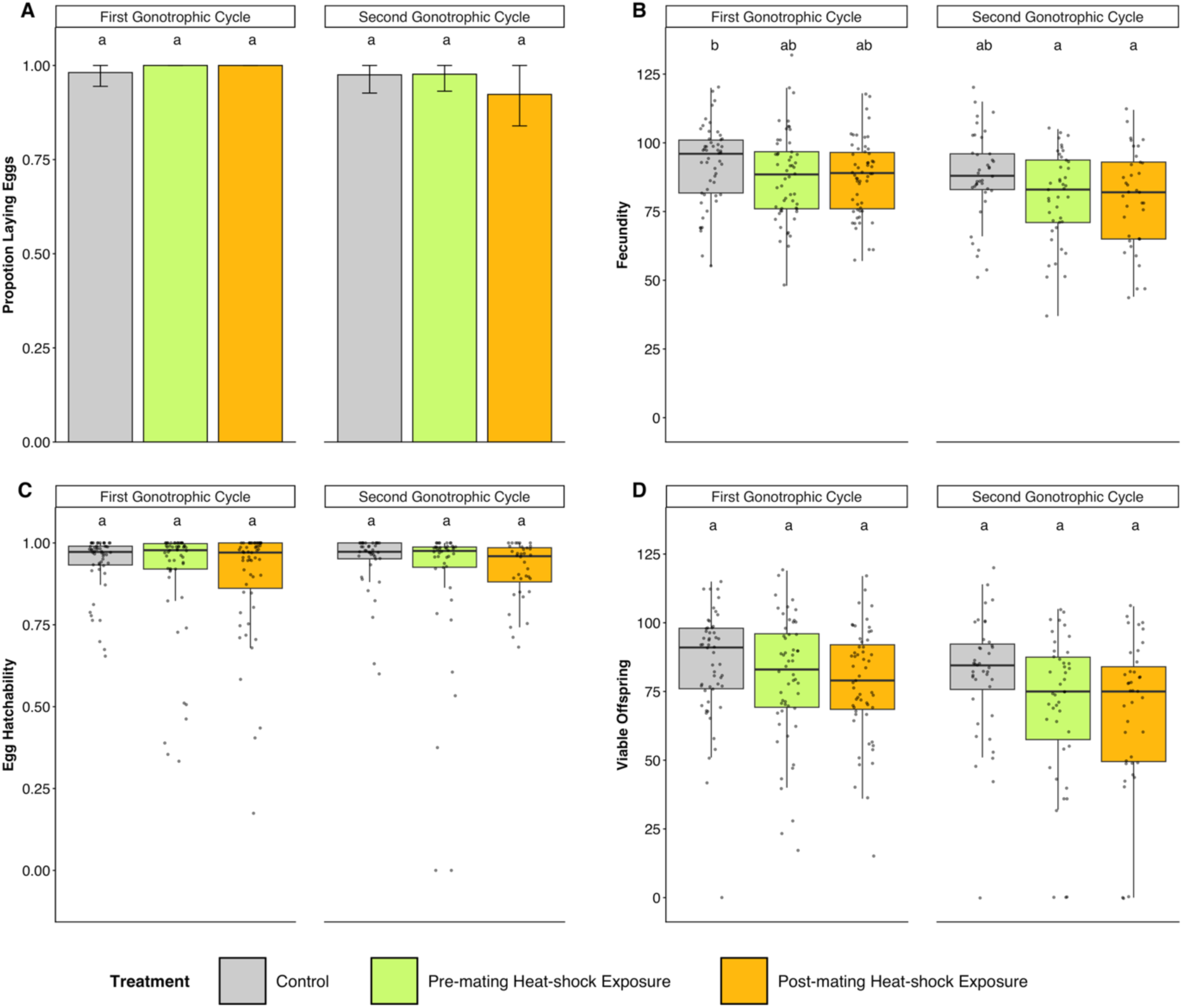
Effects of sublethal heat shock exposure (43°C) during the pre-mating and post-mating periods on the reproductive success of female *Ae. aegypti*. We measured the proportion of females laying eggs (A), fecundity (B), egg hatchability (C) and the number of viable offspring (D) across two consecutive gonotrophic cycles. Untreated controls are shown in grey, while sublethal heat shock treatments are shown in other colours. Bars in A represent mean proportions with error bars indicating 95% confidence intervals. Boxplots in B – D show medians and interquartile ranges; grey overlaid jittered dots represent individual observations. Different lowercase letters indicate significant differences; shared letters indicate no significant difference (*p* < 0.05).

## Transgenerational effects of heat shock exposure in *Ae. aegypti* fertility

We then investigated the effects of maternal sublethal heat shock exposure (41 °C) on *Ae. aegypti* offspring reproductive performance under sublethal heat stress (42 °C and 43 °C, with a 26 °C control). Our results demonstrate trait-specific transgenerational effects, with maternal heat exposure significantly increasing offspring fecundity (χ^2^ (1) = 11.795, *p* < 0.001; Figure 6B) and viable offspring production (χ^2^ (1) = 8.846, *p* = 0.003; Figure 6D), but not egg laying success or egg hatchability (all *p* > 0.124; Figure 6A, C) (Table S9). Post-hoc comparisons indicated that these differences were most pronounced when offspring were exposed to 42 °C, with offspring from heat-treated mothers producing significantly more eggs (heat-treated mothers vs untreated mothers: OR = 1.174, *z* = 3.889, *p* = 0.001) and viable offspring (heat-treated mothers vs untreated mothers: OR = 1.174, *z* = 3.378, *p* = 0.010) than offspring from untreated mothers (Table S10).

**Figure 6.**
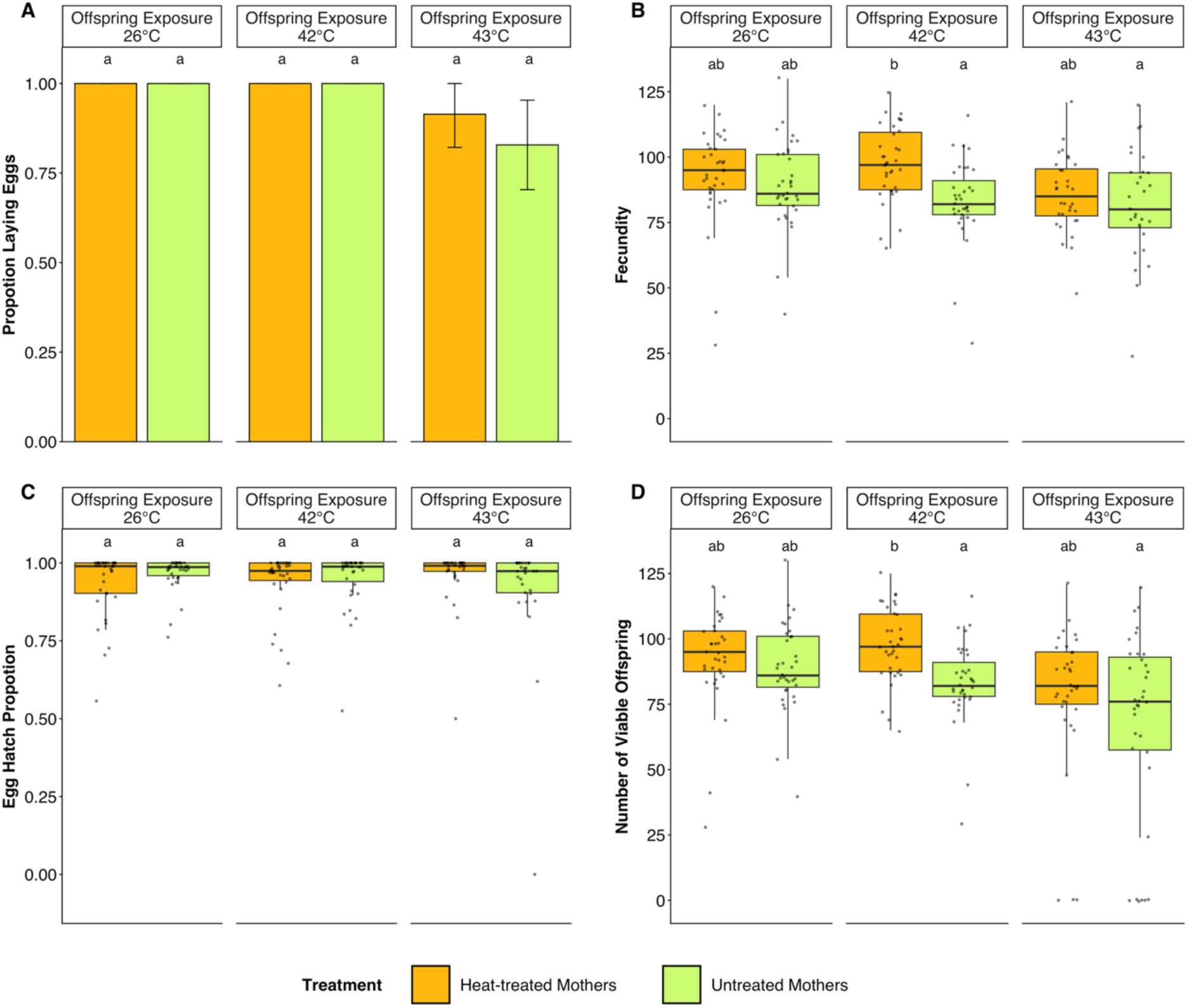
Effects of maternal heat shock exposure on the reproductive success of *Ae. aegypti* offspring. We measured the proportion of female offspring laying eggs (A), fecundity (B), egg hatchability (C) and the number of viable offspring (D). Offspring reproductive success was assessed at 26 °C (control) and under two sublethal heat treatments (42 °C and 43 °C). Bars in A represent mean proportions with error bars indicating 95% confidence intervals. Boxplots in B – D show medians and interquartile ranges; grey overlaid jittered dots represent individual observations. Different lowercase letters indicate significant differences; shared letters indicate no significant difference (*p* < 0.05).

Offspring treatment significantly affected offspring egg laying success (χ^2^ (2) = 8.945, *p* = 0.011; Figure 6A), fecundity (χ^2^ (2) = 10.204, *p* = 0.006; Figure 6B) and viable offspring production (χ^2^ (2) = 6.637, *p* = 0.036; Figure 6D), but not egg hatchability (χ^2^ (2) = 2.549, *p* = 0.280; Figure 6C) (Table S11). Among offspring from heat-treated mothers, fecundity declined at 43 °C compared to 42 °C (42 °C vs 43 °C: OR = 1.139, *z* = 3.416, *p* = 0.008; Table S10), suggesting a direct effect of extreme heat stress. Moreover, all interaction terms were non-significant (all *p* > 0.095; Table S9).

## Discussion

Understanding how acute thermal extremes influence insect fertility has become increasingly important as climate change intensifies the frequency, intensity and duration of heat-wave events (Buckley & Huey, 2016; Calvin et al., 2023). While recent studies have shown that fertility often declines at temperatures well below those causing mortality in *Drosophila* (Parratt et al., 2021; van Heerwaarden & Sgro, 2021) and other insects (Walsh et al., 2019; Weaving et al., 2024), the extent to which this phenomenon applies to mosquitoes has remained largely unexplored. Across both sexes, pupal and adult developmental stages, and multiple reproductive traits (egg-laying success, fecundity, egg hatchability, and viable offspring production) of three epidemiologically important mosquito species (*Ae. aegypti*, *Ae. notoscriptus* and *Cx*. *quinquefasciatus*), acute heat exposure rarely caused widespread reproductive collapse. Instead, there were subtle responses that varied among reproductive traits, sexes, developmental stages and species. Heat-induced male fertility impairment in *Ae. aegypti* was transient and largely recovered within 24–48 hours following heat exposure, whereas parental heat exposure modestly enhanced offspring reproductive performance under subsequent acute sublethal stress. Collectively, these findings demonstrate that mosquito fertility is not changed markedly under acute sublethal heat stress, but there are some species and sex specific effects that could influence mosquito population dynamics.

## Mosquito fertility is resilient to acute sublethal heat stress near upper thermal limits

A key finding is the resilience of mosquito fertility to acute sublethal heat stress, despite exposure to temperatures approaching each species’ upper thermal limits. This contrasts with findings in *Drosophila* (Parratt et al., 2021; van Heerwaarden & Sgro, 2021) and other insects (Walsh et al., 2021), where fertility often declines before survival is affected. Our findings suggest that thermal fertility limits relative to mortality limits may vary among insect lineages, potentially reflecting differences in their ecological and evolutionary history. This resilience may reflect adaptation to the heterogeneous thermal environments experienced by mosquitoes throughout their life cycle. Immature stages develop in small, shallow aquatic habitats where temperatures fluctuate markedly throughout the day (Richardson et al., 2011), while adults routinely alternate between exposed environments during host seeking and cooler resting habitats (Carrington et al., 2013; Vinauger & Chandrasegaran, 2024). Such fluctuating environments are characterised by repeated but relatively brief thermal extremes interspersed with opportunities for physiological recovery (Carrington et al., 2013; Colinet et al., 2015), which may favour adaptations that minimise reproductive disruption during transient heat events rather than prolonged exposure to elevated temperatures (Colinet et al., 2015). Consistent with this interpretation, the modest fertility losses observed following acute heat exposure contrast with the stronger reproductive effects reported under chronic heat exposure in mosquitoes (Pekľanská et al., 2025), indicating that both the duration and intensity of thermal exposure influence reproductive outcomes. Although the physiological mechanisms were not investigated directly here, rapid induction of heat-shock proteins, antioxidant defences and cellular repair pathways may protect reproductive tissues from irreversible thermal damage (Benoit et al., 2011; Colinet et al., 2015; Zhao et al., 2010; Zhao et al., 2014).

### Reproductive responses to acute heat stress are trait, sex, stage and species-specific

Although mosquitoes remained mostly fertile under acute sublethal heat stress, reproductive responses differed among traits, sexes, developmental stages and species, demonstrating that the reproductive consequences of acute heat stress are context dependent, as has also been reported in other insects (Meena et al., 2025; Walsh et al., 2021; Zhao et al., 2017). Acute heat exposure disrupted different components of the reproductive process depending on whether stress was experienced by males or females, highlighting the importance of assessing multiple reproductive traits when evaluating thermal fertility limits rather than relying on a single measure of reproductive performance (Walsh et al., 2019).

Female heat exposure primarily reduced fecundity while having little effect on egg-laying success or egg hatchability across the three mosquito species. This pattern suggests that acute heat stress primarily affected processes associated with egg production rather than oviposition behaviour or the ability of fertilised eggs to complete embryonic development successfully. Because oogenesis, vitellogenesis and nutrient allocation are energetically demanding processes that occur over several days following blood feeding (Attardo et al., 2005; Valzania et al., 2019), they may be more susceptible to transient physiological stress than oviposition. In contrast, male heat exposure had little effect on the fecundity of their untreated female mates but reduced egg hatchability and, in some species, viable offspring production. This suggests that male thermal stress may affect fertilisation success through impaired sperm quality, viability or function, consistent with evidence that male reproductive processes, particularly spermatogenesis, are particularly thermally sensitive (Meena et al., 2024; Parratt et al., 2021; Walsh et al., 2019). Developmental stage also influenced reproductive responses, although these effects varied among species and sexes.

## Heat-induced male reproductive impairment is largely transient, but some effects persist

Exposure to 42 °C for 1 hour caused a marked reduction in male reproductive performance depending on the trait considered. While egg hatchability recovered substantially within 24-48 hours after heat exposure, the proportion of females laying eggs remained lower regardless of mating interval. Because the temperatures used in this study approached the upper thermal limits of *Ae. aegypti* males (Pekľanská et al., 2025; Warusawithana et al., 2026), these findings suggest that acute heat stress near the critical thermal maximum can induce both transient and persistent reproductive effects.

The physiological mechanisms underlying these responses remain unclear, although temporary impairment of sperm motility or viability may explain the initial reduction in hatchability immediately after heat exposure (Porcelli et al., 2017; Sales et al., 2021; Walsh et al., 2019). Recovery may reflect restoration of sperm function, repair of reversible heat-induced damage, or replacement of thermally compromised sperm by newly produced sperm following cessation of thermal stress (Walsh et al., 2019). In contrast, the persistent reduction in egg-laying proportion suggests effects beyond the recovery of sperm function. Transient reductions in male fertility followed by recovery have been reported in *Drosophila* (Canal Domenech & Fricke, 2022) and other insects (Baur et al., 2021; Chevrier et al., 2019), whereas repeated heat stress can produce more persistent fertility impairment in *Drosophila melanogaster* (Meena et al., 2024) and other insects (Sales et al., 2018). Additionally, we found no evidence for delayed costs which have also been reported previously in *Drosophila* (Parratt et al., 2021). These findings suggest that the reproductive consequences of acute heat stress depend not only on the magnitude of thermal stress but also on the timing of mating relative to heat exposure, and post-stress recovery dynamics should be incorporated into predictions of mosquito population responses to heatwave events under climate change (Calvin et al., 2023).

## Female reproductive responses are largely independent of mating status

Unlike males, whose fertility recovered rapidly following acute heat stress, female reproductive responses to acute heat stress were largely independent of whether exposure occurred before or after mating. Because the principal difference between these treatments was whether sperm had been transferred and stored before heat exposure, any effects of acute heat stress on stored sperm appear insufficient to explain the observed reductions in reproductive performance. This contrasts with studies in *Drosophila* and other insects, where female reproductive responses to heat stress can depend on mating status and mating system (Baur et al., 2021; Meena et al., 2025), and where heat-induced damage to sperm stored within the female reproductive tract can contribute to reduced fertility (Sales et al., 2018; Walsh et al., 2022) The persistence of reduced fertility into the second gonotrophic cycle further suggests that the effects of acute heat exposure were not restricted to the immediate reproductive event, potentially reflecting cumulative physiological costs associated with successive gonotrophic cycles or age-related declines in reproductive performance (Martin et al., 2025).

## Parental heat exposure confers limited transgenerational benefits

Parental exposure to acute sublethal heat stress modestly improved offspring reproductive performance, with fecundity and viable offspring production increasing by approximately 4-17% depending on offspring temperature. Transgenerational responses to thermal stress have been reported across a range of insect taxa, although their magnitude and direction vary considerably (Cavieres et al., 2019; Donelson et al., 2017; Green et al., 2019; Sales et al., 2018; Zhu et al., 2021). For example, parental heatwaves reduced offspring reproductive performance in *Tribolium castaneum* (flour beetle) (Sales et al., 2018), whereas maternal heat exposure improved offspring thermal performance in *Drosophila suzukii* (Green et al., 2019). Our findings, together with previous studies, suggest that parental thermal history can influence offspring responses to elevated temperatures, although the magnitude and persistence of these effects appear to be context dependent (Donelson et al., 2017; Zhu et al., 2021).

## Conclusion

Collectively, our findings show that reproductive responses of mosquitoes to acute heat stress at temperatures approaching their upper thermal limits were generally modest and varied among reproductive traits, sexes, developmental stages, species, the timing of heat exposure, and parental thermal history. Rather than causing widespread reproductive collapse, acute heat stress produced context-dependent responses, highlighting the importance of assessing multiple components of reproduction when evaluating thermal vulnerability and predicting mosquito responses to climate change. Future studies could investigate the cumulative effects of repeated heat exposure under ecologically realistic fluctuating thermal environments while integrating physiological and molecular approaches to distinguish between acute and chronic thermal responses. Incorporating these processes into predictive models could improve forecasts of mosquito abundance and distribution (Kearney et al., 2009), with potential implications for disease transmission under future climate change.

## Acknowledgements

We thank Leon Hugo and Scott Ritchie for providing mosquito populations used in the study.

## Author contributions

Conceptualization: A.L.W., B.v.H., A.A.H., P.A.R.; Formal analysis: A.L.W, B.v.H., A.A.H., P.A.R.; Funding acquisition: B.v.H., A.A.H., P.A.R.; Investigation: A.L.W.; Methodology: A.L.W., B.v.H., P.A.R.; Supervision: B.v.H., A.A.H., P.A.R.; Visualization: A.L.W; Writing – original draft: A.L.W., B.v.H., A.A.H., P.A.R.; Writing – review & editing: A.L.W., B.v.H., A.A.H., P.A.R.

## References

1. Agyekum, T. P., Arko-Mensah, J., Botwe, P. K., Hogarh, J. N., Issah, I., Dwomoh, D., Billah, M. K., Dadzie, S. K., Robins, T. G., & Fobil, J. N. (2022). Effects of Elevated Temperatures on the Growth and Development of Adult *Anopheles gambiae* (s.l.) (Diptera: Culicidae) Mosquitoes. J Med Entomol, 59(4), 1413–1420. 10.1093/jme/tjac046

2. Attardo, G. M., Hansen, I. A., & Raikhel, A. S. (2005). Nutritional regulation of vitellogenesis in mosquitoes: implications for anautogeny. Insect Biochem Mol Biol, 35(7), 661–675.

3. Bader, C. A., & Williams, C. R. (2013). Mating, ovariole number and sperm production of the dengue vector mosquito *Aedes aegypti* (L.) in Australia: broad thermal optima provide the capacity for survival in a changing climate. Physiol Entomol, 37(2), 136–144. 10.1111/j.1365-3032.2011.00818.x

4. Baur, J., Jagusch, D., Michalak, P., Koppik, M., & Berger, D. (2021). The mating system affects the temperature sensitivity of male and female fertility. Funct Ecol, 36(1), 92–106.

5. Benoit, J. B., Lopez-Martinez, G., Patrick, K. R., Phillips, Z. P., Krause, T. B., & Denlinger, D. L. (2011). Drinking a hot blood meal elicits a protective heat shock response in mosquitoes. Proc Natl Acad Sci U S A, 108(19), 8026–8029. 10.1073/pnas.1105195108

6. Brady, O. J., Godfray, H. C. J., Tatem, A. J., Gething, P. W., Cohen, J. M., McKenzie, F. E., Perkins, T. A., Reiner, R. C., Tusting, L. S., & Sinka, M. E. (2016). Vectorial capacity and vector control: reconsidering sensitivity to parameters for malaria elimination. Trans R Soc Trop Med Hyg, 110(2), 107–117.

7. Brady, O. J., & Hay, S. I. (2020). The global expansion of dengue: How *Aedes aegypti* mosquitoes enabled the first pandemic arbovirus. *Annu Rev Entomol*, 65, 191–208. 10.1146/annurev-ento-011019-024918

8. Buckley, L. B., & Huey, R. B. (2016). Temperature extremes: geographic patterns, recent changes, and implications for organismal vulnerabilities. Glob Chang Biol, 22(12), 3829–3842.

9. Calvin, K., Dasgupta, D., Krinner, G., Mukherji, A., Thorne, P. W., Trisos, C., Romero, J., Aldunce, P., Barrett, K., & Blanco, G. (2023). IPCC, 2023: Climate Change 2023: Synthesis Report. Contribution of Working Groups I, II and III to the Sixth Assessment Report of the Intergovernmental Panel on Climate Change [Core Writing Team, H. Lee and J. Romero (eds.)]. *IPCC*, *Geneva*, *Switzerland* ((No Title), Issue.

10. Canal Domenech, B., & Fricke, C. (2022). Recovery from heat-induced infertility—A study of reproductive tissue responses and fitness consequences in male Drosophila melanogaster. Ecol evol, 12(12), e9563.

11. Carrington, L. B., Armijos, M. V., Lambrechts, L., Barker, C. M., & Scott, T. W. (2013). Effects of fluctuating daily temperatures at critical thermal extremes on *Aedes aegypti* life-history traits. PLoS One, 8(3), e58824. 10.1371/journal.pone.0058824

12. Cavieres, G., Alruiz, J. M., Medina, N. R., Bogdanovich, J. M., & Bozinovic, F. (2019). Transgenerational and within-generation plasticity shape thermal performance curves. *Ecol evol*, 9(4), 2072-2082. https://pmc.ncbi.nlm.nih.gov/articles/PMC6392392/

13. Chandrasegaran, K., Lahondère, C., Escobar, L. E., & Vinauger, C. (2020). Linking mosquito ecology, traits, behavior, and disease transmission. Trends in parasitol, *36*(4), 393-403.

14. Chevrier, C., Nguyen, T. M., & Bressac, C. (2019). Heat shock sensitivity of adult male fertility in the parasitoid wasp *Anisopteromalus calandrae* (Hymenoptera, Pteromalidae). J Therm Biol, 85, 102419. 10.1016/j.jtherbio.2019.102419

15. Claflin, S. B., & Webb, C. E. (2015). Ross River virus: Many vectors and unusual hosts make for an unpredictable pathogen. PLoS Pathog, 11(9), e1005070. 10.1371/journal.ppat.1005070

16. Clements, A. N. (2023). The biology of mosquitoes, Volume 1: Development, nutrition and reproduction. Cabi GB.

17. Colinet, H., Sinclair, B. J., Vernon, P., & Renault, D. (2015). Insects in fluctuating thermal environments. *Annu Rev Entomol*, 60, 123–140. 10.1146/annurev-ento-010814-021017

18. Costanzo, K., & Occhino, D. (2023). Effects of Temperature on Blood Feeding and Activity Levels in the Tiger Mosquito, *Aedes albopictus*. Insects, 14(9). 10.3390/insects14090752

19. Delatte, H., Gimonneau, G., Triboire, A., & Fontenille, D. (2009). Influence of temperature on immature development, survival, longevity, fecundity, and gonotrophic cycles of *Aedes albopictus*, vector of chikungunya and dengue in the Indian Ocean. J Med Entomol, 46(1), 33–41. 10.1603/033.046.0105

20. Donelson, J. M., Salinas, S., Munday, P. L., & Shama, L. N. S. (2017). Transgenerational plasticity and climate change experiments: Where do we go from here? Glob Chang Biol, 24(1), 13–34. 10.1111/gcb.13903

21. Ezeakacha, N. F., & Yee, D. A. (2019). The role of temperature in affecting carry-over effects and larval competition in the globally invasive mosquito *Aedes albopictus*. Parasit Vectors, 12, 1–11.

22. Fox, J., & Weisberg, S. (2018). An R companion to applied regression. Sage publications.

23. García-Robledo, C., & Baer, C. S. (2021). Positive genetic covariance and limited thermal tolerance constrain tropical insect responses to global warming. J Evol Biol, 34(9), 1432–1446.

24. Green, C. K., Moore, P. J., & Sial, A. A. (2019). Impact of heat stress on development and fertility of *Drosophila suzukii* Matsumura (Diptera: Drosophilidae). J Insect Physiol, 114, 45–52.

25. Hartig, F. (2016). DHARMa: residual diagnostics for hierarchical (multi-level/mixed) regression models. CRAN: contributed packages.

26. Hernández, B. J., Barrientos, L. M., Vélez, E. V., Guzmán, P. A., Alfonso-Parra, C., & Avila, F. W. (2026). Developmental heat stress disrupts male-induced female post-mating responses in the dengue vector mosquito, Aedes aegypti. Parasit Vectors (uder review).

27. Hoffmann, A. A., & Bridle, J. (2022). The dangers of irreversibility in an age of increased uncertainty: revisiting plasticity in invertebrates. Oikos, 2022(4), e08715.

28. Holzmann, K. L., Schmitzer, T., Abels, A., Čorkalo, M., Mitesser, O., Kortmann, M., Alonso-Alonso, P., Correa-Carmona, Y., Pinos, A., & Yon, F. (2026). Limited thermal tolerance in tropical insects and its genomic signature. Nature.

29. Jansen, C. C., & Beebe, N. W. (2010). The dengue vector *Aedes aegypti*: what comes next. Microbes Infect, 12(4), 272–279. 10.1016/j.micinf.2009.12.011

30. Kearney, M., Porter, W. P., Williams, C., Ritchie, S., C Hoffmann, A. A. (2009). Integrating biophysical models and evolutionary theory to predict climatic impacts on species’ ranges: the dengue mosquito *Aedes aegypti* in Australia. Funct Ecol, 23(3), 528–538.

31. King, A. D., Black, M. T., Min, S. K., Fischer, E. M., Mitchell, D. M., Harrington, L. J., C Perkins-Kirkpatrick, S. E. (2016). Emergence of heat extremes attributable to anthropogenic influences. Geophys Res Lett, 43(7), 3438–3443.

32. Kosmidis, I., & Firth, D. (2021). Jeffreys-prior penalty, finiteness and shrinkage in binomial-response generalized linear models. Biometrika, 108(1), 71–82.

33. Lenth, R. (2023). emmeans: Estimated Marginal Means, aka Least-Squares Means_. R package version 1.8. 5.

34. Martin, L. E., Estévez-Lao, T. Y., McCabe, T. C., & Hillyer, J. F. (2025). Warmer temperature accelerates reproductive senescence in mosquitoes. Front Physiol, 16, 1610310.

35. McGillycuddy, M., Popovic, G., Bolker, B. M., & Warton, D. I. (2025). Parsimoniously fitting large multivariate random effects in glmmTMB. J Stat Softw, 112, 1–19.

36. Mee, P. T., Buultjens, A. H., Oliver, J., Brown, K., Crowder, J. C., Porter, J. L., Hobbs, E. C., Judd, L. M., Taiaroa, G., & Puttharak, N. (2024). Mosquitoes provide a transmission route between possums and humans for Buruli ulcer in southeastern Australia. *Nat Microbiol*, 9(2), 377–389. https://www.nature.com/articles/s41564-023-01553-1.pdf

37. Meena, A., De Nardo, A. N., Maggu, K., Sbilordo, S. H., Roy, J., Snook, R. R., & Lüpold, S. (2024). Fertility loss and recovery dynamics after repeated heat stress across life stages in male *Drosophila melanogaster*: patterns and processes. R Soc Open Sci, 11(10), 241082.

38. Meena, A., Maggu, K., De Nardo, A. N., Kovalov, V., Eggs, B., & Lüpold, S. (2025). Stage-specific and cumulative effects of heat stress on male and female reproductive performance in *Drosophila melanogaster*. J Therm Biol, 104213.

39. Mitchell, S. N., & Catteruccia, F. (2017). Anopheline reproductive biology: impacts on vectorial capacity and potential avenues for malaria control. Cold Spring Harb Perspect Med, 7(12), a025593.

40. Mordecai, E. A., Caldwell, J. M., Grossman, M. K., Lippi, C. A., Johnson, L. R., Neira, M., Rohr, J. R., Ryan, S. J., Savage, V., Shocket, M. S., Sippy, R., Stewart Ibarra, A. M., Thomas, M. B., & Villena, O. (2019). Thermal biology of mosquito-borne disease. Ecol Lett, 22(10), 1690–1708. 10.1111/ele.13335

41. Mourya, D., Yadav, P., & Mishra, A. (2004). Effect of temperature stress on immature stages and susceptibility of *Aedes aegypti* mosquitoes to chikungunya virus. Am J Trop Med Hyg, 70(4), 346–350.

42. Negi, C., & Verma, P. (2018). Review on *Culex quinquefasciatus*: Southern House Mosquito. Int J Life-sci Res, 4(1), 1563–1566. 10.21276/ijlssr.2018.4.1.9

43. Padde, J. R., Zhou, Y., Chen, Y., Zhu, Y., Yang, Y., Hou, M., Chen, L., Xu, Z., Zhang, D., Chen, L., & Ji, M. (2024). Adaptation and carry over effects of extreme sporadic heat stress in *Culex* mosquitoes. Acta Trop, 260, 107417. 10.1016/j.actatropica.2024.107417

44. Parratt, S. R., Walsh, B. S., Metelmann, S., White, N., Manser, A., Bretman, A. J., Hoffmann, A. A., Snook, R. R., & Price, T. A. (2021). Temperatures that sterilize males better match global species distributions than lethal temperatures. Nat Clim Change, 11(6), 481–484.

45. Pekľanská, M., van Heerwaarden, B., Hoffmann, A. A., Nouzová, M., Šíma, R., & Ross, P. A. (2025). Elevated developmental temperatures below the lethal limit reduce *Aedes aegypti* fertility. J Exp Biol, 228(3), JEB249803.

46. Porcelli, D., Gaston, K. J., Butlin, R. K., & Snook, R. R. (2017). Local adaptation of reproductive performance during thermal stress. J Evol Biol, 30(2), 422–429.

47. Richardson, K., Hoffmann, A. A., Johnson, P., Ritchie, S., & Kearney, M. R. (2011). Thermal sensitivity of *Aedes aegypti* from Australia: empirical data and prediction of effects on distribution. J Med Entomol, 48(4), 914–923.

48. Rúa, G. L., Quiñones, M. L., Vélez, I. D., Zuluaga, J. S., Rojas, W., Poveda, G., & Ruiz, D. (2005). Laboratory estimation of the effects of increasing temperatures on the duration of gonotrophic cycle of *Anopheles albimanus* (Diptera: Culicidae). *Mem Inst Oswaldo Cruz*, *100*, 515-520. https://www.scielo.br/j/mioc/a/f8b4bDtgB8ǪmVsMPmFX4C5H/?lang=enCformat=pdf

49. Sales, K., Vasudeva, R., Dickinson, M. E., Godwin, J. L., Lumley, A. J., Michalczyk, Ł., Hebberecht, L., Thomas, P., Franco, A., & Gage, M. J. (2018). Experimental heatwaves compromise sperm function and cause transgenerational damage in a model insect. *Nat Commun*, 9(1), 4771.

50. Sales, K., Vasudeva, R., & Gage, M. J. G. (2021). Fertility and mortality impacts of thermal stress from experimental heatwaves on different life stages and their recovery in a model insect. R Soc Open Sci, 8(3), 201717. 10.1098/rsos.201717

51. Schoof, H. (1967). Mating, resting habits and dispersal of *Aedes aegypti*. Bull World Health Organ, 36(4), 600.

52. Scott, T. W., Amerasinghe, P. H., Morrison, A. C., Lorenz, L. H., Clark, G. G., Strickman, D., Kittayapong, P., & Edman, J. D. (2000). Longitudinal studies of *Aedes aegypti* (Diptera: Culicidae) in Thailand and Puerto Rico: blood feeding frequency. J Med Entomol, 37(1), 89–101. 10.1603/0022-2585-37.1.89

53. Valzania, L., Mattee, M. T., Strand, M. R., & Brown, M. R. (2019). Blood feeding activates the vitellogenic stage of oogenesis in the mosquito *Aedes aegypti* through inhibition of glycogen synthase kinase 3 by the insulin and TOR pathways. J Dev Biol, 454(1), 85–95.

54. van Heerwaarden, B., & Sgro, C. M. (2021). Male fertility thermal limits predict vulnerability to climate warming. Nat Commun, 12(1), 2214. 10.1038/s41467-021-22546-w

55. Vinauger, C., & Chandrasegaran, K. (2024). Context-specific variation in life history traits and behavior of *Aedes aegypti* mosquitoes. Front Insect Sci, 4, 1426715.

56. Walsh, B. S., Mannion, N. L., Price, T. A., & Parratt, S. R. (2021). Sex-specific sterility caused by extreme temperatures is likely to create cryptic changes to the operational sex ratio in *Drosophila virilis*. *Curr Zool*, 67(3), 341–343.

57. Walsh, B. S., Parratt, S. R., Hoffmann, A. A., Atkinson, D., Snook, R. R., Bretman, A., & Price, T. A. R. (2019). The Impact of Climate Change on Fertility. Trends Ecol Evol, 34(3), 249–259. 10.1016/j.tree.2018.12.002

58. Walsh, B. S., Parratt, S. R., Snook, R. R., Bretman, A., Atkinson, D., & Price, T. A. (2022). Female fruit flies cannot protect stored sperm from high temperature damage. J Therm Biol, 105, 103209.

59. Wang, X., & Li, J. (2024). Comparative Physiology of Mosquito Reproductive Systems. J Mosq Res, 14.

60. Warusawithana, A. L., van Heerwaarden, B., Hoffmann, A. A., & Ross, P. A. (2026). Heat hardening enhances mosquito heat tolerance in a species-specific and assay-specific manner. J Therm Biol, 104394.

61. Weaving, H., Terblanche, J. S., & English, S. (2024). Heatwaves are detrimental to fertility in the viviparous tsetse fly. Proc R Soc B: Biol Sci, 291(2018).

62. Wickham, H. (2016). Elegant graphics for data analysis. In: Springer.

63. Wickham, H., Averick, M., Bryan, J., Chang, W., McGowan, L. D. A., François, R., Grolemund, G., Hayes, A., Henry, L., & Hester, J. (2019). Welcome to the Tidyverse (R package version 2.0.0). J open source softw, 4(43), 1686.

64. Zhao, F., Hoffmann, A. A., Xing, K., & Ma, C.-s. (2017). Life stages of an aphid living under similar thermal conditions differ in thermal performance. *J Insect Physiol*, SS, 1–7.

65. Zhao, L., Becnel, J. J., Clark, G. G., & Linthicum, K. J. (2010). Expression of AeaHsp26 and AeaHsp83 in *Aedes aegypti* (Diptera: Culicidae) larvae and pupae in response to heat shock stress. J Med Entomol, 47(3), 367–375. 10.1603/me09232

66. Zhao, L., Pridgeon, J. W., Becnel, J. J., Clark, G. G., & Linthicum, K. J. (2014). Identification of genes differentially expressed during heat shock treatment in *Aedes aegypti*. J Med Entomol, 46(3), 490–495.

67. Zhu, L., Hoffmann, A. A., Li, S. M., & Ma, C. S. (2021). Extreme climate shifts pest dominance hierarchy through thermal evolution and transgenerational plasticity. Funct Ecol, 35(7), 1524–1537.

