## Supplementary data for "Mosquito fertility is (mostly) resilient to sublethal heat shocks"

1. **Sublethal heat shock effects on female fertility (*Ae. aegypti, Ae. notoscriptus* and *Cx. quinquefasciatus*)** (Table S1– S2)

Table S1. Analysis of deviance (Type II Wald Chi-square tests) of reproductive traits evaluating the effect of sublethal heat shock on female fertility

| Spp. | Trait | Factor | df | χ^2^ | *p*-value |  |
| --- | --- | --- | --- | --- | --- | --- |
| *Aedes aegypti* | |  |  |  |  |  |
|  | egg laying success | treatment | 4 | 1.196 | 0.879 |  |
|  | fecundity | treatment | 4 | 18.703 | < 0.001 | ^***^ |
|  | egg hatchability | treatment | 4 | 7.960 | 0.093 | **^⋅^** |
|  | viable offspring | treatment | 4 | 8.873 | 0.064 | **^⋅^** |
| *Aedes notoscriptus* | |  |  |  |  |  |
|  | egg laying success | treatment | 4 | 0.719 | 0.949 |  |
|  | fecundity | treatment | 4 | 18.987 | < 0.001 | ^***^ |
|  | egg hatchability | treatment | 4 | 6.083 | 0.193 |  |
|  | viable offspring | treatment | 4 | 14.904 | 0.005 | ^**^ |
| *Culex quinquefasciatus* | |  |  |  |  |  |
|  | egg laying success | treatment | 4 | 6.305 | 0.178 |  |
|  | fecundity | treatment | 4 | 14.225 | 0.007 | ^**^ |
|  | egg hatchability | treatment | 4 | 3.985 | 0.408 |  |
|  | viable offspring | treatment | 4 | 13.467 | 0.009 | ^**^ |

Significant *p*-values are indicated with asterisks (^***^ *p* < 0.001, ^**^ *p* < 0.01, ^*^ *p* < 0.05, **^⋅^** *p* < 0.1)

Table S2. Full pairwise comparisons of reproductive traits evaluating the effects of heat shock exposure effect on female fertility

| Spp. | Trait | Comparison (treatment vs. treatment / °C) | | Ratio / OR | SE | Z-ratio | *p*-value |  |
| --- | --- | --- | --- | --- | --- | --- | --- | --- |
| *Aedes aegypti* | | |  |  |  |  |  |  |
|  | Egg laying success | | |  |  |  |  |  |
|  |  | control at 26 °C / pupal exposure at 42 °C | | 1.000 | 2.050 | 0.000 | 1.000 |  |
|  |  | control at 26 °C / pupal exposure at 43 °C | | 1.040 | 2.120 | 0.017 | 1.000 |  |
|  |  | control at 26 °C / adult exposure at 42 °C | | 1.040 | 2.120 | 0.017 | 1.000 |  |
|  |  | control at 26 °C / adult exposure at 43 °C | | 3.470 | 5.850 | 0.739 | 0.947 |  |
|  |  | pupal exposure at 42 °C / pupal exposure at 43 °C | | 1.040 | 2.120 | 0.017 | 1.000 |  |
|  |  | pupal exposure at 42 °C / adult exposure at 42 °C | | 1.040 | 2.120 | 0.017 | 1.000 |  |
|  |  | pupal exposure at 42 °C / adult exposure at 43 °C | | 3.470 | 5.850 | 0.739 | 0.947 |  |
|  |  | pupal exposure at 43 °C / adult exposure at 42 °C | | 1.000 | 2.050 | 0.000 | 1.000 |  |
|  |  | pupal exposure at 43 °C / adult exposure at 43 °C | | 3.350 | 5.650 | 0.718 | 0.953 |  |
|  |  | adult exposure at 42 °C / adult exposure at 43 °C | | 3.350 | 5.650 | 0.718 | 0.953 |  |
|  | Fecundity | | |  |  |  |  |  |
|  |  | control at 26 °C / pupal exposure at 42 °C | | 0.913 | 0.031 | -2.718 | 0.051 | ^*^  **^⋅^** |
|  |  | control at 26 °C / pupal exposure at 43 °C | | 1.042 | 0.041 | 1.045 | 0.835 |  |
|  |  | control at 26 °C / adult exposure at 42 °C | | 0.907 | 0.033 | -2.678 | 0.057 | **^⋅^** |
|  |  | control at 26 °C / adult exposure at 43 °C | | 0.929 | 0.040 | -1.706 | 0.430 |  |
|  |  | pupal exposure at 42 °C / pupal exposure at 43 °C | | 1.142 | 0.046 | 3.284 | 0.009 | ^**^ |
|  |  | pupal exposure at 42 °C / adult exposure at 42 °C | | 0.994 | 0.037 | -0.158 | 1.000 |  |
|  |  | pupal exposure at 42 °C / adult exposure at 43 °C | | 1.018 | 0.045 | 0.400 | 0.995 |  |
|  |  | pupal exposure at 43 °C / adult exposure at 42 °C | | 0.870 | 0.037 | -3.247 | 0.010 | ^*^ |
|  |  | pupal exposure at 43 °C / adult exposure at 43 °C | | 0.891 | 0.044 | -2.362 | 0.126 |  |
|  |  | adult exposure at 42 °C / adult exposure at 43 °C | | 1.024 | 0.047 | 0.509 | 0.987 |  |
|  | Egg hatchability | | |  |  |  |  |  |
|  |  | control at 26 °C / pupal exposure at 42 °C | | 2.223 | 0.999 | 1.778 | 0.386 |  |
|  |  | control at 26 °C / pupal exposure at 43 °C | | 2.447 | 1.050 | 2.095 | 0.222 |  |
|  |  | control at 26 °C / adult exposure at 42 °C | | 3.967 | 2.130 | 2.570 | 0.076 | **^⋅^** |
|  |  | control at 26 °C / adult exposure at 43 °C | | 1.636 | 0.770 | 1.045 | 0.834 |  |
|  |  | pupal exposure at 42 °C / pupal exposure at 43 °C | | 1.101 | 0.399 | 0.265 | 0.999 |  |
|  |  | pupal exposure at 42 °C / adult exposure at 42 °C | | 1.784 | 0.868 | 1.190 | 0.757 |  |
|  |  | pupal exposure at 42 °C / adult exposure at 43 °C | | 0.736 | 0.304 | -0.742 | 0.947 |  |
|  |  | pupal exposure at 43 °C / adult exposure at 42 °C | | 1.621 | 0.755 | 1.037 | 0.839 |  |
|  |  | pupal exposure at 43 °C / adult exposure at 43 °C | | 0.668 | 0.260 | -1.035 | 0.839 |  |
|  |  | adult exposure at 42 °C / adult exposure at 43 °C | | 0.412 | 0.209 | -1.749 | 0.404 |  |
|  | Viable offspring | | |  |  |  |  |  |
|  |  | control at 26 °C / pupal exposure at 42 °C | | 0.941 | 0.051 | -1.118 | 0.797 |  |
|  |  | control at 26 °C / pupal exposure at 43 °C | | 1.083 | 0.060 | 1.438 | 0.603 |  |
|  |  | control at 26 °C / adult exposure at 42 °C | | 0.954 | 0.053 | -0.864 | 0.910 |  |
|  |  | control at 26 °C / adult exposure at 43 °C | | 0.946 | 0.054 | -0.975 | 0.867 |  |
|  |  | pupal exposure at 42 °C / pupal exposure at 43 °C | | 1.151 | 0.064 | 2.545 | 0.081 | **^⋅^** |
|  |  | pupal exposure at 42 °C / adult exposure at 42 °C | | 1.013 | 0.056 | 0.244 | 0.999 |  |
|  |  | pupal exposure at 42 °C / adult exposure at 43 °C | | 1.006 | 0.057 | 0.101 | 1.000 |  |
|  |  | pupal exposure at 43 °C / adult exposure at 42 °C | | 0.880 | 0.049 | -2.282 | 0.151 |  |
|  |  | pupal exposure at 43 °C / adult exposure at 43 °C | | 0.873 | 0.050 | -2.352 | 0.129 |  |
|  |  | adult exposure at 42 °C / adult exposure at 43 °C | | 0.992 | 0.057 | -0.135 | 1.000 |  |
| *Aedes notoscriptus* | | |  |  |  |  |  |  |
|  | Egg laying success | | |  |  |  |  |  |
|  |  | control at 26 °C / pupal exposure at 40 °C | | 0.778 | 0.604 | -0.324 | 0.998 |  |
|  |  | control at 26 °C / pupal exposure at 41 °C | | 1.000 | 0.733 | 0.000 | 1.000 |  |
|  |  | control at 26 °C / adult exposure at 40 °C | | 0.841 | 0.656 | -0.222 | 1.000 |  |
|  |  | control at 26 °C / adult exposure at 41 °C | | 1.378 | 0.976 | 0.453 | 0.991 |  |
|  |  | pupal exposure at 40 °C / pupal exposure at 41 °C | | 1.286 | 0.999 | 0.324 | 0.998 |  |
|  |  | pupal exposure at 40 °C / adult exposure at 40 °C | | 1.082 | 0.887 | 0.096 | 1.000 |  |
|  |  | pupal exposure at 40 °C / adult exposure at 41 °C | | 1.772 | 1.330 | 0.760 | 0.942 |  |
|  |  | pupal exposure at 41 °C / adult exposure at 40 °C | | 0.841 | 0.656 | -0.222 | 1.000 |  |
|  |  | pupal exposure at 41 °C / adult exposure at 41 °C | | 1.378 | 0.976 | 0.453 | 0.991 |  |
|  |  | adult exposure at 40 °C / adult exposure at 41 °C | | 1.638 | 1.240 | 0.653 | 0.966 |  |
|  | Fecundity | | |  |  |  |  |  |
|  |  | control at 26 °C / pupal exposure at 40 °C | | 0.935 | 0.081 | -0.772 | 0.939 |  |
|  |  | control at 26 °C / pupal exposure at 41 °C | | 0.848 | 0.073 | -1.917 | 0.308 |  |
|  |  | control at 26 °C / adult exposure at 40 °C | | 0.974 | 0.104 | -0.243 | 0.999 |  |
|  |  | control at 26 °C / adult exposure at 41 °C | | 1.262 | 0.151 | 1.945 | 0.294 |  |
|  |  | pupal exposure at 40 °C / pupal exposure at 41 °C | | 0.907 | 0.048 | -1.856 | 0.342 |  |
|  |  | pupal exposure at 40 °C / adult exposure at 40 °C | | 1.042 | 0.086 | 0.498 | 0.988 |  |
|  |  | pupal exposure at 40 °C / adult exposure at 41 °C | | 1.350 | 0.133 | 3.045 | 0.020 | ^*^ |
|  |  | pupal exposure at 41 °C / adult exposure at 40 °C | | 1.149 | 0.094 | 1.702 | 0.433 |  |
|  |  | pupal exposure at 41 °C / adult exposure at 41 °C | | 1.489 | 0.146 | 4.067 | 0.001 | ^**^ |
|  |  | adult exposure at 40 °C / adult exposure at 41 °C | | 1.295 | 0.151 | 2.220 | 0.172 |  |
|  | Egg hatchability | | |  |  |  |  |  |
|  |  | control at 26 °C / pupal exposure at 40 °C | | 1.389 | 0.541 | 0.843 | 0.917 |  |
|  |  | control at 26 °C / pupal exposure at 41 °C | | 1.169 | 0.414 | 0.441 | 0.992 |  |
|  |  | control at 26 °C / adult exposure at 40 °C | | 0.980 | 0.324 | -0.061 | 1.000 |  |
|  |  | control at 26 °C / adult exposure at 41 °C | | 2.627 | 1.200 | 2.115 | 0.214 |  |
|  |  | pupal exposure at 40 °C / pupal exposure at 41 °C | | 0.842 | 0.317 | -0.457 | 0.991 |  |
|  |  | pupal exposure at 40 °C / adult exposure at 40 °C | | 0.706 | 0.250 | -0.984 | 0.863 |  |
|  |  | pupal exposure at 40 °C / adult exposure at 41 °C | | 1.892 | 0.896 | 1.346 | 0.662 |  |
|  |  | pupal exposure at 41 °C / adult exposure at 40 °C | | 0.838 | 0.264 | -0.560 | 0.981 |  |
|  |  | pupal exposure at 41 °C / adult exposure at 41 °C | | 2.247 | 1.000 | 1.815 | 0.365 |  |
|  |  | adult exposure at 40 °C / adult exposure at 41 °C | | 2.681 | 1.150 | 2.307 | 0.143 |  |
|  | Viable offspring | | |  |  |  |  |  |
|  |  | control at 26 °C / pupal exposure at 40 °C | | 1.056 | 0.166 | 0.349 | 0.997 |  |
|  |  | control at 26 °C / pupal exposure at 41 °C | | 0.944 | 0.141 | -0.386 | 0.995 |  |
|  |  | control at 26 °C / adult exposure at 40 °C | | 1.081 | 0.170 | 0.496 | 0.988 |  |
|  |  | control at 26 °C / adult exposure at 41 °C | | 1.697 | 0.283 | 3.171 | 0.013 | ^*^ |
|  |  | pupal exposure at 40 °C / pupal exposure at 41 °C | | 0.894 | 0.137 | -0.735 | 0.948 |  |
|  |  | pupal exposure at 40 °C / adult exposure at 40 °C | | 1.023 | 0.165 | 0.144 | 1.000 |  |
|  |  | pupal exposure at 40 °C / adult exposure at 41 °C | | 1.606 | 0.273 | 2.783 | 0.043 | ^*^ |
|  |  | pupal exposure at 41 °C / adult exposure at 40 °C | | 1.145 | 0.175 | 0.886 | 0.902 |  |
|  |  | pupal exposure at 41 °C / adult exposure at 41 °C | | 1.797 | 0.292 | 3.609 | 0.003 | ^**^ |
|  |  | adult exposure at 40 °C / adult exposure at 41 °C | | 1.569 | 0.267 | 2.646 | 0.062 | **^⋅^** |
| *Culex quinquefasciatus* | | |  |  |  |  |  |  |
|  | Egg laying success | | |  |  |  |  |  |
|  |  | control at 26 °C / pupal exposure at 41 °C | | 5.299 | 8.420 | 1.049 | 0.833 |  |
|  |  | control at 26 °C / pupal exposure at 42 °C | | 12.803 | 19.400 | 1.679 | 0.447 |  |
|  |  | control at 26 °C / adult exposure at 39 °C | | 1.000 | 2.040 | 0.000 | 1.000 |  |
|  |  | control at 26 °C / adult exposure at 40 °C | | 3.087 | 5.170 | 0.673 | 0.962 |  |
|  |  | pupal exposure at 41 °C / pupal exposure at 42 °C | | 2.416 | 1.970 | 1.084 | 0.815 |  |
|  |  | pupal exposure at 41 °C / adult exposure at 39 °C | | 0.189 | 0.300 | -1.049 | 0.833 |  |
|  |  | pupal exposure at 41 °C / adult exposure at 40 °C | | 0.583 | 0.627 | -0.502 | 0.987 |  |
|  |  | pupal exposure at 42 °C / adult exposure at 39 °C | | 0.078 | 0.119 | -1.679 | 0.447 |  |
|  |  | pupal exposure at 42 °C / adult exposure at 40 °C | | 0.241 | 0.233 | -1.470 | 0.582 |  |
|  |  | adult exposure at 39 °C / adult exposure at 40 °C | | 3.087 | 5.170 | 0.673 | 0.962 |  |
|  | Fecundity | | |  |  |  |  |  |
|  |  | control at 26 °C / pupal exposure at 41 °C | | 0.897 | 0.062 | -1.563 | 0.521 |  |
|  |  | control at 26 °C / pupal exposure at 42 °C | | 0.840 | 0.073 | -2.006 | 0.263 |  |
|  |  | control at 26 °C / adult exposure at 39 °C | | 0.969 | 0.065 | -0.464 | 0.991 |  |
|  |  | control at 26 °C / adult exposure at 40 °C | | 1.139 | 0.090 | 1.641 | 0.471 |  |
|  |  | pupal exposure at 41 °C / pupal exposure at 42 °C | | 0.937 | 0.079 | -0.770 | 0.939 |  |
|  |  | pupal exposure at 41 °C / adult exposure at 39 °C | | 1.081 | 0.070 | 1.207 | 0.748 |  |
|  |  | pupal exposure at 41 °C / adult exposure at 40 °C | | 1.270 | 0.098 | 3.103 | 0.016 | ^*^ |
|  |  | pupal exposure at 42 °C / adult exposure at 39 °C | | 1.154 | 0.095 | 1.729 | 0.416 |  |
|  |  | pupal exposure at 42 °C / adult exposure at 40 °C | | 1.356 | 0.126 | 3.277 | 0.009 | ^**^ |
|  |  | adult exposure at 39 °C / adult exposure at 40 °C | | 1.175 | 0.088 | 2.158 | 0.196 |  |
|  | Egg hatchability | | |  |  |  |  |  |
|  |  | control at 26 °C / pupal exposure at 41 °C | | 0.961 | 0.198 | -0.192 | 1.000 |  |
|  |  | control at 26 °C / pupal exposure at 42 °C | | 1.492 | 0.446 | 1.337 | 0.668 |  |
|  |  | control at 26 °C / adult exposure at 39 °C | | 1.347 | 0.290 | 1.383 | 0.639 |  |
|  |  | control at 26 °C / adult exposure at 40 °C | | 1.165 | 0.255 | 0.698 | 0.957 |  |
|  |  | pupal exposure at 41 °C / pupal exposure at 42 °C | | 1.552 | 0.488 | 1.398 | 0.629 |  |
|  |  | pupal exposure at 41 °C / adult exposure at 39 °C | | 1.401 | 0.331 | 1.429 | 0.609 |  |
|  |  | pupal exposure at 41 °C / adult exposure at 40 °C | | 1.212 | 0.289 | 0.803 | 0.930 |  |
|  |  | pupal exposure at 42 °C / adult exposure at 39 °C | | 0.903 | 0.290 | -0.318 | 0.998 |  |
|  |  | pupal exposure at 42 °C / adult exposure at 40 °C | | 0.781 | 0.252 | -0.767 | 0.940 |  |
|  |  | adult exposure at 39 °C / adult exposure at 40 °C | | 0.865 | 0.214 | -0.588 | 0.977 |  |
|  | Viable offspring | | |  |  |  |  |  |
|  |  | control at 26 °C / pupal exposure at 41 °C | | 0.898 | 0.077 | -1.250 | 0.722 |  |
|  |  | control at 26 °C / pupal exposure at 42 °C | | 0.828 | 0.074 | -2.120 | 0.211 |  |
|  |  | control at 26 °C / adult exposure at 39 °C | | 0.994 | 0.083 | -0.074 | 1.000 |  |
|  |  | control at 26 °C / adult exposure at 40 °C | | 1.125 | 0.096 | 1.385 | 0.637 |  |
|  |  | pupal exposure at 41 °C / pupal exposure at 42 °C | | 0.922 | 0.084 | -0.890 | 0.901 |  |
|  |  | pupal exposure at 41 °C / adult exposure at 39 °C | | 1.106 | 0.095 | 1.179 | 0.764 |  |
|  |  | pupal exposure at 41 °C / adult exposure at 40 °C | | 1.253 | 0.110 | 2.577 | 0.075 | **^⋅^** |
|  |  | pupal exposure at 42 °C / adult exposure at 39 °C | | 1.200 | 0.107 | 2.051 | 0.242 |  |
|  |  | pupal exposure at 42 °C / adult exposure at 40 °C | | 1.359 | 0.123 | 3.389 | 0.006 | ^**^ |
|  |  | adult exposure at 39 °C / adult exposure at 40 °C | | 1.132 | 0.096 | 1.458 | 0.590 |  |

Significant *p*-values are indicated with asterisks (^***^ *p* < 0.001, ^**^ *p* < 0.01, ^*^ *p* < 0.05, **^⋅^** *p* < 0.1)

2. **Sublethal heat shock effects on male fertility (*Ae. aegypti, Ae. notoscriptus* and *Cx. quinquefasciatus*)** (Table S3– S4)

Table S3. Analysis of deviance (Type II Wald Chi-square tests) of reproductive traits evaluating the effect of sublethal heat shock on male fertility

| Spp. | Trait | Factor | df | χ^2^ | *p*-value |  |
| --- | --- | --- | --- | --- | --- | --- |
| *Aedes aegypti* | |  |  |  |  |  |
|  | egg laying success | treatment | 4 | 2.983 | 0.561 |  |
|  | fecundity | treatment | 4 | 8.712 | 0.069 | **^⋅^** |
|  | egg hatchability | treatment | 4 | 23.763 | < 0.001 | ^***^ |
|  | viable offspring | treatment | 4 | 9.625 | 0.047 | ^*^ |
| *Aedes notoscriptus* | |  |  |  |  |  |
|  | egg laying success | treatment | 4 | 3.319 | 0.506 |  |
|  | fecundity | treatment | 4 | 4.700 | 0.320 |  |
|  | egg hatchability | treatment | 4 | 3.592 | 0.464 |  |
|  | viable offspring | treatment | 4 | 1.421 | 0.841 |  |
| *Culex quinquefasciatus* | |  |  |  |  |  |
|  | egg laying success | treatment | 4 | 1.450 | 0.836 |  |
|  | fecundity | treatment | 4 | 2.122 | 0.713 |  |
|  | egg hatchability | treatment | 4 | 12.044 | 0.017 | ^*^ |
|  | viable offspring | treatment | 4 | 2.559 | 0.634 |  |

Significant *p*-values are indicated with asterisks (^***^ *p* < 0.001, ^**^ *p* < 0.01, ^*^ *p* < 0.05, **^⋅^** *p* < 0.1)

Table S4. Full pairwise comparisons of reproductive traits evaluating the effects of heat shock exposure effect on male fertility

| Spp. | Trait | Comparison (treatment vs. treatment / °C) | | Ratio / OR | SE | Z-ratio | *p*-value |  |
| --- | --- | --- | --- | --- | --- | --- | --- | --- |
| *Aedes aegypti* | | |  |  |  |  |  |  |
|  | Egg laying success | | |  |  |  |  |  |
|  |  | control at 26 °C / pupal exposure at 41 °C | | 1.000 | 2.050 | 0.000 | 1.000 |  |
|  |  | control at 26 °C / pupal exposure at 42 °C | | 0.966 | 1.980 | -0.017 | 1.000 |  |
|  |  | control at 26 °C / adult exposure at 41 °C | | 1.075 | 2.210 | 0.035 | 1.000 |  |
|  |  | control at 26 °C / adult exposure at 42 °C | | 5.588 | 8.950 | 1.075 | 0.820 |  |
|  |  | pupal exposure at 41 °C / pupal exposure at 42 °C | | 0.966 | 1.980 | -0.017 | 1.000 |  |
|  |  | pupal exposure at 41 °C / adult exposure at 41 °C | | 1.075 | 2.210 | 0.035 | 1.000 |  |
|  |  | pupal exposure at 41 °C / adult exposure at 42 °C | | 5.588 | 8.950 | 1.075 | 0.820 |  |
|  |  | pupal exposure at 42 °C / adult exposure at 41 °C | | 1.113 | 2.290 | 0.052 | 1.000 |  |
|  |  | pupal exposure at 42 °C / adult exposure at 42 °C | | 5.784 | 9.250 | 1.097 | 0.808 |  |
|  |  | adult exposure at 41 °C / adult exposure at 42 °C | | 5.196 | 8.330 | 1.028 | 0.843 |  |
|  | Fecundity | | |  |  |  |  |  |
|  |  | control at 26 °C / pupal exposure at 41 °C | | 1.017 | 0.044 | 0.396 | 0.995 |  |
|  |  | control at 26 °C / pupal exposure at 42 °C | | 1.075 | 0.053 | 1.469 | 0.583 |  |
|  |  | control at 26 °C / adult exposure at 41 °C | | 1.106 | 0.057 | 1.960 | 0.286 |  |
|  |  | control at 26 °C / adult exposure at 42 °C | | 1.103 | 0.056 | 1.919 | 0.307 |  |
|  |  | pupal exposure at 41 °C / pupal exposure at 42 °C | | 1.057 | 0.040 | 1.451 | 0.595 |  |
|  |  | pupal exposure at 41 °C / adult exposure at 41 °C | | 1.087 | 0.045 | 2.048 | 0.243 |  |
|  |  | pupal exposure at 41 °C / adult exposure at 42 °C | | 1.084 | 0.044 | 2.004 | 0.264 |  |
|  |  | pupal exposure at 42 °C / adult exposure at 41 °C | | 1.029 | 0.048 | 0.610 | 0.974 |  |
|  |  | pupal exposure at 42 °C / adult exposure at 42 °C | | 1.026 | 0.048 | 0.551 | 0.982 |  |
|  |  | adult exposure at 41 °C / adult exposure at 42 °C | | 0.997 | 0.049 | -0.063 | 1.000 |  |
|  | Egg hatchability | | |  |  |  |  |  |
|  |  | control at 26 °C / pupal exposure at 41 °C | | 7.016 | 5.560 | 2.457 | 0.101 |  |
|  |  | control at 26 °C / pupal exposure at 42 °C | | 8.396 | 6.800 | 2.627 | 0.066 | **^⋅^** |
|  |  | control at 26 °C / adult exposure at 41 °C | | 1.593 | 1.360 | 0.544 | 0.983 |  |
|  |  | control at 26 °C / adult exposure at 42 °C | | 1.962 | 1.800 | 0.735 | 0.949 |  |
|  |  | pupal exposure at 41 °C / pupal exposure at 42 °C | | 1.197 | 0.392 | 0.548 | 0.982 |  |
|  |  | pupal exposure at 41 °C / adult exposure at 41 °C | | 0.227 | 0.097 | -3.457 | 0.005 | ^**^ |
|  |  | pupal exposure at 41 °C / adult exposure at 42 °C | | 0.280 | 0.151 | -2.356 | 0.128 |  |
|  |  | pupal exposure at 42 °C / adult exposure at 41 °C | | 0.190 | 0.087 | -3.618 | 0.003 | ^**^ |
|  |  | pupal exposure at 42 °C / adult exposure at 42 °C | | 0.234 | 0.132 | -2.571 | 0.076 | **^⋅^** |
|  |  | adult exposure at 41 °C / adult exposure at 42 °C | | 1.232 | 0.775 | 0.331 | 0.997 |  |
|  | Viable offspring | | |  |  |  |  |  |
|  |  | control at 26 °C / pupal exposure at 41 °C | | 1.138 | 0.069 | 2.130 | 0.207 |  |
|  |  | control at 26 °C / pupal exposure at 42 °C | | 1.199 | 0.072 | 3.003 | 0.023 | ^*^ |
|  |  | control at 26 °C / adult exposure at 41 °C | | 1.119 | 0.069 | 1.822 | 0.361 |  |
|  |  | control at 26 °C / adult exposure at 42 °C | | 1.109 | 0.070 | 1.645 | 0.468 |  |
|  |  | pupal exposure at 41 °C / pupal exposure at 42 °C | | 1.054 | 0.064 | 0.855 | 0.913 |  |
|  |  | pupal exposure at 41 °C / adult exposure at 41 °C | | 0.983 | 0.061 | -0.268 | 0.999 |  |
|  |  | pupal exposure at 41 °C / adult exposure at 42 °C | | 0.975 | 0.062 | -0.400 | 0.995 |  |
|  |  | pupal exposure at 42 °C / adult exposure at 41 °C | | 0.933 | 0.058 | -1.109 | 0.802 |  |
|  |  | pupal exposure at 42 °C / adult exposure at 42 °C | | 0.925 | 0.059 | -1.225 | 0.737 |  |
|  |  | adult exposure at 41 °C / adult exposure at 42 °C | | 0.991 | 0.064 | -0.135 | 1.000 |  |
| *Aedes notoscriptus* | | |  |  |  |  |  |  |
|  | Egg laying success | | |  |  |  |  |  |
|  |  | control at 26 °C / pupal exposure at 39 °C | | 1.562 | 1.070 | 0.651 | 0.967 |  |
|  |  | control at 26 °C / pupal exposure at 40 °C | | 1.270 | 0.895 | 0.339 | 0.997 |  |
|  |  | control at 26 °C / adult exposure at 39 °C | | 1.322 | 0.934 | 0.395 | 0.995 |  |
|  |  | control at 26 °C / adult exposure at 40 °C | | 2.869 | 1.880 | 1.606 | 0.494 |  |
|  |  | pupal exposure at 39 °C / pupal exposure at 40 °C | | 0.813 | 0.533 | -0.316 | 0.998 |  |
|  |  | pupal exposure at 39 °C / adult exposure at 39 °C | | 0.846 | 0.556 | -0.254 | 0.999 |  |
|  |  | pupal exposure at 39 °C / adult exposure at 40 °C | | 1.836 | 1.110 | 1.008 | 0.852 |  |
|  |  | pupal exposure at 40 °C / adult exposure at 39 °C | | 1.041 | 0.705 | 0.059 | 1.000 |  |
|  |  | pupal exposure at 40 °C / adult exposure at 40 °C | | 2.259 | 1.410 | 1.304 | 0.689 |  |
|  |  | adult exposure at 39 °C / adult exposure at 40 °C | | 2.170 | 1.360 | 1.237 | 0.730 |  |
|  | Fecundity | | |  |  |  |  |  |
|  |  | control at 26 °C / pupal exposure at 39 °C | | 0.870 | 0.077 | -1.579 | 0.511 |  |
|  |  | control at 26 °C / pupal exposure at 40 °C | | 0.879 | 0.088 | -1.283 | 0.702 |  |
|  |  | control at 26 °C / adult exposure at 39 °C | | 0.966 | 0.102 | -0.330 | 0.997 |  |
|  |  | control at 26 °C / adult exposure at 40 °C | | 0.978 | 0.103 | -0.208 | 1.000 |  |
|  |  | pupal exposure at 39 °C / pupal exposure at 40 °C | | 1.011 | 0.076 | 0.141 | 1.000 |  |
|  |  | pupal exposure at 39 °C / adult exposure at 39 °C | | 1.110 | 0.091 | 1.278 | 0.705 |  |
|  |  | pupal exposure at 39 °C / adult exposure at 40 °C | | 1.125 | 0.091 | 1.445 | 0.598 |  |
|  |  | pupal exposure at 40 °C / adult exposure at 39 °C | | 1.098 | 0.104 | 0.992 | 0.859 |  |
|  |  | pupal exposure at 40 °C / adult exposure at 40 °C | | 1.113 | 0.105 | 1.134 | 0.789 |  |
|  |  | adult exposure at 39 °C / adult exposure at 40 °C | | 1.013 | 0.101 | 0.130 | 1.000 |  |
|  | Egg hatchability | | |  |  |  |  |  |
|  |  | control at 26 °C / pupal exposure at 39 °C | | 0.911 | 0.341 | -0.248 | 0.999 |  |
|  |  | control at 26 °C / pupal exposure at 40 °C | | 1.496 | 0.524 | 1.150 | 0.780 |  |
|  |  | control at 26 °C / adult exposure at 39 °C | | 1.094 | 0.346 | 0.284 | 0.999 |  |
|  |  | control at 26 °C / adult exposure at 40 °C | | 0.683 | 0.333 | -0.782 | 0.936 |  |
|  |  | pupal exposure at 39 °C / pupal exposure at 40 °C | | 1.642 | 0.592 | 1.375 | 0.644 |  |
|  |  | pupal exposure at 39 °C / adult exposure at 39 °C | | 1.200 | 0.393 | 0.557 | 0.981 |  |
|  |  | pupal exposure at 39 °C / adult exposure at 40 °C | | 0.749 | 0.371 | -0.584 | 0.978 |  |
|  |  | pupal exposure at 40 °C / adult exposure at 39 °C | | 0.731 | 0.220 | -1.041 | 0.836 |  |
|  |  | pupal exposure at 40 °C / adult exposure at 40 °C | | 0.456 | 0.218 | -1.643 | 0.470 |  |
|  |  | adult exposure at 39 °C / adult exposure at 40 °C | | 0.624 | 0.283 | -1.040 | 0.837 |  |
|  | Viable offspring | | |  |  |  |  |  |
|  |  | control at 26 °C / pupal exposure at 39 °C | | 0.906 | 0.108 | -0.823 | 0.924 |  |
|  |  | control at 26 °C / pupal exposure at 40 °C | | 1.004 | 0.118 | 0.036 | 1.000 |  |
|  |  | control at 26 °C / adult exposure at 39 °C | | 1.014 | 0.137 | 0.104 | 1.000 |  |
|  |  | control at 26 °C / adult exposure at 40 °C | | 0.879 | 0.171 | -0.665 | 0.964 |  |
|  |  | pupal exposure at 39 °C / pupal exposure at 40 °C | | 1.108 | 0.132 | 0.858 | 0.912 |  |
|  |  | pupal exposure at 39 °C / adult exposure at 39 °C | | 1.119 | 0.153 | 0.822 | 0.924 |  |
|  |  | pupal exposure at 39 °C / adult exposure at 40 °C | | 0.969 | 0.190 | -0.158 | 1.000 |  |
|  |  | pupal exposure at 40 °C / adult exposure at 39 °C | | 1.010 | 0.136 | 0.073 | 1.000 |  |
|  |  | pupal exposure at 40 °C / adult exposure at 40 °C | | 0.875 | 0.170 | -0.686 | 0.960 |  |
|  |  | adult exposure at 39 °C / adult exposure at 40 °C | | 0.866 | 0.178 | -0.697 | 0.957 |  |
| *Culex quinquefasciatus* | | |  |  |  |  |  |  |
|  | Egg laying success | | |  |  |  |  |  |
|  |  | control at 26 °C / pupal exposure at 40 °C | | 1.000 | 2.040 | 0.000 | 1.000 |  |
|  |  | control at 26 °C / pupal exposure at 41 °C | | 1.030 | 2.110 | 0.014 | 1.000 |  |
|  |  | control at 26 °C / adult exposure at 38 °C | | 1.030 | 2.110 | 0.014 | 1.000 |  |
|  |  | control at 26 °C / adult exposure at 39 °C | | 3.910 | 6.560 | 0.811 | 0.927 |  |
|  |  | pupal exposure at 40 °C / pupal exposure at 41 °C | | 1.030 | 2.110 | 0.014 | 1.000 |  |
|  |  | pupal exposure at 40 °C / adult exposure at 38 °C | | 1.030 | 2.110 | 0.014 | 1.000 |  |
|  |  | pupal exposure at 40 °C / adult exposure at 39 °C | | 3.910 | 6.560 | 0.811 | 0.927 |  |
|  |  | pupal exposure at 41 °C / adult exposure at 38 °C | | 1.000 | 2.050 | 0.000 | 1.000 |  |
|  |  | pupal exposure at 41 °C / adult exposure at 39 °C | | 3.790 | 6.370 | 0.794 | 0.933 |  |
|  |  | adult exposure at 38 °C / adult exposure at 39 °C | | 3.790 | 6.370 | 0.794 | 0.933 |  |
|  | Fecundity | | |  |  |  |  |  |
|  |  | control at 26 °C / pupal exposure at 40 °C | | 1.037 | 0.063 | 0.595 | 0.976 |  |
|  |  | control at 26 °C / pupal exposure at 41 °C | | 0.980 | 0.064 | -0.311 | 0.998 |  |
|  |  | control at 26 °C / adult exposure at 38 °C | | 1.040 | 0.064 | 0.646 | 0.967 |  |
|  |  | control at 26 °C / adult exposure at 39 °C | | 1.068 | 0.062 | 1.131 | 0.790 |  |
|  |  | pupal exposure at 40 °C / pupal exposure at 41 °C | | 0.945 | 0.068 | -0.777 | 0.937 |  |
|  |  | pupal exposure at 40 °C / adult exposure at 38 °C | | 1.004 | 0.069 | 0.055 | 1.000 |  |
|  |  | pupal exposure at 40 °C / adult exposure at 39 °C | | 1.030 | 0.068 | 0.451 | 0.991 |  |
|  |  | pupal exposure at 41 °C / adult exposure at 38 °C | | 1.062 | 0.078 | 0.820 | 0.925 |  |
|  |  | pupal exposure at 41 °C / adult exposure at 39 °C | | 1.090 | 0.077 | 1.223 | 0.738 |  |
|  |  | adult exposure at 38 °C / adult exposure at 39 °C | | 1.026 | 0.069 | 0.388 | 0.995 |  |
|  | Egg hatchability | | |  |  |  |  |  |
|  |  | control at 26 °C / pupal exposure at 40 °C | | 1.593 | 0.303 | 2.449 | 0.103 |  |
|  |  | control at 26 °C / pupal exposure at 41 °C | | 1.589 | 0.343 | 2.147 | 0.200 |  |
|  |  | control at 26 °C / adult exposure at 38 °C | | 1.311 | 0.287 | 1.236 | 0.730 |  |
|  |  | control at 26 °C / adult exposure at 39 °C | | 2.015 | 0.440 | 3.209 | 0.012 | ^*^ |
|  |  | pupal exposure at 40 °C / pupal exposure at 41 °C | | 0.997 | 0.206 | -0.012 | 1.000 |  |
|  |  | pupal exposure at 40 °C / adult exposure at 38 °C | | 0.823 | 0.173 | -0.924 | 0.888 |  |
|  |  | pupal exposure at 40 °C / adult exposure at 39 °C | | 1.265 | 0.265 | 1.122 | 0.795 |  |
|  |  | pupal exposure at 41 °C / adult exposure at 38 °C | | 0.825 | 0.193 | -0.821 | 0.924 |  |
|  |  | pupal exposure at 41 °C / adult exposure at 39 °C | | 1.268 | 0.296 | 1.020 | 0.846 |  |
|  |  | adult exposure at 38 °C / adult exposure at 39 °C | | 1.537 | 0.363 | 1.819 | 0.363 |  |
|  | Viable offspring | | |  |  |  |  |  |
|  |  | control at 26 °C / pupal exposure at 40 °C | | 1.046 | 0.075 | 0.629 | 0.971 |  |
|  |  | control at 26 °C / pupal exposure at 41 °C | | 1.006 | 0.073 | 0.079 | 1.000 |  |
|  |  | control at 26 °C / adult exposure at 38 °C | | 1.055 | 0.076 | 0.745 | 0.946 |  |
|  |  | control at 26 °C / adult exposure at 39 °C | | 1.117 | 0.087 | 1.420 | 0.615 |  |
|  |  | pupal exposure at 40 °C / pupal exposure at 41 °C | | 0.961 | 0.070 | -0.540 | 0.983 |  |
|  |  | pupal exposure at 40 °C / adult exposure at 38 °C | | 1.008 | 0.073 | 0.116 | 1.000 |  |
|  |  | pupal exposure at 40 °C / adult exposure at 39 °C | | 1.068 | 0.084 | 0.833 | 0.921 |  |
|  |  | pupal exposure at 41 °C / adult exposure at 38 °C | | 1.049 | 0.077 | 0.655 | 0.966 |  |
|  |  | pupal exposure at 41 °C / adult exposure at 39 °C | | 1.111 | 0.088 | 1.329 | 0.673 |  |
|  |  | adult exposure at 38 °C / adult exposure at 39 °C | | 1.059 | 0.083 | 0.725 | 0.951 |  |

Significant *p*-values are indicated with asterisks (^***^ *p* < 0.001, ^**^ *p* < 0.01, ^*^ *p* < 0.05, **^⋅^** *p* < 0.1)

3. **Delayed effects of sublethal heat shock on *Ae. aegypti* male fertility** (Table S5– S6)

Table S5. Analysis of deviance (Type II Wald Chi-square tests) of reproductive traits evaluating the delayed effect of sublethal heat shock on *Ae. aegypti* male fertility

| Trait | Factor | df | χ^2^ | *p*-value |  |
| --- | --- | --- | --- | --- | --- |
| Egg laying success | |  |  |  |  |
|  | heat exposure | 3 | 38.417 | <0.001 | ^**^ |
|  | delayed mating time | 3 | 1.569 | 0.666 |  |
|  | interaction | 9 | 10.892 | 0.283 |  |
| Fecundity | |  |  |  |  |
|  | heat exposure | 3 | 4.179 | 0.243 |  |
|  | delayed mating time | 3 | 15.207 | 0.002 | ^**^ |
|  | interaction | 9 | 13.423 | 0.144 |  |
| Egg hatchability | |  |  |  |  |
|  | heat exposure | 3 | 11.188 | 0.011 | ^*^ |
|  | delayed mating time | 3 | 27.029 | <0.001 | ^***^ |
|  | interaction | 9 | 21.311 | 0.011 | ^*^ |
| Viable offspring | |  |  |  |  |
|  | heat exposure | 3 | 2.623 | 0.454 |  |
|  | delayed mating time | 3 | 14.638 | 0.002 | ^**^ |
|  | interaction | 9 | 9.953 | 0.354 |  |

Significant *p*-values are indicated with asterisks (^***^ *p* < 0.001, ^**^ *p* < 0.01, ^*^ *p* < 0.05, **^⋅^** *p* < 0.1)

Table S6. Full pairwise comparisons of reproductive traits evaluating the delayed effect of sublethal heat shock exposure on male fertility

| Trait | Comparison (heat exposure vs. Delayed mating time after heat exposure) | Ratio / OR | | SE | Z-ratio | *p*-value |  |
| --- | --- | --- | --- | --- | --- | --- | --- |
| Egg laying success | |  | |  |  |  |  |
|  | 1 hr at 41 °C – mated after 1 hr / 1 hr at 42 °C – mated after 1 hr | 2.209 | | 1.330 | 1.320 | 0.995 |  |
|  | 1 hr at 41 °C – mated after 1 hr / 4 hr at 38 °C – mated after 1 hr | 0.233 | | 0.226 | -1.504 | 0.982 |  |
|  | 1 hr at 41 °C – mated after 1 hr / 4 hr at 39 °C – mated after 1 hr | 0.413 | | 0.337 | -1.085 | 1.000 |  |
|  | 1 hr at 41 °C – mated after 1 hr / 1 hr at 41 °C – mated after 24 hr | 0.596 | | 0.442 | -0.699 | 1.000 |  |
|  | 1 hr at 41 °C – mated after 1 hr / 1 hr at 42 °C – mated after 24 hr | 1.940 | | 1.210 | 1.065 | 1.000 |  |
|  | 1 hr at 41 °C – mated after 1 hr / 4 hr at 38 °C – mated after 24 hr | 0.076 | | 0.115 | -1.700 | 0.947 |  |
|  | 1 hr at 41 °C – mated after 1 hr / 4 hr at 39 °C – mated after 24 hr | 0.233 | | 0.226 | -1.504 | 0.982 |  |
|  | 1 hr at 41 °C – mated after 1 hr / 1 hr at 41 °C – mated after 48 hr | 0.233 | | 0.226 | -1.504 | 0.982 |  |
|  | 1 hr at 41 °C – mated after 1 hr / 1 hr at 42 °C – mated after 48 hr | 4.055 | | 2.360 | 2.406 | 0.543 |  |
|  | 1 hr at 41 °C – mated after 1 hr / 4 hr at 38 °C – mated after 48 hr | 0.233 | | 0.226 | -1.504 | 0.982 |  |
|  | 1 hr at 41 °C – mated after 1 hr / 4 hr at 39 °C – mated after 48 hr | 0.078 | | 0.118 | -1.680 | 0.952 |  |
|  | 1 hr at 41 °C – mated after 1 hr / 1 hr at 41 °C – mated after 72 hr | 0.076 | | 0.115 | -1.700 | 0.947 |  |
|  | 1 hr at 41 °C – mated after 1 hr / 1 hr at 42 °C – mated after 72 hr | 1.658 | | 1.020 | 0.820 | 1.000 |  |
|  | 1 hr at 41 °C – mated after 1 hr / 4 hr at 38 °C – mated after 72 hr | 0.616 | | 0.457 | -0.654 | 1.000 |  |
|  | 1 hr at 41 °C – mated after 1 hr / 4 hr at 39 °C – mated after 72 hr | 0.078 | | 0.118 | -1.680 | 0.952 |  |
|  | 1 hr at 42 °C – mated after 1 hr / 4 hr at 38 °C – mated after 1 hr | 0.106 | | 0.098 | -2.433 | 0.522 |  |
|  | 1 hr at 42 °C – mated after 1 hr / 4 hr at 39 °C – mated after 1 hr | 0.187 | | 0.142 | -2.200 | 0.698 |  |
|  | 1 hr at 42 °C – mated after 1 hr / 1 hr at 41 °C – mated after 24 hr | 0.270 | | 0.184 | -1.921 | 0.867 |  |
|  | 1 hr at 42 °C – mated after 1 hr / 1 hr at 42 °C – mated after 24 hr | 0.878 | | 0.484 | -0.235 | 1.000 |  |
|  | 1 hr at 42 °C – mated after 1 hr / 4 hr at 38 °C – mated after 24 hr | 0.034 | | 0.051 | -2.263 | 0.651 |  |
|  | 1 hr at 42 °C – mated after 1 hr / 4 hr at 39 °C – mated after 24 hr | 0.106 | | 0.098 | -2.433 | 0.522 |  |
|  | 1 hr at 42 °C – mated after 1 hr / 1 hr at 41 °C – mated after 48 hr | 0.106 | | 0.098 | -2.433 | 0.522 |  |
|  | 1 hr at 42 °C – mated after 1 hr / 1 hr at 42 °C – mated after 48 hr | 1.836 | | 0.927 | 1.204 | 0.998 |  |
|  | 1 hr at 42 °C – mated after 1 hr / 4 hr at 38 °C – mated after 48 hr | 0.106 | | 0.098 | -2.433 | 0.522 |  |
|  | 1 hr at 42 °C – mated after 1 hr / 4 hr at 39 °C – mated after 48 hr | 0.035 | | 0.053 | -2.243 | 0.667 |  |
|  | 1 hr at 42 °C – mated after 1 hr / 1 hr at 41 °C – mated after 72 hr | 0.034 | | 0.051 | -2.263 | 0.651 |  |
|  | 1 hr at 42 °C – mated after 1 hr / 1 hr at 42 °C – mated after 72 hr | 0.751 | | 0.409 | -0.527 | 1.000 |  |
|  | 1 hr at 42 °C – mated after 1 hr / 4 hr at 38 °C – mated after 72 hr | 0.279 | | 0.190 | -1.871 | 0.889 |  |
|  | 1 hr at 42 °C – mated after 1 hr / 4 hr at 39 °C – mated after 72 hr | 0.035 | | 0.053 | -2.243 | 0.667 |  |
|  | 4 hr at 38 °C – mated after 1 hr / 4 hr at 39 °C – mated after 1 hr | 1.769 | | 1.900 | 0.530 | 1.000 |  |
|  | 4 hr at 38 °C – mated after 1 hr / 1 hr at 41 °C – mated after 24 hr | 2.556 | | 2.610 | 0.919 | 1.000 |  |
|  | 4 hr at 38 °C – mated after 1 hr / 1 hr at 42 °C – mated after 24 hr | 8.319 | | 7.810 | 2.257 | 0.656 |  |
|  | 4 hr at 38 °C – mated after 1 hr / 4 hr at 38 °C – mated after 24 hr | 0.324 | | 0.542 | -0.673 | 1.000 |  |
|  | 4 hr at 38 °C – mated after 1 hr / 4 hr at 39 °C – mated after 24 hr | 1.000 | | 1.200 | 0.000 | 1.000 |  |
|  | 4 hr at 38 °C – mated after 1 hr / 1 hr at 41 °C – mated after 48 hr | 1.000 | | 1.200 | 0.000 | 1.000 |  |
|  | 4 hr at 38 °C – mated after 1 hr / 1 hr at 42 °C – mated after 48 hr | 17.390 | | 15.900 | 3.131 | 0.120 |  |
|  | 4 hr at 38 °C – mated after 1 hr / 4 hr at 38 °C – mated after 48 hr | 1.000 | | 1.200 | 0.000 | 1.000 |  |
|  | 4 hr at 38 °C – mated after 1 hr / 4 hr at 39 °C – mated after 48 hr | 0.333 | | 0.558 | -0.656 | 1.000 |  |
|  | 4 hr at 38 °C – mated after 1 hr / 1 hr at 41 °C – mated after 72 hr | 0.324 | | 0.542 | -0.673 | 1.000 |  |
|  | 4 hr at 38 °C – mated after 1 hr / 1 hr at 42 °C – mated after 72 hr | 7.109 | | 6.650 | 2.098 | 0.767 |  |
|  | 4 hr at 38 °C – mated after 1 hr / 4 hr at 38 °C – mated after 72 hr | 2.639 | | 2.700 | 0.950 | 1.000 |  |
|  | 4 hr at 38 °C – mated after 1 hr / 4 hr at 39 °C – mated after 72 hr | 0.333 | | 0.558 | -0.656 | 1.000 |  |
|  | 4 hr at 39 °C – mated after 1 hr / 1 hr at 41 °C – mated after 24 hr | 1.444 | | 1.270 | 0.419 | 1.000 |  |
|  | 4 hr at 39 °C – mated after 1 hr / 1 hr at 42 °C – mated after 24 hr | 4.702 | | 3.670 | 1.984 | 0.835 |  |
|  | 4 hr at 39 °C – mated after 1 hr / 4 hr at 38 °C – mated after 24 hr | 0.183 | | 0.291 | -1.067 | 1.000 |  |
|  | 4 hr at 39 °C – mated after 1 hr / 4 hr at 39 °C – mated after 24 hr | 0.565 | | 0.608 | -0.530 | 1.000 |  |
|  | 4 hr at 39 °C – mated after 1 hr / 1 hr at 41 °C – mated after 48 hr | 0.565 | | 0.608 | -0.530 | 1.000 |  |
|  | 4 hr at 39 °C – mated after 1 hr / 1 hr at 42 °C – mated after 48 hr | 9.829 | | 7.360 | 3.054 | 0.147 |  |
|  | 4 hr at 39 °C – mated after 1 hr / 4 hr at 38 °C – mated after 48 hr | 0.565 | | 0.608 | -0.530 | 1.000 |  |
|  | 4 hr at 39 °C – mated after 1 hr / 4 hr at 39 °C – mated after 48 hr | 0.188 | | 0.300 | -1.049 | 1.000 |  |
|  | 4 hr at 39 °C – mated after 1 hr / 1 hr at 41 °C – mated after 72 hr | 0.183 | | 0.291 | -1.067 | 1.000 |  |
|  | 4 hr at 39 °C – mated after 1 hr / 1 hr at 42 °C – mated after 72 hr | 4.018 | | 3.120 | 1.793 | 0.919 |  |
|  | 4 hr at 39 °C – mated after 1 hr / 4 hr at 38 °C – mated after 72 hr | 1.492 | | 1.310 | 0.455 | 1.000 |  |
|  | 4 hr at 39 °C – mated after 1 hr / 4 hr at 39 °C – mated after 72 hr | 0.188 | | 0.300 | -1.049 | 1.000 |  |
|  | 1 hr at 41 °C – mated after 24 hr / 1 hr at 42 °C – mated after 24 hr | | 3.255 | 2.280 | 1.682 | 0.952 |  |
|  | 1 hr at 41 °C – mated after 24 hr / 4 hr at 38 °C – mated after 24 hr | | 0.127 | 0.197 | -1.330 | 0.995 |  |
|  | 1 hr at 41 °C – mated after 24 hr / 4 hr at 39 °C – mated after 24 hr | | 0.391 | 0.400 | -0.919 | 1.000 |  |
|  | 1 hr at 41 °C – mated after 24 hr / 1 hr at 41 °C – mated after 48 hr | | 0.391 | 0.400 | -0.919 | 1.000 |  |
|  | 1 hr at 41 °C – mated after 24 hr / 1 hr at 42 °C – mated after 48 hr | | 6.805 | 4.530 | 2.880 | 0.225 |  |
|  | 1 hr at 41 °C – mated after 24 hr / 4 hr at 38 °C – mated after 48 hr | | 0.391 | 0.400 | -0.919 | 1.000 |  |
|  | 1 hr at 41 °C – mated after 24 hr / 4 hr at 39 °C – mated after 48 hr | | 0.130 | 0.203 | -1.311 | 0.996 |  |
|  | 1 hr at 41 °C – mated after 24 hr / 1 hr at 41 °C – mated after 72 hr | | 0.127 | 0.197 | -1.330 | 0.995 |  |
|  | 1 hr at 41 °C – mated after 24 hr / 1 hr at 42 °C – mated after 72 hr | | 2.782 | 1.940 | 1.469 | 0.986 |  |
|  | 1 hr at 41 °C – mated after 24 hr / 4 hr at 38 °C – mated after 72 hr | | 1.033 | 0.836 | 0.040 | 1.000 |  |
|  | 1 hr at 41 °C – mated after 24 hr / 4 hr at 39 °C – mated after 72 hr | | 0.130 | 0.203 | -1.311 | 0.996 |  |
|  | 1 hr at 42 °C – mated after 24 hr / 4 hr at 38 °C – mated after 24 hr | | 0.039 | 0.058 | -2.163 | 0.723 |  |
|  | 1 hr at 42 °C – mated after 24 hr / 4 hr at 39 °C – mated after 24 hr | | 0.120 | 0.113 | -2.257 | 0.656 |  |
|  | 1 hr at 42 °C – mated after 24 hr / 1 hr at 41 °C – mated after 48 hr | | 0.120 | 0.113 | -2.257 | 0.656 |  |
|  | 1 hr at 42 °C – mated after 24 hr / 1 hr at 42 °C – mated after 48 hr | | 2.090 | 1.110 | 1.389 | 0.992 |  |
|  | 1 hr at 42 °C – mated after 24 hr / 4 hr at 38 °C – mated after 48 hr | | 0.120 | 0.113 | -2.257 | 0.656 |  |
|  | 1 hr at 42 °C – mated after 24 hr / 4 hr at 39 °C – mated after 48 hr | | 0.040 | 0.060 | -2.143 | 0.737 |  |
|  | 1 hr at 42 °C – mated after 24 hr / 1 hr at 41 °C – mated after 72 hr | | 0.039 | 0.058 | -2.163 | 0.723 |  |
|  | 1 hr at 42 °C – mated after 24 hr / 1 hr at 42 °C – mated after 72 hr | | 0.855 | 0.486 | -0.276 | 1.000 |  |
|  | 1 hr at 42 °C – mated after 24 hr / 4 hr at 38 °C – mated after 72 hr | | 0.317 | 0.223 | -1.634 | 0.962 |  |
|  | 1 hr at 42 °C – mated after 24 hr / 4 hr at 39 °C – mated after 72 hr | | 0.040 | 0.060 | -2.143 | 0.737 |  |
|  | 4 hr at 38 °C – mated after 24 hr / 4 hr at 39 °C – mated after 24 hr | | 3.087 | 5.170 | 0.673 | 1.000 |  |
|  | 4 hr at 38 °C – mated after 24 hr / 1 hr at 41 °C – mated after 48 hr | | 3.087 | 5.170 | 0.673 | 1.000 |  |
|  | 4 hr at 38 °C – mated after 24 hr / 1 hr at 42 °C – mated after 48 hr | | 53.683 | 79.700 | 2.684 | 0.341 |  |
|  | 4 hr at 38 °C – mated after 24 hr / 4 hr at 38 °C – mated after 48 hr | | 3.087 | 5.170 | 0.673 | 1.000 |  |
|  | 4 hr at 38 °C – mated after 24 hr / 4 hr at 39 °C – mated after 48 hr | | 1.029 | 2.100 | 0.014 | 1.000 |  |
|  | 4 hr at 38 °C – mated after 24 hr / 1 hr at 41 °C – mated after 72 hr | | 1.000 | 2.040 | 0.000 | 1.000 |  |
|  | 4 hr at 38 °C – mated after 24 hr / 1 hr at 42 °C – mated after 72 hr | | 21.946 | 32.900 | 2.062 | 0.790 |  |
|  | 4 hr at 38 °C – mated after 24 hr / 4 hr at 38 °C – mated after 72 hr | | 8.148 | 12.700 | 1.350 | 0.994 |  |
|  | 4 hr at 38 °C – mated after 24 hr / 4 hr at 39 °C – mated after 72 hr | | 1.029 | 2.100 | 0.014 | 1.000 |  |
|  | 4 hr at 39 °C – mated after 24 hr / 1 hr at 41 °C – mated after 48 hr | | 1.000 | 1.200 | 0.000 | 1.000 |  |
|  | 4 hr at 39 °C – mated after 24 hr / 1 hr at 42 °C – mated after 48 hr | | 17.390 | 15.900 | 3.131 | 0.120 |  |
|  | 4 hr at 39 °C – mated after 24 hr / 4 hr at 38 °C – mated after 48 hr | | 1.000 | 1.200 | 0.000 | 1.000 |  |
|  | 4 hr at 39 °C – mated after 24 hr / 4 hr at 39 °C – mated after 48 hr | | 0.333 | 0.558 | -0.656 | 1.000 |  |
|  | 4 hr at 39 °C – mated after 24 hr / 1 hr at 41 °C – mated after 72 hr | | 0.324 | 0.542 | -0.673 | 1.000 |  |
|  | 4 hr at 39 °C – mated after 24 hr / 1 hr at 42 °C – mated after 72 hr | | 7.109 | 6.650 | 2.098 | 0.767 |  |
|  | 4 hr at 39 °C – mated after 24 hr / 4 hr at 38 °C – mated after 72 hr | | 2.639 | 2.700 | 0.950 | 1.000 |  |
|  | 4 hr at 39 °C – mated after 24 hr / 4 hr at 39 °C – mated after 72 hr | | 0.333 | 0.558 | -0.656 | 1.000 |  |
|  | 1 hr at 41 °C – mated after 48 hr / 1 hr at 42 °C – mated after 48 hr | | 17.390 | 15.900 | 3.131 | 0.120 |  |
|  | 1 hr at 41 °C – mated after 48 hr / 4 hr at 38 °C – mated after 48 hr | | 1.000 | 1.200 | 0.000 | 1.000 |  |
|  | 1 hr at 41 °C – mated after 48 hr / 4 hr at 39 °C – mated after 48 hr | | 0.333 | 0.558 | -0.656 | 1.000 |  |
|  | 1 hr at 41 °C – mated after 48 hr / 1 hr at 41 °C – mated after 72 hr | | 0.324 | 0.542 | -0.673 | 1.000 |  |
|  | 1 hr at 41 °C – mated after 48 hr / 1 hr at 42 °C – mated after 72 hr | | 7.109 | 6.650 | 2.098 | 0.767 |  |
|  | 1 hr at 41 °C – mated after 48 hr / 4 hr at 38 °C – mated after 72 hr | | 2.639 | 2.700 | 0.950 | 1.000 |  |
|  | 1 hr at 41 °C – mated after 48 hr / 4 hr at 39 °C – mated after 72 hr | | 0.333 | 0.558 | -0.656 | 1.000 |  |
|  | 1 hr at 42 °C – mated after 48 hr / 4 hr at 38 °C – mated after 48 hr | | 0.058 | 0.052 | -3.131 | 0.120 |  |
|  | 1 hr at 42 °C – mated after 48 hr / 4 hr at 39 °C – mated after 48 hr | | 0.019 | 0.029 | -2.663 | 0.355 |  |
|  | 1 hr at 42 °C – mated after 48 hr / 1 hr at 41 °C – mated after 72 hr | | 0.019 | 0.028 | -2.684 | 0.341 |  |
|  | 1 hr at 42 °C – mated after 48 hr / 1 hr at 42 °C – mated after 72 hr | | 0.409 | 0.214 | -1.706 | 0.946 |  |
|  | 1 hr at 42 °C – mated after 48 hr / 4 hr at 38 °C – mated after 72 hr | | 0.152 | 0.101 | -2.827 | 0.253 |  |
|  | 1 hr at 42 °C – mated after 48 hr / 4 hr at 39 °C – mated after 72 hr | | 0.019 | 0.029 | -2.663 | 0.355 |  |
|  | 4 hr at 38 °C – mated after 48 hr / 4 hr at 39 °C – mated after 48 hr | | 0.333 | 0.558 | -0.656 | 1.000 |  |
|  | 4 hr at 38 °C – mated after 48 hr / 1 hr at 41 °C – mated after 72 hr | | 0.324 | 0.542 | -0.673 | 1.000 |  |
|  | 4 hr at 38 °C – mated after 48 hr / 1 hr at 42 °C – mated after 72 hr | | 7.109 | 6.650 | 2.098 | 0.767 |  |
|  | 4 hr at 38 °C – mated after 48 hr / 4 hr at 38 °C – mated after 72 hr | | 2.639 | 2.700 | 0.950 | 1.000 |  |
|  | 4 hr at 38 °C – mated after 48 hr / 4 hr at 39 °C – mated after 72 hr | | 0.333 | 0.558 | -0.656 | 1.000 |  |
|  | 4 hr at 39 °C – mated after 48 hr / 1 hr at 41 °C – mated after 72 hr | | 0.972 | 1.990 | -0.014 | 1.000 |  |
|  | 4 hr at 39 °C – mated after 48 hr / 1 hr at 42 °C – mated after 72 hr | | 21.327 | 32.000 | 2.041 | 0.802 |  |
|  | 4 hr at 39 °C – mated after 48 hr / 4 hr at 38 °C – mated after 72 hr | | 7.918 | 12.300 | 1.331 | 0.995 |  |
|  | 4 hr at 39 °C – mated after 48 hr / 4 hr at 39 °C – mated after 72 hr | | 1.000 | 2.040 | 0.000 | 1.000 |  |
|  | 1 hr at 41 °C – mated after 72 hr / 1 hr at 42 °C – mated after 72 hr | | 21.946 | 32.900 | 2.062 | 0.790 |  |
|  | 1 hr at 41 °C – mated after 72 hr / 4 hr at 38 °C – mated after 72 hr | | 8.148 | 12.700 | 1.350 | 0.994 |  |
|  | 1 hr at 41 °C – mated after 72 hr / 4 hr at 39 °C – mated after 72 hr | | 1.029 | 2.100 | 0.014 | 1.000 |  |
|  | 1 hr at 42 °C – mated after 72 hr / 4 hr at 38 °C – mated after 72 hr | | 0.371 | 0.259 | -1.420 | 0.990 |  |
|  | 1 hr at 42 °C – mated after 72 hr / 4 hr at 39 °C – mated after 72 hr | | 0.047 | 0.070 | -2.041 | 0.802 |  |
|  | 4 hr at 38 °C – mated after 72 hr / 4 hr at 39 °C – mated after 72 hr | | 0.126 | 0.196 | -1.331 | 0.995 |  |
| Fecundity | |  | |  |  |  |  |
|  | 1 hr at 41 °C – mated after 1 hr / 1 hr at 42 °C – mated after 1 hr | 1.067 | | 0.083 | 0.828 | 1.000 |  |
|  | 1 hr at 41 °C – mated after 1 hr / 4 hr at 38 °C – mated after 1 hr | 0.993 | | 0.080 | -0.090 | 1.000 |  |
|  | 1 hr at 41 °C – mated after 1 hr / 4 hr at 39 °C – mated after 1 hr | 1.050 | | 0.076 | 0.675 | 1.000 |  |
|  | 1 hr at 41 °C – mated after 1 hr / 1 hr at 41 °C – mated after 24 hr | 0.921 | | 0.070 | -1.086 | 1.000 |  |
|  | 1 hr at 41 °C – mated after 1 hr / 1 hr at 42 °C – mated after 24 hr | 1.046 | | 0.119 | 0.391 | 1.000 |  |
|  | 1 hr at 41 °C – mated after 1 hr / 4 hr at 38 °C – mated after 24 hr | 0.936 | | 0.065 | -0.948 | 1.000 |  |
|  | 1 hr at 41 °C – mated after 1 hr / 4 hr at 39 °C – mated after 24 hr | 1.044 | | 0.089 | 0.500 | 1.000 |  |
|  | 1 hr at 41 °C – mated after 1 hr / 1 hr at 41 °C – mated after 48 hr | 0.910 | | 0.059 | -1.473 | 0.986 |  |
|  | 1 hr at 41 °C – mated after 1 hr / 1 hr at 42 °C – mated after 48 hr | 0.847 | | 0.058 | -2.407 | 0.542 |  |
|  | 1 hr at 41 °C – mated after 1 hr / 4 hr at 38 °C – mated after 48 hr | 0.867 | | 0.064 | -1.947 | 0.854 |  |
|  | 1 hr at 41 °C – mated after 1 hr / 4 hr at 39 °C – mated after 48 hr | 0.964 | | 0.079 | -0.447 | 1.000 |  |
|  | 1 hr at 41 °C – mated after 1 hr / 1 hr at 41 °C – mated after 72 hr | 0.854 | | 0.054 | -2.478 | 0.488 |  |
|  | 1 hr at 41 °C – mated after 1 hr / 1 hr at 42 °C – mated after 72 hr | 0.976 | | 0.092 | -0.254 | 1.000 |  |
|  | 1 hr at 41 °C – mated after 1 hr / 4 hr at 38 °C – mated after 72 hr | 1.006 | | 0.071 | 0.088 | 1.000 |  |
|  | 1 hr at 41 °C – mated after 1 hr / 4 hr at 39 °C – mated after 72 hr | 0.916 | | 0.073 | -1.094 | 0.999 |  |
|  | 1 hr at 42 °C – mated after 1 hr / 4 hr at 38 °C – mated after 1 hr | 0.931 | | 0.076 | -0.876 | 1.000 |  |
|  | 1 hr at 42 °C – mated after 1 hr / 4 hr at 39 °C – mated after 1 hr | 0.985 | | 0.073 | -0.208 | 1.000 |  |
|  | 1 hr at 42 °C – mated after 1 hr / 1 hr at 41 °C – mated after 24 hr | 0.864 | | 0.066 | -1.906 | 0.874 |  |
|  | 1 hr at 42 °C – mated after 1 hr / 1 hr at 42 °C – mated after 24 hr | 0.980 | | 0.113 | -0.174 | 1.000 |  |
|  | 1 hr at 42 °C – mated after 1 hr / 4 hr at 38 °C – mated after 24 hr | 0.878 | | 0.062 | -1.835 | 0.904 |  |
|  | 1 hr at 42 °C – mated after 1 hr / 4 hr at 39 °C – mated after 24 hr | 0.979 | | 0.085 | -0.250 | 1.000 |  |
|  | 1 hr at 42 °C – mated after 1 hr / 1 hr at 41 °C – mated after 48 hr | 0.853 | | 0.056 | -2.417 | 0.535 |  |
|  | 1 hr at 42 °C – mated after 1 hr / 1 hr at 42 °C – mated after 48 hr | 0.794 | | 0.056 | -3.274 | 0.080 | **^⋅^** |
|  | 1 hr at 42 °C – mated after 1 hr / 4 hr at 38 °C – mated after 48 hr | 0.813 | | 0.061 | -2.775 | 0.283 |  |
|  | 1 hr at 42 °C – mated after 1 hr / 4 hr at 39 °C – mated after 48 hr | 0.904 | | 0.075 | -1.217 | 0.998 |  |
|  | 1 hr at 42 °C – mated after 1 hr / 1 hr at 41 °C – mated after 72 hr | 0.801 | | 0.052 | -3.406 | 0.053 | ^*^ |
|  | 1 hr at 42 °C – mated after 1 hr / 1 hr at 42 °C – mated after 72 hr | 0.915 | | 0.087 | -0.926 | 1.000 |  |
|  | 1 hr at 42 °C – mated after 1 hr / 4 hr at 38 °C – mated after 72 hr | 0.943 | | 0.068 | -0.812 | 1.000 |  |
|  | 1 hr at 42 °C – mated after 1 hr / 4 hr at 39 °C – mated after 72 hr | 0.859 | | 0.070 | -1.869 | 0.890 |  |
|  | 4 hr at 38 °C – mated after 1 hr / 4 hr at 39 °C – mated after 1 hr | 1.058 | | 0.081 | 0.733 | 1.000 |  |
|  | 4 hr at 38 °C – mated after 1 hr / 1 hr at 41 °C – mated after 24 hr | 0.928 | | 0.074 | -0.939 | 1.000 |  |
|  | 4 hr at 38 °C – mated after 1 hr / 1 hr at 42 °C – mated after 24 hr | 1.053 | | 0.123 | 0.444 | 1.000 |  |
|  | 4 hr at 38 °C – mated after 1 hr / 4 hr at 38 °C – mated after 24 hr | 0.943 | | 0.070 | -0.793 | 1.000 |  |
|  | 4 hr at 38 °C – mated after 1 hr / 4 hr at 39 °C – mated after 24 hr | 1.051 | | 0.094 | 0.561 | 1.000 |  |
|  | 4 hr at 38 °C – mated after 1 hr / 1 hr at 41 °C – mated after 48 hr | 0.916 | | 0.063 | -1.266 | 0.997 |  |
|  | 4 hr at 38 °C – mated after 1 hr / 1 hr at 42 °C – mated after 48 hr | 0.854 | | 0.063 | -2.161 | 0.725 |  |
|  | 4 hr at 38 °C – mated after 1 hr / 4 hr at 38 °C – mated after 48 hr | 0.873 | | 0.068 | -1.747 | 0.934 |  |
|  | 4 hr at 38 °C – mated after 1 hr / 4 hr at 39 °C – mated after 48 hr | 0.971 | | 0.083 | -0.343 | 1.000 |  |
|  | 4 hr at 38 °C – mated after 1 hr / 1 hr at 41 °C – mated after 72 hr | 0.860 | | 0.059 | -2.198 | 0.699 |  |
|  | 4 hr at 38 °C – mated after 1 hr / 1 hr at 42 °C – mated after 72 hr | 0.983 | | 0.096 | -0.171 | 1.000 |  |
|  | 4 hr at 38 °C – mated after 1 hr / 4 hr at 38 °C – mated after 72 hr | 1.014 | | 0.076 | 0.180 | 1.000 |  |
|  | 4 hr at 38 °C – mated after 1 hr / 4 hr at 39 °C – mated after 72 hr | 0.923 | | 0.078 | -0.957 | 1.000 |  |
|  | 4 hr at 39 °C – mated after 1 hr / 1 hr at 41 °C – mated after 24 hr | 0.877 | | 0.063 | -1.833 | 0.905 |  |
|  | 4 hr at 39 °C – mated after 1 hr / 1 hr at 42 °C – mated after 24 hr | 0.996 | | 0.111 | -0.040 | 1.000 |  |
|  | 4 hr at 39 °C – mated after 1 hr / 4 hr at 38 °C – mated after 24 hr | 0.891 | | 0.058 | -1.763 | 0.929 |  |
|  | 4 hr at 39 °C – mated after 1 hr / 4 hr at 39 °C – mated after 24 hr | 0.994 | | 0.082 | -0.076 | 1.000 |  |
|  | 4 hr at 39 °C – mated after 1 hr / 1 hr at 41 °C – mated after 48 hr | 0.866 | | 0.052 | -2.415 | 0.536 |  |
|  | 4 hr at 39 °C – mated after 1 hr / 1 hr at 42 °C – mated after 48 hr | 0.807 | | 0.052 | -3.334 | 0.067 | **^⋅^** |
|  | 4 hr at 39 °C – mated after 1 hr / 4 hr at 38 °C – mated after 48 hr | 0.826 | | 0.057 | -2.774 | 0.284 |  |
|  | 4 hr at 39 °C – mated after 1 hr / 4 hr at 39 °C – mated after 48 hr | 0.918 | | 0.072 | -1.096 | 0.999 |  |
|  | 4 hr at 39 °C – mated after 1 hr / 1 hr at 41 °C – mated after 72 hr | 0.813 | | 0.048 | -3.515 | 0.038 | ^*^ |
|  | 4 hr at 39 °C – mated after 1 hr / 1 hr at 42 °C – mated after 72 hr | 0.930 | | 0.085 | -0.801 | 1.000 |  |
|  | 4 hr at 39 °C – mated after 1 hr / 4 hr at 38 °C – mated after 72 hr | 0.958 | | 0.063 | -0.649 | 1.000 |  |
|  | 4 hr at 39 °C – mated after 1 hr / 4 hr at 39 °C – mated after 72 hr | 0.872 | | 0.067 | -1.790 | 0.920 |  |
|  | 1 hr at 41 °C – mated after 24 hr / 1 hr at 42 °C – mated after 24 hr | | 1.135 | 0.128 | 1.118 | 0.999 |  |
|  | 1 hr at 41 °C – mated after 24 hr / 4 hr at 38 °C – mated after 24 hr | | 1.016 | 0.070 | 0.234 | 1.000 |  |
|  | 1 hr at 41 °C – mated after 24 hr / 4 hr at 39 °C – mated after 24 hr | | 1.133 | 0.096 | 1.474 | 0.986 |  |
|  | 1 hr at 41 °C – mated after 24 hr / 1 hr at 41 °C – mated after 48 hr | | 0.987 | 0.062 | -0.201 | 1.000 |  |
|  | 1 hr at 41 °C – mated after 24 hr / 1 hr at 42 °C – mated after 48 hr | | 0.920 | 0.062 | -1.237 | 0.998 |  |
|  | 1 hr at 41 °C – mated after 24 hr / 4 hr at 38 °C – mated after 48 hr | | 0.941 | 0.068 | -0.840 | 1.000 |  |
|  | 1 hr at 41 °C – mated after 24 hr / 4 hr at 39 °C – mated after 48 hr | | 1.046 | 0.085 | 0.562 | 1.000 |  |
|  | 1 hr at 41 °C – mated after 24 hr / 1 hr at 41 °C – mated after 72 hr | | 0.927 | 0.058 | -1.214 | 0.998 |  |
|  | 1 hr at 41 °C – mated after 24 hr / 1 hr at 42 °C – mated after 72 hr | | 1.060 | 0.099 | 0.622 | 1.000 |  |
|  | 1 hr at 41 °C – mated after 24 hr / 4 hr at 38 °C – mated after 72 hr | | 1.092 | 0.076 | 1.274 | 0.997 |  |
|  | 1 hr at 41 °C – mated after 24 hr / 4 hr at 39 °C – mated after 72 hr | | 0.994 | 0.079 | -0.072 | 1.000 |  |
|  | 1 hr at 42 °C – mated after 24 hr / 4 hr at 38 °C – mated after 24 hr | | 0.895 | 0.098 | -1.011 | 1.000 |  |
|  | 1 hr at 42 °C – mated after 24 hr / 4 hr at 39 °C – mated after 24 hr | | 0.998 | 0.120 | -0.015 | 1.000 |  |
|  | 1 hr at 42 °C – mated after 24 hr / 1 hr at 41 °C – mated after 48 hr | | 0.870 | 0.092 | -1.313 | 0.996 |  |
|  | 1 hr at 42 °C – mated after 24 hr / 1 hr at 42 °C – mated after 48 hr | | 0.810 | 0.088 | -1.931 | 0.862 |  |
|  | 1 hr at 42 °C – mated after 24 hr / 4 hr at 38 °C – mated after 48 hr | | 0.829 | 0.093 | -1.676 | 0.953 |  |
|  | 1 hr at 42 °C – mated after 24 hr / 4 hr at 39 °C – mated after 48 hr | | 0.922 | 0.108 | -0.691 | 1.000 |  |
|  | 1 hr at 42 °C – mated after 24 hr / 1 hr at 41 °C – mated after 72 hr | | 0.817 | 0.086 | -1.914 | 0.870 |  |
|  | 1 hr at 42 °C – mated after 24 hr / 1 hr at 42 °C – mated after 72 hr | | 0.934 | 0.118 | -0.541 | 1.000 |  |
|  | 1 hr at 42 °C – mated after 24 hr / 4 hr at 38 °C – mated after 72 hr | | 0.962 | 0.106 | -0.349 | 1.000 |  |
|  | 1 hr at 42 °C – mated after 24 hr / 4 hr at 39 °C – mated after 72 hr | | 0.876 | 0.102 | -1.137 | 0.999 |  |
|  | 4 hr at 38 °C – mated after 24 hr / 4 hr at 39 °C – mated after 24 hr | | 1.115 | 0.089 | 1.369 | 0.993 |  |
|  | 4 hr at 38 °C – mated after 24 hr / 1 hr at 41 °C – mated after 48 hr | | 0.972 | 0.054 | -0.513 | 1.000 |  |
|  | 4 hr at 38 °C – mated after 24 hr / 1 hr at 42 °C – mated after 48 hr | | 0.905 | 0.055 | -1.633 | 0.963 |  |
|  | 4 hr at 38 °C – mated after 24 hr / 4 hr at 38 °C – mated after 48 hr | | 0.926 | 0.061 | -1.161 | 0.999 |  |
|  | 4 hr at 38 °C – mated after 24 hr / 4 hr at 39 °C – mated after 48 hr | | 1.030 | 0.078 | 0.390 | 1.000 |  |
|  | 4 hr at 38 °C – mated after 24 hr / 1 hr at 41 °C – mated after 72 hr | | 0.912 | 0.050 | -1.664 | 0.956 |  |
|  | 4 hr at 38 °C – mated after 24 hr / 1 hr at 42 °C – mated after 72 hr | | 1.043 | 0.093 | 0.474 | 1.000 |  |
|  | 4 hr at 38 °C – mated after 24 hr / 4 hr at 38 °C – mated after 72 hr | | 1.075 | 0.068 | 1.150 | 0.999 |  |
|  | 4 hr at 38 °C – mated after 24 hr / 4 hr at 39 °C – mated after 72 hr | | 0.979 | 0.072 | -0.295 | 1.000 |  |
|  | 4 hr at 39 °C – mated after 24 hr / 1 hr at 41 °C – mated after 48 hr | | 0.872 | 0.065 | -1.836 | 0.904 |  |
|  | 4 hr at 39 °C – mated after 24 hr / 1 hr at 42 °C – mated after 48 hr | | 0.812 | 0.064 | -2.645 | 0.367 |  |
|  | 4 hr at 39 °C – mated after 24 hr / 4 hr at 38 °C – mated after 48 hr | | 0.831 | 0.069 | -2.242 | 0.667 |  |
|  | 4 hr at 39 °C – mated after 24 hr / 4 hr at 39 °C – mated after 48 hr | | 0.924 | 0.084 | -0.879 | 1.000 |  |
|  | 4 hr at 39 °C – mated after 24 hr / 1 hr at 41 °C – mated after 72 hr | | 0.818 | 0.061 | -2.698 | 0.331 |  |
|  | 4 hr at 39 °C – mated after 24 hr / 1 hr at 42 °C – mated after 72 hr | | 0.935 | 0.095 | -0.655 | 1.000 |  |
|  | 4 hr at 39 °C – mated after 24 hr / 4 hr at 38 °C – mated after 72 hr | | 0.964 | 0.077 | -0.457 | 1.000 |  |
|  | 4 hr at 39 °C – mated after 24 hr / 4 hr at 39 °C – mated after 72 hr | | 0.878 | 0.078 | -1.469 | 0.986 |  |
|  | 1 hr at 41 °C – mated after 48 hr / 1 hr at 42 °C – mated after 48 hr | | 0.931 | 0.051 | -1.295 | 0.996 |  |
|  | 1 hr at 41 °C – mated after 48 hr / 4 hr at 38 °C – mated after 48 hr | | 0.953 | 0.057 | -0.796 | 1.000 |  |
|  | 1 hr at 41 °C – mated after 48 hr / 4 hr at 39 °C – mated after 48 hr | | 1.060 | 0.075 | 0.824 | 1.000 |  |
|  | 1 hr at 41 °C – mated after 48 hr / 1 hr at 41 °C – mated after 72 hr | | 0.939 | 0.045 | -1.310 | 0.996 |  |
|  | 1 hr at 41 °C – mated after 48 hr / 1 hr at 42 °C – mated after 72 hr | | 1.073 | 0.091 | 0.835 | 1.000 |  |
|  | 1 hr at 41 °C – mated after 48 hr / 4 hr at 38 °C – mated after 72 hr | | 1.106 | 0.063 | 1.776 | 0.925 |  |
|  | 1 hr at 41 °C – mated after 48 hr / 4 hr at 39 °C – mated after 72 hr | | 1.007 | 0.069 | 0.101 | 1.000 |  |
|  | 1 hr at 42 °C – mated after 48 hr / 4 hr at 38 °C – mated after 48 hr | | 1.023 | 0.067 | 0.354 | 1.000 |  |
|  | 1 hr at 42 °C – mated after 48 hr / 4 hr at 39 °C – mated after 48 hr | | 1.138 | 0.085 | 1.729 | 0.940 |  |
|  | 1 hr at 42 °C – mated after 48 hr / 1 hr at 41 °C – mated after 72 hr | | 1.008 | 0.055 | 0.147 | 1.000 |  |
|  | 1 hr at 42 °C – mated after 48 hr / 1 hr at 42 °C – mated after 72 hr | | 1.152 | 0.102 | 1.607 | 0.967 |  |
|  | 1 hr at 42 °C – mated after 48 hr / 4 hr at 38 °C – mated after 72 hr | | 1.187 | 0.073 | 2.777 | 0.282 |  |
|  | 1 hr at 42 °C – mated after 48 hr / 4 hr at 39 °C – mated after 72 hr | | 1.081 | 0.079 | 1.070 | 1.000 |  |
|  | 4 hr at 38 °C – mated after 48 hr / 4 hr at 39 °C – mated after 48 hr | | 1.112 | 0.088 | 1.346 | 0.994 |  |
|  | 4 hr at 38 °C – mated after 48 hr / 1 hr at 41 °C – mated after 72 hr | | 0.985 | 0.059 | -0.254 | 1.000 |  |
|  | 4 hr at 38 °C – mated after 48 hr / 1 hr at 42 °C – mated after 72 hr | | 1.126 | 0.103 | 1.295 | 0.996 |  |
|  | 4 hr at 38 °C – mated after 48 hr / 4 hr at 38 °C – mated after 72 hr | | 1.160 | 0.077 | 2.229 | 0.676 |  |
|  | 4 hr at 38 °C – mated after 48 hr / 4 hr at 39 °C – mated after 72 hr | | 1.056 | 0.081 | 0.713 | 1.000 |  |
|  | 4 hr at 39 °C – mated after 48 hr / 1 hr at 41 °C – mated after 72 hr | | 0.886 | 0.062 | -1.733 | 0.938 |  |
|  | 4 hr at 39 °C – mated after 48 hr / 1 hr at 42 °C – mated after 72 hr | | 1.013 | 0.100 | 0.129 | 1.000 |  |
|  | 4 hr at 39 °C – mated after 48 hr / 4 hr at 38 °C – mated after 72 hr | | 1.044 | 0.079 | 0.562 | 1.000 |  |
|  | 4 hr at 39 °C – mated after 48 hr / 4 hr at 39 °C – mated after 72 hr | | 0.950 | 0.081 | -0.600 | 1.000 |  |
|  | 1 hr at 41 °C – mated after 72 hr / 1 hr at 42 °C – mated after 72 hr | | 1.143 | 0.096 | 1.589 | 0.971 |  |
|  | 1 hr at 41 °C – mated after 72 hr / 4 hr at 38 °C – mated after 72 hr | | 1.178 | 0.066 | 2.924 | 0.203 |  |
|  | 1 hr at 41 °C – mated after 72 hr / 4 hr at 39 °C – mated after 72 hr | | 1.072 | 0.073 | 1.031 | 1.000 |  |
|  | 1 hr at 42 °C – mated after 72 hr / 4 hr at 38 °C – mated after 72 hr | | 1.031 | 0.092 | 0.337 | 1.000 |  |
|  | 1 hr at 42 °C – mated after 72 hr / 4 hr at 39 °C – mated after 72 hr | | 0.938 | 0.091 | -0.656 | 1.000 |  |
|  | 4 hr at 38 °C – mated after 72 hr / 4 hr at 39 °C – mated after 72 hr | | 0.910 | 0.068 | -1.264 | 0.997 |  |
| Egg hatchability | |  | |  |  |  |  |
|  | 1 hr at 41 °C – mated after 1 hr / 1 hr at 42 °C – mated after 1 hr | 3.167 | | 1.390 | 2.628 | 0.379 |  |
|  | 1 hr at 41 °C – mated after 1 hr / 4 hr at 38 °C – mated after 1 hr | 0.912 | | 0.382 | -0.220 | 1.000 |  |
|  | 1 hr at 41 °C – mated after 1 hr / 4 hr at 39 °C – mated after 1 hr | 0.656 | | 0.284 | -0.974 | 1.000 |  |
|  | 1 hr at 41 °C – mated after 1 hr / 1 hr at 41 °C – mated after 24 hr | 1.494 | | 0.601 | 0.997 | 1.000 |  |
|  | 1 hr at 41 °C – mated after 1 hr / 1 hr at 42 °C – mated after 24 hr | 0.341 | | 0.155 | -2.370 | 0.571 |  |
|  | 1 hr at 41 °C – mated after 1 hr / 4 hr at 38 °C – mated after 24 hr | 0.779 | | 0.330 | -0.589 | 1.000 |  |
|  | 1 hr at 41 °C – mated after 1 hr / 4 hr at 39 °C – mated after 24 hr | 0.353 | | 0.130 | -2.821 | 0.257 |  |
|  | 1 hr at 41 °C – mated after 1 hr / 1 hr at 41 °C – mated after 48 hr | 0.446 | | 0.195 | -1.848 | 0.899 |  |
|  | 1 hr at 41 °C – mated after 1 hr / 1 hr at 42 °C – mated after 48 hr | 0.222 | | 0.114 | -2.939 | 0.196 |  |
|  | 1 hr at 41 °C – mated after 1 hr / 4 hr at 38 °C – mated after 48 hr | 0.328 | | 0.130 | -2.809 | 0.263 |  |
|  | 1 hr at 41 °C – mated after 1 hr / 4 hr at 39 °C – mated after 48 hr | 0.283 | | 0.105 | -3.395 | 0.055 | **^⋅^** |
|  | 1 hr at 41 °C – mated after 1 hr / 1 hr at 41 °C – mated after 72 hr | 0.820 | | 0.340 | -0.480 | 1.000 |  |
|  | 1 hr at 41 °C – mated after 1 hr / 1 hr at 42 °C – mated after 72 hr | 0.744 | | 0.367 | -0.600 | 1.000 |  |
|  | 1 hr at 41 °C – mated after 1 hr / 4 hr at 38 °C – mated after 72 hr | 0.517 | | 0.194 | -1.757 | 0.931 |  |
|  | 1 hr at 41 °C – mated after 1 hr / 4 hr at 39 °C – mated after 72 hr | 0.496 | | 0.195 | -1.783 | 0.923 |  |
|  | 1 hr at 42 °C – mated after 1 hr / 4 hr at 38 °C – mated after 1 hr | 0.288 | | 0.136 | -2.629 | 0.378 |  |
|  | 1 hr at 42 °C – mated after 1 hr / 4 hr at 39 °C – mated after 1 hr | 0.207 | | 0.101 | -3.239 | 0.088 | **^⋅^** |
|  | 1 hr at 42 °C – mated after 1 hr / 1 hr at 41 °C – mated after 24 hr | 0.472 | | 0.216 | -1.637 | 0.962 |  |
|  | 1 hr at 42 °C – mated after 1 hr / 1 hr at 42 °C – mated after 24 hr | 0.108 | | 0.054 | -4.416 | 0.001 | ^**^ |
|  | 1 hr at 42 °C – mated after 1 hr / 4 hr at 38 °C – mated after 24 hr | 0.246 | | 0.117 | -2.937 | 0.197 |  |
|  | 1 hr at 42 °C – mated after 1 hr / 4 hr at 39 °C – mated after 24 hr | 0.111 | | 0.048 | -5.102 | < 0.001 | ^***^ |
|  | 1 hr at 42 °C – mated after 1 hr / 1 hr at 41 °C – mated after 48 hr | 0.141 | | 0.069 | -4.007 | 0.006 | ^**^ |
|  | 1 hr at 42 °C – mated after 1 hr / 1 hr at 42 °C – mated after 48 hr | 0.070 | | 0.039 | -4.766 | < 0.001 | ^***^ |
|  | 1 hr at 42 °C – mated after 1 hr / 4 hr at 38 °C – mated after 48 hr | 0.103 | | 0.047 | -4.994 | < 0.001 | ^***^ |
|  | 1 hr at 42 °C – mated after 1 hr / 4 hr at 39 °C – mated after 48 hr | 0.089 | | 0.039 | -5.586 | < 0.001 | ^***^ |
|  | 1 hr at 42 °C – mated after 1 hr / 1 hr at 41 °C – mated after 72 hr | 0.259 | | 0.121 | -2.881 | 0.225 |  |
|  | 1 hr at 42 °C – mated after 1 hr / 1 hr at 42 °C – mated after 72 hr | 0.235 | | 0.127 | -2.684 | 0.340 |  |
|  | 1 hr at 42 °C – mated after 1 hr / 4 hr at 38 °C – mated after 72 hr | 0.163 | | 0.071 | -4.161 | 0.003 | ^**^ |
|  | 1 hr at 42 °C – mated after 1 hr / 4 hr at 39 °C – mated after 72 hr | 0.157 | | 0.071 | -4.113 | 0.004 | ^**^ |
|  | 4 hr at 38 °C – mated after 1 hr / 4 hr at 39 °C – mated after 1 hr | 0.719 | | 0.337 | -0.704 | 1.000 |  |
|  | 4 hr at 38 °C – mated after 1 hr / 1 hr at 41 °C – mated after 24 hr | 1.638 | | 0.721 | 1.121 | 0.999 |  |
|  | 4 hr at 38 °C – mated after 1 hr / 1 hr at 42 °C – mated after 24 hr | 0.374 | | 0.182 | -2.016 | 0.817 |  |
|  | 4 hr at 38 °C – mated after 1 hr / 4 hr at 38 °C – mated after 24 hr | 0.855 | | 0.393 | -0.342 | 1.000 |  |
|  | 4 hr at 38 °C – mated after 1 hr / 4 hr at 39 °C – mated after 24 hr | 0.387 | | 0.159 | -2.315 | 0.612 |  |
|  | 4 hr at 38 °C – mated after 1 hr / 1 hr at 41 °C – mated after 48 hr | 0.489 | | 0.231 | -1.516 | 0.981 |  |
|  | 4 hr at 38 °C – mated after 1 hr / 1 hr at 42 °C – mated after 48 hr | 0.243 | | 0.132 | -2.606 | 0.394 |  |
|  | 4 hr at 38 °C – mated after 1 hr / 4 hr at 38 °C – mated after 48 hr | 0.359 | | 0.156 | -2.351 | 0.585 |  |
|  | 4 hr at 38 °C – mated after 1 hr / 4 hr at 39 °C – mated after 48 hr | 0.310 | | 0.128 | -2.838 | 0.247 |  |
|  | 4 hr at 38 °C – mated after 1 hr / 1 hr at 41 °C – mated after 72 hr | 0.899 | | 0.406 | -0.237 | 1.000 |  |
|  | 4 hr at 38 °C – mated after 1 hr / 1 hr at 42 °C – mated after 72 hr | 0.816 | | 0.428 | -0.388 | 1.000 |  |
|  | 4 hr at 38 °C – mated after 1 hr / 4 hr at 38 °C – mated after 72 hr | 0.567 | | 0.236 | -1.365 | 0.993 |  |
|  | 4 hr at 38 °C – mated after 1 hr / 4 hr at 39 °C – mated after 72 hr | 0.544 | | 0.235 | -1.410 | 0.991 |  |
|  | 4 hr at 39 °C – mated after 1 hr / 1 hr at 41 °C – mated after 24 hr | 2.279 | | 1.030 | 1.814 | 0.912 |  |
|  | 4 hr at 39 °C – mated after 1 hr / 1 hr at 42 °C – mated after 24 hr | 0.520 | | 0.260 | -1.307 | 0.996 |  |
|  | 4 hr at 39 °C – mated after 1 hr / 4 hr at 38 °C – mated after 24 hr | 1.189 | | 0.562 | 0.365 | 1.000 |  |
|  | 4 hr at 39 °C – mated after 1 hr / 4 hr at 39 °C – mated after 24 hr | 0.538 | | 0.229 | -1.459 | 0.987 |  |
|  | 4 hr at 39 °C – mated after 1 hr / 1 hr at 41 °C – mated after 48 hr | 0.681 | | 0.330 | -0.795 | 1.000 |  |
|  | 4 hr at 39 °C – mated after 1 hr / 1 hr at 42 °C – mated after 48 hr | 0.338 | | 0.187 | -1.958 | 0.849 |  |
|  | 4 hr at 39 °C – mated after 1 hr / 4 hr at 38 °C – mated after 48 hr | 0.500 | | 0.224 | -1.545 | 0.977 |  |
|  | 4 hr at 39 °C – mated after 1 hr / 4 hr at 39 °C – mated after 48 hr | 0.431 | | 0.184 | -1.970 | 0.842 |  |
|  | 4 hr at 39 °C – mated after 1 hr / 1 hr at 41 °C – mated after 72 hr | 1.250 | | 0.581 | 0.481 | 1.000 |  |
|  | 4 hr at 39 °C – mated after 1 hr / 1 hr at 42 °C – mated after 72 hr | 1.135 | | 0.608 | 0.235 | 1.000 |  |
|  | 4 hr at 39 °C – mated after 1 hr / 4 hr at 38 °C – mated after 72 hr | 0.788 | | 0.339 | -0.553 | 1.000 |  |
|  | 4 hr at 39 °C – mated after 1 hr / 4 hr at 39 °C – mated after 72 hr | 0.757 | | 0.337 | -0.626 | 1.000 |  |
|  | 1 hr at 41 °C – mated after 24 hr / 1 hr at 42 °C – mated after 24 hr | | 0.228 | 0.108 | -3.119 | 0.124 |  |
|  | 1 hr at 41 °C – mated after 24 hr / 4 hr at 38 °C – mated after 24 hr | | 0.522 | 0.232 | -1.464 | 0.986 |  |
|  | 1 hr at 41 °C – mated after 24 hr / 4 hr at 39 °C – mated after 24 hr | | 0.236 | 0.093 | -3.671 | 0.022 | ^*^ |
|  | 1 hr at 41 °C – mated after 24 hr / 1 hr at 41 °C – mated after 48 hr | | 0.299 | 0.136 | -2.645 | 0.367 |  |
|  | 1 hr at 41 °C – mated after 24 hr / 1 hr at 42 °C – mated after 48 hr | | 0.148 | 0.079 | -3.600 | 0.028 | ^*^ |
|  | 1 hr at 41 °C – mated after 24 hr / 4 hr at 38 °C – mated after 48 hr | | 0.219 | 0.092 | -3.617 | 0.027 | ^*^ |
|  | 1 hr at 41 °C – mated after 24 hr / 4 hr at 39 °C – mated after 48 hr | | 0.189 | 0.075 | -4.206 | 0.003 | ^**^ |
|  | 1 hr at 41 °C – mated after 24 hr / 1 hr at 41 °C – mated after 72 hr | | 0.549 | 0.239 | -1.378 | 0.993 |  |
|  | 1 hr at 41 °C – mated after 24 hr / 1 hr at 42 °C – mated after 72 hr | | 0.498 | 0.255 | -1.363 | 0.993 |  |
|  | 1 hr at 41 °C – mated after 24 hr / 4 hr at 38 °C – mated after 72 hr | | 0.346 | 0.138 | -2.659 | 0.357 |  |
|  | 1 hr at 41 °C – mated after 24 hr / 4 hr at 39 °C – mated after 72 hr | | 0.332 | 0.138 | -2.652 | 0.362 |  |
|  | 1 hr at 42 °C – mated after 24 hr / 4 hr at 38 °C – mated after 24 hr | | 2.285 | 1.120 | 1.681 | 0.952 |  |
|  | 1 hr at 42 °C – mated after 24 hr / 4 hr at 39 °C – mated after 24 hr | | 1.034 | 0.461 | 0.075 | 1.000 |  |
|  | 1 hr at 42 °C – mated after 24 hr / 1 hr at 41 °C – mated after 48 hr | | 1.308 | 0.658 | 0.534 | 1.000 |  |
|  | 1 hr at 42 °C – mated after 24 hr / 1 hr at 42 °C – mated after 48 hr | | 0.650 | 0.371 | -0.756 | 1.000 |  |
|  | 1 hr at 42 °C – mated after 24 hr / 4 hr at 38 °C – mated after 48 hr | | 0.960 | 0.450 | -0.087 | 1.000 |  |
|  | 1 hr at 42 °C – mated after 24 hr / 4 hr at 39 °C – mated after 48 hr | | 0.828 | 0.371 | -0.421 | 1.000 |  |
|  | 1 hr at 42 °C – mated after 24 hr / 1 hr at 41 °C – mated after 72 hr | | 2.403 | 1.160 | 1.813 | 0.912 |  |
|  | 1 hr at 42 °C – mated after 24 hr / 1 hr at 42 °C – mated after 72 hr | | 2.181 | 1.210 | 1.410 | 0.991 |  |
|  | 1 hr at 42 °C – mated after 24 hr / 4 hr at 38 °C – mated after 72 hr | | 1.515 | 0.683 | 0.921 | 1.000 |  |
|  | 1 hr at 42 °C – mated after 24 hr / 4 hr at 39 °C – mated after 72 hr | | 1.455 | 0.677 | 0.805 | 1.000 |  |
|  | 4 hr at 38 °C – mated after 24 hr / 4 hr at 39 °C – mated after 24 hr | | 0.453 | 0.188 | -1.911 | 0.872 |  |
|  | 4 hr at 38 °C – mated after 24 hr / 1 hr at 41 °C – mated after 48 hr | | 0.573 | 0.272 | -1.172 | 0.999 |  |
|  | 4 hr at 38 °C – mated after 24 hr / 1 hr at 42 °C – mated after 48 hr | | 0.285 | 0.155 | -2.301 | 0.623 |  |
|  | 4 hr at 38 °C – mated after 24 hr / 4 hr at 38 °C – mated after 48 hr | | 0.420 | 0.185 | -1.971 | 0.842 |  |
|  | 4 hr at 38 °C – mated after 24 hr / 4 hr at 39 °C – mated after 48 hr | | 0.363 | 0.151 | -2.431 | 0.524 |  |
|  | 4 hr at 38 °C – mated after 24 hr / 1 hr at 41 °C – mated after 72 hr | | 1.052 | 0.479 | 0.111 | 1.000 |  |
|  | 4 hr at 38 °C – mated after 24 hr / 1 hr at 42 °C – mated after 72 hr | | 0.955 | 0.504 | -0.088 | 1.000 |  |
|  | 4 hr at 38 °C – mated after 24 hr / 4 hr at 38 °C – mated after 72 hr | | 0.663 | 0.279 | -0.977 | 1.000 |  |
|  | 4 hr at 38 °C – mated after 24 hr / 4 hr at 39 °C – mated after 72 hr | | 0.637 | 0.278 | -1.035 | 1.000 |  |
|  | 4 hr at 39 °C – mated after 24 hr / 1 hr at 41 °C – mated after 48 hr | | 1.265 | 0.541 | 0.549 | 1.000 |  |
|  | 4 hr at 39 °C – mated after 24 hr / 1 hr at 42 °C – mated after 48 hr | | 0.629 | 0.318 | -0.919 | 1.000 |  |
|  | 4 hr at 39 °C – mated after 24 hr / 4 hr at 38 °C – mated after 48 hr | | 0.929 | 0.360 | -0.191 | 1.000 |  |
|  | 4 hr at 39 °C – mated after 24 hr / 4 hr at 39 °C – mated after 48 hr | | 0.801 | 0.290 | -0.613 | 1.000 |  |
|  | 4 hr at 39 °C – mated after 24 hr / 1 hr at 41 °C – mated after 72 hr | | 2.324 | 0.942 | 2.080 | 0.778 |  |
|  | 4 hr at 39 °C – mated after 24 hr / 1 hr at 42 °C – mated after 72 hr | | 2.109 | 1.020 | 1.536 | 0.979 |  |
|  | 4 hr at 39 °C – mated after 24 hr / 4 hr at 38 °C – mated after 72 hr | | 1.465 | 0.536 | 1.043 | 1.000 |  |
|  | 4 hr at 39 °C – mated after 24 hr / 4 hr at 39 °C – mated after 72 hr | | 1.407 | 0.540 | 0.889 | 1.000 |  |
|  | 1 hr at 41 °C – mated after 48 hr / 1 hr at 42 °C – mated after 48 hr | | 0.497 | 0.276 | -1.257 | 0.997 |  |
|  | 1 hr at 41 °C – mated after 48 hr / 4 hr at 38 °C – mated after 48 hr | | 0.734 | 0.332 | -0.684 | 1.000 |  |
|  | 1 hr at 41 °C – mated after 48 hr / 4 hr at 39 °C – mated after 48 hr | | 0.633 | 0.273 | -1.062 | 1.000 |  |
|  | 1 hr at 41 °C – mated after 48 hr / 1 hr at 41 °C – mated after 72 hr | | 1.837 | 0.859 | 1.301 | 0.996 |  |
|  | 1 hr at 41 °C – mated after 48 hr / 1 hr at 42 °C – mated after 72 hr | | 1.667 | 0.898 | 0.949 | 1.000 |  |
|  | 1 hr at 41 °C – mated after 48 hr / 4 hr at 38 °C – mated after 72 hr | | 1.158 | 0.502 | 0.338 | 1.000 |  |
|  | 1 hr at 41 °C – mated after 48 hr / 4 hr at 39 °C – mated after 72 hr | | 1.112 | 0.499 | 0.237 | 1.000 |  |
|  | 1 hr at 42 °C – mated after 48 hr / 4 hr at 38 °C – mated after 48 hr | | 1.477 | 0.777 | 0.741 | 1.000 |  |
|  | 1 hr at 42 °C – mated after 48 hr / 4 hr at 39 °C – mated after 48 hr | | 1.274 | 0.646 | 0.477 | 1.000 |  |
|  | 1 hr at 42 °C – mated after 48 hr / 1 hr at 41 °C – mated after 72 hr | | 3.696 | 1.990 | 2.426 | 0.528 |  |
|  | 1 hr at 42 °C – mated after 48 hr / 1 hr at 42 °C – mated after 72 hr | | 3.355 | 2.020 | 2.011 | 0.820 |  |
|  | 1 hr at 42 °C – mated after 48 hr / 4 hr at 38 °C – mated after 72 hr | | 2.330 | 1.190 | 1.659 | 0.957 |  |
|  | 1 hr at 42 °C – mated after 48 hr / 4 hr at 39 °C – mated after 72 hr | | 2.238 | 1.170 | 1.540 | 0.978 |  |
|  | 4 hr at 38 °C – mated after 48 hr / 4 hr at 39 °C – mated after 48 hr | | 0.863 | 0.337 | -0.379 | 1.000 |  |
|  | 4 hr at 38 °C – mated after 48 hr / 1 hr at 41 °C – mated after 72 hr | | 2.503 | 1.080 | 2.129 | 0.747 |  |
|  | 4 hr at 38 °C – mated after 48 hr / 1 hr at 42 °C – mated after 72 hr | | 2.271 | 1.150 | 1.617 | 0.966 |  |
|  | 4 hr at 38 °C – mated after 48 hr / 4 hr at 38 °C – mated after 72 hr | | 1.578 | 0.622 | 1.157 | 1.000 |  |
|  | 4 hr at 38 °C – mated after 48 hr / 4 hr at 39 °C – mated after 72 hr | | 1.515 | 0.622 | 1.012 | 1.000 |  |
|  | 4 hr at 39 °C – mated after 48 hr / 1 hr at 41 °C – mated after 72 hr | | 2.902 | 1.180 | 2.611 | 0.391 |  |
|  | 4 hr at 39 °C – mated after 48 hr / 1 hr at 42 °C – mated after 72 hr | | 2.634 | 1.290 | 1.984 | 0.835 |  |
|  | 4 hr at 39 °C – mated after 48 hr / 4 hr at 38 °C – mated after 72 hr | | 1.829 | 0.675 | 1.637 | 0.962 |  |
|  | 4 hr at 39 °C – mated after 48 hr / 4 hr at 39 °C – mated after 72 hr | | 1.756 | 0.679 | 1.458 | 0.987 |  |
|  | 1 hr at 41 °C – mated after 72 hr / 1 hr at 42 °C – mated after 72 hr | | 0.908 | 0.473 | -0.186 | 1.000 |  |
|  | 1 hr at 41 °C – mated after 72 hr / 4 hr at 38 °C – mated after 72 hr | | 0.630 | 0.259 | -1.122 | 0.999 |  |
|  | 1 hr at 41 °C – mated after 72 hr / 4 hr at 39 °C – mated after 72 hr | | 0.605 | 0.259 | -1.176 | 0.999 |  |
|  | 1 hr at 42 °C – mated after 72 hr / 4 hr at 38 °C – mated after 72 hr | | 0.695 | 0.341 | -0.743 | 1.000 |  |
|  | 1 hr at 42 °C – mated after 72 hr / 4 hr at 39 °C – mated after 72 hr | | 0.667 | 0.336 | -0.803 | 1.000 |  |
|  | 4 hr at 38 °C – mated after 72 hr / 4 hr at 39 °C – mated after 72 hr | | 0.960 | 0.374 | -0.104 | 1.000 |  |
| Viable offspring | |  | |  |  |  |  |
|  | 1 hr at 41 °C – mated after 1 hr / 1 hr at 42 °C – mated after 1 hr | 1.172 | | 0.205 | 0.904 | 1.000 |  |
|  | 1 hr at 41 °C – mated after 1 hr / 4 hr at 38 °C – mated after 1 hr | 0.979 | | 0.110 | -0.191 | 1.000 |  |
|  | 1 hr at 41 °C – mated after 1 hr / 4 hr at 39 °C – mated after 1 hr | 1.014 | | 0.096 | 0.151 | 1.000 |  |
|  | 1 hr at 41 °C – mated after 1 hr / 1 hr at 41 °C – mated after 24 hr | 0.936 | | 0.120 | -0.515 | 1.000 |  |
|  | 1 hr at 41 °C – mated after 1 hr / 1 hr at 42 °C – mated after 24 hr | 0.862 | | 0.075 | -1.715 | 0.943 |  |
|  | 1 hr at 41 °C – mated after 1 hr / 4 hr at 38 °C – mated after 24 hr | 0.889 | | 0.088 | -1.192 | 0.999 |  |
|  | 1 hr at 41 °C – mated after 1 hr / 4 hr at 39 °C – mated after 24 hr | 0.945 | | 0.094 | -0.565 | 1.000 |  |
|  | 1 hr at 41 °C – mated after 1 hr / 1 hr at 41 °C – mated after 48 hr | 0.867 | | 0.072 | -1.725 | 0.941 |  |
|  | 1 hr at 41 °C – mated after 1 hr / 1 hr at 42 °C – mated after 48 hr | 0.777 | | 0.067 | -2.940 | 0.196 |  |
|  | 1 hr at 41 °C – mated after 1 hr / 4 hr at 38 °C – mated after 48 hr | 0.810 | | 0.071 | -2.397 | 0.550 |  |
|  | 1 hr at 41 °C – mated after 1 hr / 4 hr at 39 °C – mated after 48 hr | 0.892 | | 0.090 | -1.131 | 0.999 |  |
|  | 1 hr at 41 °C – mated after 1 hr / 1 hr at 41 °C – mated after 72 hr | 0.795 | | 0.074 | -2.471 | 0.494 |  |
|  | 1 hr at 41 °C – mated after 1 hr / 1 hr at 42 °C – mated after 72 hr | 0.843 | | 0.082 | -1.750 | 0.933 |  |
|  | 1 hr at 41 °C – mated after 1 hr / 4 hr at 38 °C – mated after 72 hr | 0.964 | | 0.083 | -0.430 | 1.000 |  |
|  | 1 hr at 41 °C – mated after 1 hr / 4 hr at 39 °C – mated after 72 hr | 0.842 | | 0.077 | -1.883 | 0.884 |  |
|  | 1 hr at 42 °C – mated after 1 hr / 4 hr at 38 °C – mated after 1 hr | 0.835 | | 0.154 | -0.978 | 1.000 |  |
|  | 1 hr at 42 °C – mated after 1 hr / 4 hr at 39 °C – mated after 1 hr | 0.866 | | 0.150 | -0.830 | 1.000 |  |
|  | 1 hr at 42 °C – mated after 1 hr / 1 hr at 41 °C – mated after 24 hr | 0.799 | | 0.155 | -1.157 | 0.999 |  |
|  | 1 hr at 42 °C – mated after 1 hr / 1 hr at 42 °C – mated after 24 hr | 0.736 | | 0.125 | -1.814 | 0.912 |  |
|  | 1 hr at 42 °C – mated after 1 hr / 4 hr at 38 °C – mated after 24 hr | 0.759 | | 0.133 | -1.571 | 0.973 |  |
|  | 1 hr at 42 °C – mated after 1 hr / 4 hr at 39 °C – mated after 24 hr | 0.807 | | 0.142 | -1.218 | 0.998 |  |
|  | 1 hr at 42 °C – mated after 1 hr / 1 hr at 41 °C – mated after 48 hr | 0.740 | | 0.124 | -1.801 | 0.917 |  |
|  | 1 hr at 42 °C – mated after 1 hr / 1 hr at 42 °C – mated after 48 hr | 0.664 | | 0.112 | -2.431 | 0.524 |  |
|  | 1 hr at 42 °C – mated after 1 hr / 4 hr at 38 °C – mated after 48 hr | 0.692 | | 0.117 | -2.171 | 0.718 |  |
|  | 1 hr at 42 °C – mated after 1 hr / 4 hr at 39 °C – mated after 48 hr | 0.761 | | 0.135 | -1.540 | 0.978 |  |
|  | 1 hr at 42 °C – mated after 1 hr / 1 hr at 41 °C – mated after 72 hr | 0.679 | | 0.117 | -2.246 | 0.664 |  |
|  | 1 hr at 42 °C – mated after 1 hr / 1 hr at 42 °C – mated after 72 hr | 0.719 | | 0.126 | -1.880 | 0.885 |  |
|  | 1 hr at 42 °C – mated after 1 hr / 4 hr at 38 °C – mated after 72 hr | 0.822 | | 0.139 | -1.156 | 0.999 |  |
|  | 1 hr at 42 °C – mated after 1 hr / 4 hr at 39 °C – mated after 72 hr | 0.718 | | 0.123 | -1.926 | 0.865 |  |
|  | 4 hr at 38 °C – mated after 1 hr / 4 hr at 39 °C – mated after 1 hr | 1.037 | | 0.114 | 0.326 | 1.000 |  |
|  | 4 hr at 38 °C – mated after 1 hr / 1 hr at 41 °C – mated after 24 hr | 0.956 | | 0.134 | -0.319 | 1.000 |  |
|  | 4 hr at 38 °C – mated after 1 hr / 1 hr at 42 °C – mated after 24 hr | 0.881 | | 0.091 | -1.234 | 0.998 |  |
|  | 4 hr at 38 °C – mated after 1 hr / 4 hr at 38 °C – mated after 24 hr | 0.908 | | 0.103 | -0.849 | 1.000 |  |
|  | 4 hr at 38 °C – mated after 1 hr / 4 hr at 39 °C – mated after 24 hr | 0.966 | | 0.110 | -0.304 | 1.000 |  |
|  | 4 hr at 38 °C – mated after 1 hr / 1 hr at 41 °C – mated after 48 hr | 0.886 | | 0.088 | -1.216 | 0.998 |  |
|  | 4 hr at 38 °C – mated after 1 hr / 1 hr at 42 °C – mated after 48 hr | 0.794 | | 0.081 | -2.254 | 0.658 |  |
|  | 4 hr at 38 °C – mated after 1 hr / 4 hr at 38 °C – mated after 48 hr | 0.828 | | 0.086 | -1.817 | 0.911 |  |
|  | 4 hr at 38 °C – mated after 1 hr / 4 hr at 39 °C – mated after 48 hr | 0.911 | | 0.105 | -0.804 | 1.000 |  |
|  | 4 hr at 38 °C – mated after 1 hr / 1 hr at 41 °C – mated after 72 hr | 0.813 | | 0.088 | -1.919 | 0.868 |  |
|  | 4 hr at 38 °C – mated after 1 hr / 1 hr at 42 °C – mated after 72 hr | 0.861 | | 0.097 | -1.329 | 0.995 |  |
|  | 4 hr at 38 °C – mated after 1 hr / 4 hr at 38 °C – mated after 72 hr | 0.985 | | 0.101 | -0.152 | 1.000 |  |
|  | 4 hr at 38 °C – mated after 1 hr / 4 hr at 39 °C – mated after 72 hr | 0.860 | | 0.092 | -1.409 | 0.991 |  |
|  | 4 hr at 39 °C – mated after 1 hr / 1 hr at 41 °C – mated after 24 hr | 0.923 | | 0.117 | -0.638 | 1.000 |  |
|  | 4 hr at 39 °C – mated after 1 hr / 1 hr at 42 °C – mated after 24 hr | 0.850 | | 0.071 | -1.956 | 0.849 |  |
|  | 4 hr at 39 °C – mated after 1 hr / 4 hr at 38 °C – mated after 24 hr | 0.876 | | 0.084 | -1.377 | 0.993 |  |
|  | 4 hr at 39 °C – mated after 1 hr / 4 hr at 39 °C – mated after 24 hr | 0.932 | | 0.090 | -0.729 | 1.000 |  |
|  | 4 hr at 39 °C – mated after 1 hr / 1 hr at 41 °C – mated after 48 hr | 0.854 | | 0.068 | -1.982 | 0.836 |  |
|  | 4 hr at 39 °C – mated after 1 hr / 1 hr at 42 °C – mated after 48 hr | 0.766 | | 0.063 | -3.236 | 0.089 | **^⋅^** |
|  | 4 hr at 39 °C – mated after 1 hr / 4 hr at 38 °C – mated after 48 hr | 0.799 | | 0.067 | -2.661 | 0.356 |  |
|  | 4 hr at 39 °C – mated after 1 hr / 4 hr at 39 °C – mated after 48 hr | 0.879 | | 0.086 | -1.310 | 0.996 |  |
|  | 4 hr at 39 °C – mated after 1 hr / 1 hr at 41 °C – mated after 72 hr | 0.784 | | 0.070 | -2.717 | 0.319 |  |
|  | 4 hr at 39 °C – mated after 1 hr / 1 hr at 42 °C – mated after 72 hr | 0.831 | | 0.079 | -1.956 | 0.849 |  |
|  | 4 hr at 39 °C – mated after 1 hr / 4 hr at 38 °C – mated after 72 hr | 0.950 | | 0.079 | -0.620 | 1.000 |  |
|  | 4 hr at 39 °C – mated after 1 hr / 4 hr at 39 °C – mated after 72 hr | 0.830 | | 0.073 | -2.112 | 0.758 |  |
|  | 1 hr at 41 °C – mated after 24 hr / 1 hr at 42 °C – mated after 24 hr | | 0.921 | 0.111 | -0.685 | 1.000 |  |
|  | 1 hr at 41 °C – mated after 24 hr / 4 hr at 38 °C – mated after 24 hr | | 0.950 | 0.123 | -0.399 | 1.000 |  |
|  | 1 hr at 41 °C – mated after 24 hr / 4 hr at 39 °C – mated after 24 hr | | 1.010 | 0.131 | 0.078 | 1.000 |  |
|  | 1 hr at 41 °C – mated after 24 hr / 1 hr at 41 °C – mated after 48 hr | | 0.926 | 0.109 | -0.652 | 1.000 |  |
|  | 1 hr at 41 °C – mated after 24 hr / 1 hr at 42 °C – mated after 48 hr | | 0.831 | 0.099 | -1.551 | 0.976 |  |
|  | 1 hr at 41 °C – mated after 24 hr / 4 hr at 38 °C – mated after 48 hr | | 0.866 | 0.105 | -1.189 | 0.999 |  |
|  | 1 hr at 41 °C – mated after 24 hr / 4 hr at 39 °C – mated after 48 hr | | 0.953 | 0.125 | -0.367 | 1.000 |  |
|  | 1 hr at 41 °C – mated after 24 hr / 1 hr at 41 °C – mated after 72 hr | | 0.850 | 0.106 | -1.304 | 0.996 |  |
|  | 1 hr at 41 °C – mated after 24 hr / 1 hr at 42 °C – mated after 72 hr | | 0.901 | 0.116 | -0.814 | 1.000 |  |
|  | 1 hr at 41 °C – mated after 24 hr / 4 hr at 38 °C – mated after 72 hr | | 1.030 | 0.124 | 0.243 | 1.000 |  |
|  | 1 hr at 41 °C – mated after 24 hr / 4 hr at 39 °C – mated after 72 hr | | 0.899 | 0.111 | -0.858 | 1.000 |  |
|  | 1 hr at 42 °C – mated after 24 hr / 4 hr at 38 °C – mated after 24 hr | | 1.031 | 0.091 | 0.350 | 1.000 |  |
|  | 1 hr at 42 °C – mated after 24 hr / 4 hr at 39 °C – mated after 24 hr | | 1.097 | 0.097 | 1.044 | 1.000 |  |
|  | 1 hr at 42 °C – mated after 24 hr / 1 hr at 41 °C – mated after 48 hr | | 1.006 | 0.070 | 0.083 | 1.000 |  |
|  | 1 hr at 42 °C – mated after 24 hr / 1 hr at 42 °C – mated after 48 hr | | 0.902 | 0.065 | -1.421 | 0.990 |  |
|  | 1 hr at 42 °C – mated after 24 hr / 4 hr at 38 °C – mated after 48 hr | | 0.940 | 0.071 | -0.821 | 1.000 |  |
|  | 1 hr at 42 °C – mated after 24 hr / 4 hr at 39 °C – mated after 48 hr | | 1.035 | 0.093 | 0.380 | 1.000 |  |
|  | 1 hr at 42 °C – mated after 24 hr / 1 hr at 41 °C – mated after 72 hr | | 0.923 | 0.075 | -0.994 | 1.000 |  |
|  | 1 hr at 42 °C – mated after 24 hr / 1 hr at 42 °C – mated after 72 hr | | 0.978 | 0.085 | -0.257 | 1.000 |  |
|  | 1 hr at 42 °C – mated after 24 hr / 4 hr at 38 °C – mated after 72 hr | | 1.118 | 0.082 | 1.520 | 0.981 |  |
|  | 1 hr at 42 °C – mated after 24 hr / 4 hr at 39 °C – mated after 72 hr | | 0.976 | 0.078 | -0.300 | 1.000 |  |
|  | 4 hr at 38 °C – mated after 24 hr / 4 hr at 39 °C – mated after 24 hr | | 1.064 | 0.107 | 0.614 | 1.000 |  |
|  | 4 hr at 38 °C – mated after 24 hr / 1 hr at 41 °C – mated after 48 hr | | 0.975 | 0.082 | -0.297 | 1.000 |  |
|  | 4 hr at 38 °C – mated after 24 hr / 1 hr at 42 °C – mated after 48 hr | | 0.875 | 0.076 | -1.541 | 0.978 |  |
|  | 4 hr at 38 °C – mated after 24 hr / 4 hr at 38 °C – mated after 48 hr | | 0.912 | 0.081 | -1.038 | 1.000 |  |
|  | 4 hr at 38 °C – mated after 24 hr / 4 hr at 39 °C – mated after 48 hr | | 1.004 | 0.103 | 0.035 | 1.000 |  |
|  | 4 hr at 38 °C – mated after 24 hr / 1 hr at 41 °C – mated after 72 hr | | 0.895 | 0.084 | -1.183 | 0.999 |  |
|  | 4 hr at 38 °C – mated after 24 hr / 1 hr at 42 °C – mated after 72 hr | | 0.948 | 0.094 | -0.536 | 1.000 |  |
|  | 4 hr at 38 °C – mated after 24 hr / 4 hr at 38 °C – mated after 72 hr | | 1.084 | 0.095 | 0.923 | 1.000 |  |
|  | 4 hr at 38 °C – mated after 24 hr / 4 hr at 39 °C – mated after 72 hr | | 0.947 | 0.088 | -0.589 | 1.000 |  |
|  | 4 hr at 39 °C – mated after 24 hr / 1 hr at 41 °C – mated after 48 hr | | 0.917 | 0.078 | -1.022 | 1.000 |  |
|  | 4 hr at 39 °C – mated after 24 hr / 1 hr at 42 °C – mated after 48 hr | | 0.822 | 0.072 | -2.233 | 0.673 |  |
|  | 4 hr at 39 °C – mated after 24 hr / 4 hr at 38 °C – mated after 48 hr | | 0.857 | 0.077 | -1.720 | 0.942 |  |
|  | 4 hr at 39 °C – mated after 24 hr / 4 hr at 39 °C – mated after 48 hr | | 0.944 | 0.097 | -0.566 | 1.000 |  |
|  | 4 hr at 39 °C – mated after 24 hr / 1 hr at 41 °C – mated after 72 hr | | 0.841 | 0.079 | -1.829 | 0.906 |  |
|  | 4 hr at 39 °C – mated after 24 hr / 1 hr at 42 °C – mated after 72 hr | | 0.892 | 0.089 | -1.154 | 0.999 |  |
|  | 4 hr at 39 °C – mated after 24 hr / 4 hr at 38 °C – mated after 72 hr | | 1.019 | 0.090 | 0.216 | 1.000 |  |
|  | 4 hr at 39 °C – mated after 24 hr / 4 hr at 39 °C – mated after 72 hr | | 0.890 | 0.083 | -1.245 | 0.998 |  |
|  | 1 hr at 41 °C – mated after 48 hr / 1 hr at 42 °C – mated after 48 hr | | 0.897 | 0.061 | -1.602 | 0.969 |  |
|  | 1 hr at 41 °C – mated after 48 hr / 4 hr at 38 °C – mated after 48 hr | | 0.935 | 0.066 | -0.954 | 1.000 |  |
|  | 1 hr at 41 °C – mated after 48 hr / 4 hr at 39 °C – mated after 48 hr | | 1.029 | 0.089 | 0.330 | 1.000 |  |
|  | 1 hr at 41 °C – mated after 48 hr / 1 hr at 41 °C – mated after 72 hr | | 0.918 | 0.070 | -1.122 | 0.999 |  |
|  | 1 hr at 41 °C – mated after 48 hr / 1 hr at 42 °C – mated after 72 hr | | 0.972 | 0.080 | -0.338 | 1.000 |  |
|  | 1 hr at 41 °C – mated after 48 hr / 4 hr at 38 °C – mated after 72 hr | | 1.112 | 0.077 | 1.536 | 0.978 |  |
|  | 1 hr at 41 °C – mated after 48 hr / 4 hr at 39 °C – mated after 72 hr | | 0.971 | 0.073 | -0.393 | 1.000 |  |
|  | 1 hr at 42 °C – mated after 48 hr / 4 hr at 38 °C – mated after 48 hr | | 1.042 | 0.077 | 0.563 | 1.000 |  |
|  | 1 hr at 42 °C – mated after 48 hr / 4 hr at 39 °C – mated after 48 hr | | 1.147 | 0.102 | 1.539 | 0.978 |  |
|  | 1 hr at 42 °C – mated after 48 hr / 1 hr at 41 °C – mated after 72 hr | | 1.023 | 0.081 | 0.287 | 1.000 |  |
|  | 1 hr at 42 °C – mated after 48 hr / 1 hr at 42 °C – mated after 72 hr | | 1.084 | 0.093 | 0.948 | 1.000 |  |
|  | 1 hr at 42 °C – mated after 48 hr / 4 hr at 38 °C – mated after 72 hr | | 1.239 | 0.089 | 2.974 | 0.180 |  |
|  | 1 hr at 42 °C – mated after 48 hr / 4 hr at 39 °C – mated after 72 hr | | 1.082 | 0.085 | 1.010 | 1.000 |  |
|  | 4 hr at 38 °C – mated after 48 hr / 4 hr at 39 °C – mated after 48 hr | | 1.101 | 0.100 | 1.051 | 1.000 |  |
|  | 4 hr at 38 °C – mated after 48 hr / 1 hr at 41 °C – mated after 72 hr | | 0.981 | 0.080 | -0.228 | 1.000 |  |
|  | 4 hr at 38 °C – mated after 48 hr / 1 hr at 42 °C – mated after 72 hr | | 1.040 | 0.091 | 0.451 | 1.000 |  |
|  | 4 hr at 38 °C – mated after 48 hr / 4 hr at 38 °C – mated after 72 hr | | 1.189 | 0.089 | 2.320 | 0.608 |  |
|  | 4 hr at 38 °C – mated after 48 hr / 4 hr at 39 °C – mated after 72 hr | | 1.038 | 0.084 | 0.467 | 1.000 |  |
|  | 4 hr at 39 °C – mated after 48 hr / 1 hr at 41 °C – mated after 72 hr | | 0.892 | 0.086 | -1.193 | 0.998 |  |
|  | 4 hr at 39 °C – mated after 48 hr / 1 hr at 42 °C – mated after 72 hr | | 0.945 | 0.095 | -0.560 | 1.000 |  |
|  | 4 hr at 39 °C – mated after 48 hr / 4 hr at 38 °C – mated after 72 hr | | 1.080 | 0.097 | 0.858 | 1.000 |  |
|  | 4 hr at 39 °C – mated after 48 hr / 4 hr at 39 °C – mated after 72 hr | | 0.943 | 0.090 | -0.612 | 1.000 |  |
|  | 1 hr at 41 °C – mated after 72 hr / 1 hr at 42 °C – mated after 72 hr | | 1.060 | 0.098 | 0.629 | 1.000 |  |
|  | 1 hr at 41 °C – mated after 72 hr / 4 hr at 38 °C – mated after 72 hr | | 1.211 | 0.097 | 2.386 | 0.558 |  |
|  | 1 hr at 41 °C – mated after 72 hr / 4 hr at 39 °C – mated after 72 hr | | 1.058 | 0.091 | 0.655 | 1.000 |  |
|  | 1 hr at 42 °C – mated after 72 hr / 4 hr at 38 °C – mated after 72 hr | | 1.143 | 0.098 | 1.554 | 0.976 |  |
|  | 1 hr at 42 °C – mated after 72 hr / 4 hr at 39 °C – mated after 72 hr | | 0.998 | 0.091 | -0.019 | 1.000 |  |
|  | 4 hr at 38 °C – mated after 72 hr / 4 hr at 39 °C – mated after 72 hr | | 0.873 | 0.069 | -1.710 | 0.945 |  |

Significant *p*-values are indicated with asterisks (^***^ *p* < 0.001, ^**^ *p* < 0.01, ^*^ *p* < 0.05, **^⋅^** *p* < 0.1)

4. **Effect of mating status on heat shock induced changes in *Ae. aegypti* female fertility** (Table S7– S8)

Table S7. Analysis of deviance (Type II Wald Chi-square tests) of reproductive traits evaluating the effect of mating status on heat shock induced changes in *Ae. aegypti* female fertility

| Trait | Factor | df | χ^2^ | *p*-value |  |
| --- | --- | --- | --- | --- | --- |
| Egg laying success | |  |  |  |  |
|  | treatment (mating status) | 2 | 0.970 | 0.616 |  |
|  | gonotrophic cycle | 1 | 1.902 | 0.168 |  |
|  | interaction | 2 | 1.167 | 0.558 |  |
| Fecundity | |  |  |  |  |
|  | treatment (mating status) | 2 | 8.539 | 0.014 | ^*^ |
|  | gonotrophic cycle | 1 | 9.891 | 0.002 | ^**^ |
|  | interaction | 2 | 0.800 | 0.670 |  |
| Egg hatchability | |  |  |  |  |
|  | treatment (mating status) | 2 | 6.388 | 0.041 | ^*^ |
|  | gonotrophic cycle | 1 | 1.745 | 0.187 |  |
|  | interaction | 2 | 0.444 | 0.801 |  |
| Viable offspring | |  |  |  |  |
|  | treatment (mating status) | 2 | 8.744 | 0.013 | ^*^ |
|  | gonotrophic cycle | 1 | 2.878 | 0.090 | **^⋅^** |
|  | interaction | 2 | 0.129 | 0.938 |  |

Significant *p*-values are indicated with asterisks (^***^ *p* < 0.001, ^**^ *p* < 0.01, ^*^ *p* < 0.05, **^⋅^** *p* < 0.1)

Table S8. Full pairwise comparisons of reproductive traits evaluating the effect of mating status on heat shock induced changes in *Ae. aegypti* female fertility

| Trait | Comparison (treatment vs. gonotrophic cycle) | Ratio / OR | SE | Z-ratio | *p*-value |  |
| --- | --- | --- | --- | --- | --- | --- |
| Egg laying success | |  |  |  |  |  |
|  | control in gonotrophic cycle 1 /  pre-mating in gonotrophic cycle 1 | 0.321 | 0.533 | -0.684 | 0.984 |  |
|  | control in gonotrophic cycle 1 / post-mating in gonotrophic cycle 1 | 0.315 | 0.523 | -0.696 | 0.983 |  |
|  | control in gonotrophic cycle 1 / control in gonotrophic cycle 2 | 1.329 | 1.580 | 0.240 | 1.000 |  |
|  | control in gonotrophic cycle 1 / pre-mating in gonotrophic cycle 2 | 1.235 | 1.460 | 0.178 | 1.000 |  |
|  | control in gonotrophic cycle 1 / post-mating in gonotrophic cycle 2 | 3.356 | 3.390 | 1.199 | 0.838 |  |
|  | pre-mating in gonotrophic cycle 1 / post-mating in gonotrophic cycle 1 | 0.982 | 1.990 | -0.009 | 1.000 |  |
|  | pre-mating in gonotrophic cycle 1 / control in gonotrophic cycle 2 | 4.139 | 6.880 | 0.854 | 0.957 |  |
|  | pre-mating in gonotrophic cycle 1 / pre-mating in gonotrophic cycle 2 | 3.847 | 6.390 | 0.811 | 0.966 |  |
|  | pre-mating in gonotrophic cycle 1 / post-mating in gonotrophic cycle 2 | 10.452 | 16.100 | 1.522 | 0.650 |  |
|  | post-mating in gonotrophic cycle 1 / control in gonotrophic cycle 2 | 4.215 | 7.010 | 0.865 | 0.955 |  |
|  | post-mating in gonotrophic cycle 1 / pre-mating in gonotrophic cycle 2 | 3.918 | 6.510 | 0.822 | 0.964 |  |
|  | post-mating in gonotrophic cycle 1 / post-mating in gonotrophic cycle 2 | 10.644 | 16.400 | 1.534 | 0.642 |  |
|  | control in gonotrophic cycle 2 / pre-mating in gonotrophic cycle 2 | 0.929 | 1.110 | -0.062 | 1.000 |  |
|  | control in gonotrophic cycle 2 / post-mating in gonotrophic cycle 2 | 2.525 | 2.560 | 0.913 | 0.944 |  |
|  | pre-mating in gonotrophic cycle 2 / post-mating in gonotrophic cycle 2 | 2.717 | 2.750 | 0.986 | 0.923 |  |
| Fecundity | |  |  |  |  |  |
|  | control in gonotrophic cycle 1 /  pre-mating in gonotrophic cycle 1 | 1.051 | 0.035 | 1.503 | 0.663 |  |
|  | control in gonotrophic cycle 1 / post-mating in gonotrophic cycle 1 | 1.053 | 0.032 | 1.696 | 0.535 |  |
|  | control in gonotrophic cycle 1 / control in gonotrophic cycle 2 | 1.048 | 0.039 | 1.263 | 0.805 |  |
|  | control in gonotrophic cycle 1 / pre-mating in gonotrophic cycle 2 | 1.146 | 0.045 | 3.499 | 0.006 | ^**^ |
|  | control in gonotrophic cycle 1 / post-mating in gonotrophic cycle 2 | 1.154 | 0.050 | 3.305 | 0.012 | ^*^ |
|  | pre-mating in gonotrophic cycle 1 / post-mating in gonotrophic cycle 1 | 1.002 | 0.033 | 0.055 | 1.000 |  |
|  | pre-mating in gonotrophic cycle 1 / control in gonotrophic cycle 2 | 0.997 | 0.039 | -0.084 | 1.000 |  |
|  | pre-mating in gonotrophic cycle 1 / pre-mating in gonotrophic cycle 2 | 1.090 | 0.044 | 2.127 | 0.273 |  |
|  | pre-mating in gonotrophic cycle 1 / post-mating in gonotrophic cycle 2 | 1.098 | 0.049 | 2.083 | 0.296 |  |
|  | post-mating in gonotrophic cycle 1 / control in gonotrophic cycle 2 | 0.995 | 0.036 | -0.139 | 1.000 |  |
|  | post-mating in gonotrophic cycle 1 / pre-mating in gonotrophic cycle 2 | 1.088 | 0.042 | 2.203 | 0.236 |  |
|  | post-mating in gonotrophic cycle 1 / post-mating in gonotrophic cycle 2 | 1.096 | 0.047 | 2.138 | 0.268 |  |
|  | control in gonotrophic cycle 2 / pre-mating in gonotrophic cycle 2 | 1.094 | 0.048 | 2.051 | 0.314 |  |
|  | control in gonotrophic cycle 2 / post-mating in gonotrophic cycle 2 | 1.101 | 0.052 | 2.028 | 0.326 |  |
|  | pre-mating in gonotrophic cycle 2 / post-mating in gonotrophic cycle 2 | 1.007 | 0.050 | 0.146 | 1.000 |  |
| Egg hatchability | |  |  |  |  |  |
|  | control in gonotrophic cycle 1 /  pre-mating in gonotrophic cycle 1 | 1.709 | 0.508 | 1.803 | 0.464 |  |
|  | control in gonotrophic cycle 1 / post-mating in gonotrophic cycle 1 | 1.842 | 0.533 | 2.110 | 0.282 |  |
|  | control in gonotrophic cycle 1 / control in gonotrophic cycle 2 | 0.931 | 0.282 | -0.238 | 1.000 |  |
|  | control in gonotrophic cycle 1 / pre-mating in gonotrophic cycle 2 | 1.270 | 0.389 | 0.779 | 0.971 |  |
|  | control in gonotrophic cycle 1 / post-mating in gonotrophic cycle 2 | 1.316 | 0.370 | 0.978 | 0.925 |  |
|  | pre-mating in gonotrophic cycle 1 / post-mating in gonotrophic cycle 1 | 1.078 | 0.334 | 0.241 | 1.000 |  |
|  | pre-mating in gonotrophic cycle 1 / control in gonotrophic cycle 2 | 0.544 | 0.175 | -1.887 | 0.410 |  |
|  | pre-mating in gonotrophic cycle 1 / pre-mating in gonotrophic cycle 2 | 0.743 | 0.242 | -0.911 | 0.944 |  |
|  | pre-mating in gonotrophic cycle 1 / post-mating in gonotrophic cycle 2 | 0.770 | 0.233 | -0.864 | 0.955 |  |
|  | post-mating in gonotrophic cycle 1 / control in gonotrophic cycle 2 | 0.505 | 0.159 | -2.168 | 0.253 |  |
|  | post-mating in gonotrophic cycle 1 / pre-mating in gonotrophic cycle 2 | 0.689 | 0.220 | -1.166 | 0.853 |  |
|  | post-mating in gonotrophic cycle 1 / post-mating in gonotrophic cycle 2 | 0.715 | 0.210 | -1.141 | 0.864 |  |
|  | control in gonotrophic cycle 2 / pre-mating in gonotrophic cycle 2 | 1.365 | 0.451 | 0.940 | 0.936 |  |
|  | control in gonotrophic cycle 2 / post-mating in gonotrophic cycle 2 | 1.415 | 0.435 | 1.128 | 0.870 |  |
|  | pre-mating in gonotrophic cycle 2 / post-mating in gonotrophic cycle 2 | 1.037 | 0.323 | 0.116 | 1.000 |  |
| Viable offspring | |  |  |  |  |  |
|  | control in gonotrophic cycle 1 /  pre-mating in gonotrophic cycle 1 | 1.083 | 0.056 | 1.534 | 0.642 |  |
|  | control in gonotrophic cycle 1 / post-mating in gonotrophic cycle 1 | 1.105 | 0.057 | 1.935 | 0.381 |  |
|  | control in gonotrophic cycle 1 / control in gonotrophic cycle 2 | 1.040 | 0.059 | 0.696 | 0.983 |  |
|  | control in gonotrophic cycle 1 / pre-mating in gonotrophic cycle 2 | 1.157 | 0.065 | 2.583 | 0.101 |  |
|  | control in gonotrophic cycle 1 / post-mating in gonotrophic cycle 2 | 1.175 | 0.068 | 2.776 | 0.061 | **^⋅^** |
|  | pre-mating in gonotrophic cycle 1 / post-mating in gonotrophic cycle 1 | 1.021 | 0.053 | 0.398 | 0.999 |  |
|  | pre-mating in gonotrophic cycle 1 / control in gonotrophic cycle 2 | 0.961 | 0.054 | -0.717 | 0.980 |  |
|  | pre-mating in gonotrophic cycle 1 / pre-mating in gonotrophic cycle 2 | 1.068 | 0.060 | 1.178 | 0.848 |  |
|  | pre-mating in gonotrophic cycle 1 / post-mating in gonotrophic cycle 2 | 1.085 | 0.063 | 1.416 | 0.717 |  |
|  | post-mating in gonotrophic cycle 1 / control in gonotrophic cycle 2 | 0.941 | 0.053 | -1.084 | 0.888 |  |
|  | post-mating in gonotrophic cycle 1 / pre-mating in gonotrophic cycle 2 | 1.047 | 0.059 | 0.816 | 0.965 |  |
|  | post-mating in gonotrophic cycle 1 / post-mating in gonotrophic cycle 2 | 1.064 | 0.061 | 1.066 | 0.895 |  |
|  | control in gonotrophic cycle 2 / pre-mating in gonotrophic cycle 2 | 1.112 | 0.067 | 1.761 | 0.491 |  |
|  | control in gonotrophic cycle 2 / post-mating in gonotrophic cycle 2 | 1.130 | 0.070 | 1.970 | 0.360 |  |
|  | pre-mating in gonotrophic cycle 2 / post-mating in gonotrophic cycle 2 | 1.016 | 0.063 | 0.257 | 1.000 |  |

Significant *p*-values are indicated with asterisks (^***^ *p* < 0.001, ^**^ *p* < 0.01, ^*^ *p* < 0.05, **^⋅^** *p* < 0.1)

5. **Transgenerational effects of heat shock exposure in *Ae. aegypti* female fertility** (Table S9– S10)

Table S9. Analysis of deviance (Type II Wald Chi-square tests) of reproductive traits evaluating the transgenerational effects of heat shock exposure in *Ae. aegypti* fertility

| Trait | Factor | | df | | χ^2^ | | *p*-value | |  |
| --- | --- | --- | --- | --- | --- | --- | --- | --- | --- |
| Egg laying success | |  | |  | |  | |  | |
|  | maternal exposure | | 1 | | 0.791 | | 0.374 | |  |
|  | offspring temperature | | 2 | | 8.945 | | 0.011 | | ^*^ |
|  | interaction | | 2 | | 0.197 | | 0.906 | |  |
| Fecundity | |  | |  | |  | |  | |
|  | maternal exposure | | 1 | | 11.795 | | < 0.001 | | ^***^ |
|  | offspring temperature | | 2 | | 10.204 | | 0.006 | | ^**^ |
|  | interaction | | 2 | | 4.672 | | 0.097 | | **^⋅^** |
| Egg hatchability | |  | |  | |  | |  | |
|  | maternal exposure | | 1 | | 2.372 | | 0.124 | |  |
|  | offspring temperature | | 2 | | 2.549 | | 0.280 | |  |
|  | interaction | | 2 | | 4.708 | | 0.095 | | **^⋅^** |
| Viable offspring | |  | |  | |  | |  | |
|  | maternal exposure | | 1 | | 8.846 | | 0.003 | | ^**^ |
|  | offspring temperature | | 2 | | 6.637 | | 0.036 | | ^*^ |
|  | interaction | | 2 | | 4.041 | | 0.133 | |  |

Significant *p*-values are indicated with asterisks (^***^ *p* < 0.001, ^**^ *p* < 0.01, ^*^ *p* < 0.05, **^⋅^** *p* < 0.1)

Table S10. Full pairwise comparisons for all reproductive traits evaluating the transgenerational effects of heat shock exposure in *Ae. aegypti* fertility

| Trait | Comparison (maternal heat-exposure vs. offspring temperature / °C) | Ratio / OR | SE | Z-ratio | *p*-value |  |
| --- | --- | --- | --- | --- | --- | --- |
| Egg laying success | |  |  |  |  |  |
|  | treated mothers’ offspring at 26 °C / untreated mothers' offspring at 26 °C | 1.00 | 2.04 | 0.000 | 1.000 |  |
|  | treated mothers’ offspring at 26 °C / treated mothers’ offspring at 42 °C | 1.00 | 2.04 | 0.000 | 1.000 |  |
|  | treated mothers’ offspring at 26 °C / untreated mothers' offspring at 42 °C | 1.00 | 2.04 | 0.000 | 1.000 |  |
|  | treated mothers’ offspring at 26 °C / treated mothers’ offspring at 43 °C | 7.65 | 11.90 | 1.310 | 0.780 |  |
|  | treated mothers’ offspring at 26 °C / untreated mothers' offspring at 43 °C | 15.64 | 23.60 | 1.822 | 0.452 |  |
|  | untreated mothers' offspring at 26 °C / treated mothers’ offspring at 42 °C | 1.00 | 2.04 | 0.000 | 1.000 |  |
|  | untreated mothers' offspring at 26 °C / untreated mothers' offspring at 42 °C | 1.00 | 2.04 | 0.000 | 1.000 |  |
|  | untreated mothers' offspring at 26 °C / treated mothers’ offspring at 43 °C | 7.65 | 11.90 | 1.310 | 0.780 |  |
|  | untreated mothers' offspring at 26 °C / untreated mothers' offspring at 43 °C | 15.64 | 23.60 | 1.822 | 0.452 |  |
|  | treated mothers’ offspring at 42 °C / untreated mothers' offspring at 42 °C | 1.00 | 2.04 | 0.000 | 1.000 |  |
|  | treated mothers’ offspring at 42 °C / treated mothers’ offspring at 43 °C | 7.65 | 11.90 | 1.310 | 0.780 |  |
|  | treated mothers’ offspring at 42 °C / untreated mothers' offspring at 43 °C | 15.64 | 23.60 | 1.822 | 0.452 |  |
|  | untreated mothers' offspring at 42 °C / treated mothers’ offspring at 43 °C | 7.65 | 11.90 | 1.310 | 0.780 |  |
|  | untreated mothers' offspring at 42 °C / untreated mothers' offspring at 43 °C | 15.64 | 23.60 | 1.822 | 0.452 |  |
|  | treated mothers' offspring at 43 °C / untreated mothers' offspring at 43 °C | 2.05 | 1.47 | 0.994 | 0.920 |  |
| Fecundity | |  |  |  |  |  |
|  | treated mothers' offspring at 26 °C / untreated mothers' offspring at 26 °C | 1.039 | 0.049 | 0.807 | 0.966 |  |
|  | treated mothers' offspring at 26 °C / treated mothers' offspring at 42 °C | 0.948 | 0.041 | -1.244 | 0.815 |  |
|  | treated mothers' offspring at 26 °C / untreated mothers' offspring at 42 °C | 1.113 | 0.054 | 2.197 | 0.239 |  |
|  | treated mothers' offspring at 26 °C / treated mothers' offspring at 43 °C | 1.080 | 0.050 | 1.672 | 0.551 |  |
|  | treated mothers' offspring at 26 °C / untreated mothers' offspring at 43 °C | 1.132 | 0.067 | 2.083 | 0.296 |  |
|  | untreated mothers' offspring at 26 °C / treated mothers' offspring at 42 °C | 0.912 | 0.037 | -2.298 | 0.195 |  |
|  | untreated mothers' offspring at 26 °C / untreated mothers' offspring at 42 °C | 1.071 | 0.049 | 1.489 | 0.672 |  |
|  | untreated mothers' offspring at 26 °C / treated mothers' offspring at 43 °C | 1.039 | 0.045 | 0.893 | 0.948 |  |
|  | untreated mothers' offspring at 26 °C / untreated mothers' offspring at 43 °C | 1.089 | 0.062 | 1.490 | 0.671 |  |
|  | treated mothers' offspring at 42 °C / untreated mothers' offspring at 42 °C | 1.174 | 0.048 | 3.889 | 0.001 | ^**^ |
|  | treated mothers' offspring at 42 °C / treated mothers' offspring at 43 °C | 1.139 | 0.044 | 3.416 | 0.008 | ^**^ |
|  | treated mothers' offspring at 42 °C / untreated mothers' offspring at 43 °C | 1.194 | 0.064 | 3.313 | 0.012 | ^*^ |
|  | untreated mothers' offspring at 42 °C / treated mothers' offspring at 43 °C | 0.971 | 0.043 | -0.671 | 0.985 |  |
|  | untreated mothers' offspring at 42 °C / untreated mothers' offspring at 43 °C | 1.017 | 0.059 | 0.289 | 1.000 |  |
|  | treated mothers' offspring at 43 °C / untreated mothers' offspring at 43 °C | 1.048 | 0.059 | 0.832 | 0.962 |  |
| Egg hatchability | |  |  |  |  |  |
|  | treated mothers' offspring at 26 °C / untreated mothers' offspring at 26 °C | 0.395 | 0.148 | -2.487 | 0.128 |  |
|  | treated mothers' offspring at 26 °C / treated mothers' offspring at 42 °C | 0.993 | 0.399 | -0.018 | 1.000 |  |
|  | treated mothers' offspring at 26 °C / untreated mothers' offspring at 42 °C | 0.779 | 0.327 | -0.595 | 0.991 |  |
|  | treated mothers' offspring at 26 °C / treated mothers' offspring at 43 °C | 0.605 | 0.280 | -1.084 | 0.888 |  |
|  | treated mothers' offspring at 26 °C / untreated mothers' offspring at 43 °C | 0.838 | 0.348 | -0.426 | 0.998 |  |
|  | untreated mothers' offspring at 26 °C / treated mothers' offspring at 42 °C | 2.516 | 0.868 | 2.673 | 0.081 | **^⋅^** |
|  | untreated mothers' offspring at 26 °C / untreated mothers' offspring at 42 °C | 1.975 | 0.722 | 1.862 | 0.426 |  |
|  | untreated mothers' offspring at 26 °C / treated mothers' offspring at 43 °C | 1.534 | 0.637 | 1.031 | 0.908 |  |
|  | untreated mothers' offspring at 26 °C / untreated mothers' offspring at 43 °C | 2.123 | 0.765 | 2.090 | 0.293 |  |
|  | treated mothers' offspring at 42 °C / untreated mothers' offspring at 42 °C | 0.785 | 0.309 | -0.615 | 0.990 |  |
|  | treated mothers' offspring at 42 °C / treated mothers' offspring at 43 °C | 0.610 | 0.268 | -1.124 | 0.872 |  |
|  | treated mothers' offspring at 42 °C / untreated mothers' offspring at 43 °C | 0.844 | 0.328 | -0.436 | 0.998 |  |
|  | untreated mothers' offspring at 42 °C / treated mothers' offspring at 43 °C | 0.777 | 0.355 | -0.553 | 0.994 |  |
|  | untreated mothers' offspring at 42 °C / untreated mothers' offspring at 43 °C | 1.075 | 0.438 | 0.178 | 1.000 |  |
|  | treated mothers' offspring at 43 °C / untreated mothers' offspring at 43 °C | 1.384 | 0.626 | 0.719 | 0.980 |  |
| Viable offspring | |  |  |  |  |  |
|  | treated mothers' offspring at 26 °C / untreated mothers' offspring at 26 °C | 1.039 | 0.049 | 0.810 | 0.966 |  |
|  | treated mothers' offspring at 26 °C / treated mothers' offspring at 42 °C | 0.948 | 0.045 | -1.136 | 0.866 |  |
|  | treated mothers' offspring at 26 °C / untreated mothers' offspring at 42 °C | 1.113 | 0.053 | 2.242 | 0.218 |  |
|  | treated mothers' offspring at 26 °C / treated mothers' offspring at 43 °C | 1.080 | 0.053 | 1.584 | 0.609 |  |
|  | treated mothers' offspring at 26 °C / untreated mothers' offspring at 43 °C | 1.132 | 0.057 | 2.465 | 0.135 |  |
|  | untreated mothers' offspring at 26 °C / treated mothers' offspring at 42 °C | 0.912 | 0.043 | -1.947 | 0.374 |  |
|  | untreated mothers' offspring at 26 °C / untreated mothers' offspring at 42 °C | 1.071 | 0.051 | 1.432 | 0.707 |  |
|  | untreated mothers' offspring at 26 °C / treated mothers' offspring at 43 °C | 1.039 | 0.051 | 0.792 | 0.969 |  |
|  | untreated mothers' offspring at 26 °C / untreated mothers' offspring at 43 °C | 1.089 | 0.055 | 1.695 | 0.535 |  |
|  | treated mothers' offspring at 42 °C / untreated mothers' offspring at 42 °C | 1.174 | 0.056 | 3.378 | 0.010 | ^**^ |
|  | treated mothers' offspring at 42 °C / treated mothers' offspring at 43 °C | 1.139 | 0.055 | 2.693 | 0.077 | **^⋅^** |
|  | treated mothers' offspring at 42 °C / untreated mothers' offspring at 43 °C | 1.194 | 0.060 | 3.544 | 0.005 | ^**^ |
|  | untreated mothers' offspring at 42 °C / treated mothers' offspring at 43 °C | 0.971 | 0.048 | -0.608 | 0.991 |  |
|  | untreated mothers' offspring at 42 °C / untreated mothers' offspring at 43 °C | 1.017 | 0.051 | 0.333 | 1.000 |  |
|  | treated mothers' offspring at 43 °C / untreated mothers' offspring at 43 °C | 1.048 | 0.054 | 0.905 | 0.945 |  |

Significant *p*-values are indicated with asterisks (^***^ *p* < 0.001, ^**^ *p* < 0.01, ^*^ *p* < 0.05, **^⋅^** *p* < 0.1)
